# Peroxiredoxinylation buffers the redox state of the proteome upon cellular stress

**DOI:** 10.1101/2023.12.13.571451

**Authors:** Gerhard Seisenbacher, Zrinka Raguz Nakic, Eva Borràs, Eduard Sabidó, Uwe Sauer, Eulalia de Nadal, Francesc Posas

## Abstract

The redox state of proteins is essential for their function and guarantees cell fitness. Peroxiredoxins protect cells against oxidative stress, maintain redox homeostasis, act as chaperones and transmit hydrogen peroxide signals to redox regulators. Despite the profound structural and functional knowledge of peroxiredoxins action, information on how the different functions are concerted is still scare. Using global proteomic analyses, we show here that the yeast peroxiredoxin Tsa1 binds hundreds of proteins of essential biological processes, including protein turnover and carbohydrate metabolism. Several of these interactions are of covalent nature and failure of this peroxiredoxinylation leads to global changes in the metabolome and reduced stress resistance. Thioredoxins directly remove TSA1-formed mixed disulfide intermediates, thus expanding the role of the thioredoxin-peroxiredoxin redox cycle pair to buffer the redox state of proteins in an unprecedented way.

## Introduction

The redox modifications of proteins constantly changes by internal and external cues and its regulation is essential for protein function and to guarantee cell fitness^1^. The small antioxidant proteins Peroxiredoxins, which are found in all domains of life, arose as central players in the cellular antioxidant response. Although they have moderate peroxidase activity, their abundance enables them to detoxify up to 90% of cytosolic H_2_O_2_^2^. Peroxiredoxins are essential in redox processes ranging from oxidative stress protection to redox signaling^3^. Disruption of peroxiredoxins results in decreased cell fitness, and alteration of their expression and function is associated with oxidative stress-related conditions such as aging, inflammatory and neurological diseases, and cancer^4^. 2-Cys peroxiredoxins (2-Cys Prxs) are ubiquitous detoxifiers of H_2_O_2_ that rely on two key residues: the peroxidatic cysteine (C_P_) which reacts with peroxides and a C-terminal resolving cysteine (C_R_) that reacts with the sulfenic oxidized C_P_ and drives the catalytic redox cycle^5^.

Originally, the oxidative modification of proteins was considered primarily as damaging^6^. Nowadays the physiological role of H_2_O_2_ as a second messenger is well established^7,8^. Thiol oxidations have been added to a big list of cysteine post-translational modifications such as glutathionylation, sulfhydrylation, nitrosation, and disulfide bonding^9^. Many of these modifications serve-possibly mutually exclusive or competing-for redox regulation. For instance, the ATPase alpha subunit has a redox-modified cysteine (C294) that directly affects cellular ATP production^10^. Glutathionylation of the conserved Cys374 of actin enhances actin depolymerization and plays a role in stress fiber formation^11^. Transient glutathionylation of glycerol aldehyde-3-phosphate dehydrogenase protects its enzymatic site from irreversible oxidation during high oxidative stress^12,13^.

A milestone in understanding how hydrogen peroxide function as a second messenger was the functional dissection of the molecular environment of the peroxidatic cysteine that is critical for the reactivity towards H_2_O_2_. While the key residues in the PXXX(S/T)XXC_P_ motif are highly conserved, the molecular environment underwent changes that favor the oxidative inactivation of eukaryotic peroxiredoxin compared to bacterial ones^14^. The C_P_ hyperoxidation inactivates the peroxidase function and allows H_2_O_2_ levels-otherwise kept very low-to rise and act as second messengers for redox-sensitive signaling factors^7,8,15,16^. This inactivation shunt leads to a shift from peroxidase to signaling function of peroxiredoxins^17^. Another consequence of peroxiredoxin hyperoxidation of the peroxidatic cysteine to the sulfonic acid form is the formation of higher order structures with chaperone function^18^. In contrast to the 2-Cys peroxidatic cysteines the majority of cysteine thiol group found in proteins have moderate reactivity towards low, physiological H_2_O_2_ concentrations. This conundrum has been reconciled with the findings that peroxiredoxins function as mediators of redox signaling^19–21^. The oxidized peroxidatic cysteine forms transient mixed disulfide intermediates (MDIs) with a target protein thiols, which reacts with a structural vicinal cysteine in the target protein leading to a disulfide bridge modified protein and thereby transmitting the cellular redox state to regulatory proteins such as Stat3 or Bcy1^19,22^. Recently it has been shown that association of the peroxiredoxin Tpx1 with Sty1 leads to the formation of a signaling complex that activates SAPK signaling independent of the canonical activation pathway in *S. pombe*^23^. Thus, redox sensitive peroxiredoxin - protein interactions are potentially widespread and novel modes of action are most likely to be discovered upon identification of new targets. In this line, large-scale proteomics experiments in eukaryotic organisms revealed extended formation of MDIs by peroxiredoxins^24,25^. Yet, how PRXs select their targets or how the formation of adventitious mixed disulfides is prevented is currently far from being understood^21,24,26^.

The highly expressed peroxiredoxin *TSA1* from the yeast *Saccharomyces cerevisiae* has been pivotal in the structural and functional understanding of eukaryotic 2-Cys PRX^20,27–29^. Here, we show, using global proteomic analyses that Tsa1 physically interact with hundreds of proteins of central biological processes, such as proteins involved in metabolism and translation. Interestingly, the majority of those interactions is redox sensitive and several are covalent by the formation of disulfide bridges involving the peroxidatic cysteine and cysteines in the target protein changing its activity. Remarkably, this binding is independent of its hyperoxidized chaperone function and independent of the recently described mechanism during oxidative aging involving sulfiredoxin and the Hsp70/Hsp104 heat shock protein complex^30^. Instead, we find that the covalent interaction of peroxiredoxins with its interactome is induced by various stresses and is regulated by the thioredoxin system. Thus, indicating a role for the association of Tsa1 in buffering the redox state of the proteome upon cellular stress.

## Results

### Tsa1 forms wide-spread mixed disulfide intermediates

Mixed disulfide intermediates (MDI) of peroxiredoxins and target proteins are the base of redox relays widely found in eukaryotes^25,31,32^. Similar to what has been proposed in mammalian cells we suspected that important functions of the major yeast peroxiredoxin Tsa1 could be mediated by MDIs^21,24^. To this aim we tested wild-type, the C48S (peroxidatic cysteine) and the C171S (resolving cysteine) mutant of Tsa1 for their ability to form MDIs and distinguish the importance of the respective cysteines in their formation (Fig.1a). All strains were tagged C-terminally by a single HA epitope in their endogenous locus which does not affect Tsa1 expression nor function (Supplementary Fig. 1a-c). Yeast whole cell extracts were prepared under either non-reducing conditions, in the presence of 2mM S-methyl methanethiosulfonate (MMTS) as free thiol blocker, or reducing conditions (2mM DTT). We found that HA-tagged and untagged wild-type Tsa1 forms - next to partially and fully disulfide linked dimers - MDIs in a range of 50 to 200 kDa to which we refer herein as <u>T</u>sa1-Induced <u>M</u>ixed <u>D</u>isulfide Intermediates (TIMDIs) (Fig. 1b and Supplementary Fig. 1d,e). TIMDIs are strongly enhanced in the resolving cysteine (C171S) mutant and absent in the peroxidatic cysteine mutant (C48S) (Fig. 1b and Supplementary Fig. 1d). The denaturing base of SDS-PAGE confirms the covalent nature of these adducts and their absence in reducing condition indicates the redox-sensitivity of the potential Tsa1-target protein attachment. Interestingly, while both, the peroxidatic and resolving mutant display in the redox cycle of peroxiredoxins we show that in TIMDIs formation, they have completely opposing phenotypes. To proof that the high molecular weight TIMDIs contain Tsa1 we performed a two dimensional SDS-PAGE outlined in Supplementary Fig. 1f. The collapse of the high-molecular weight Tsa1^C171S^-HA TIMDIs to monomeric Tsa1 in reducing conditions confirmed the association of Tsa1 with target proteins of various molecular weights (Fig. 1c).

**Fig. 1.**
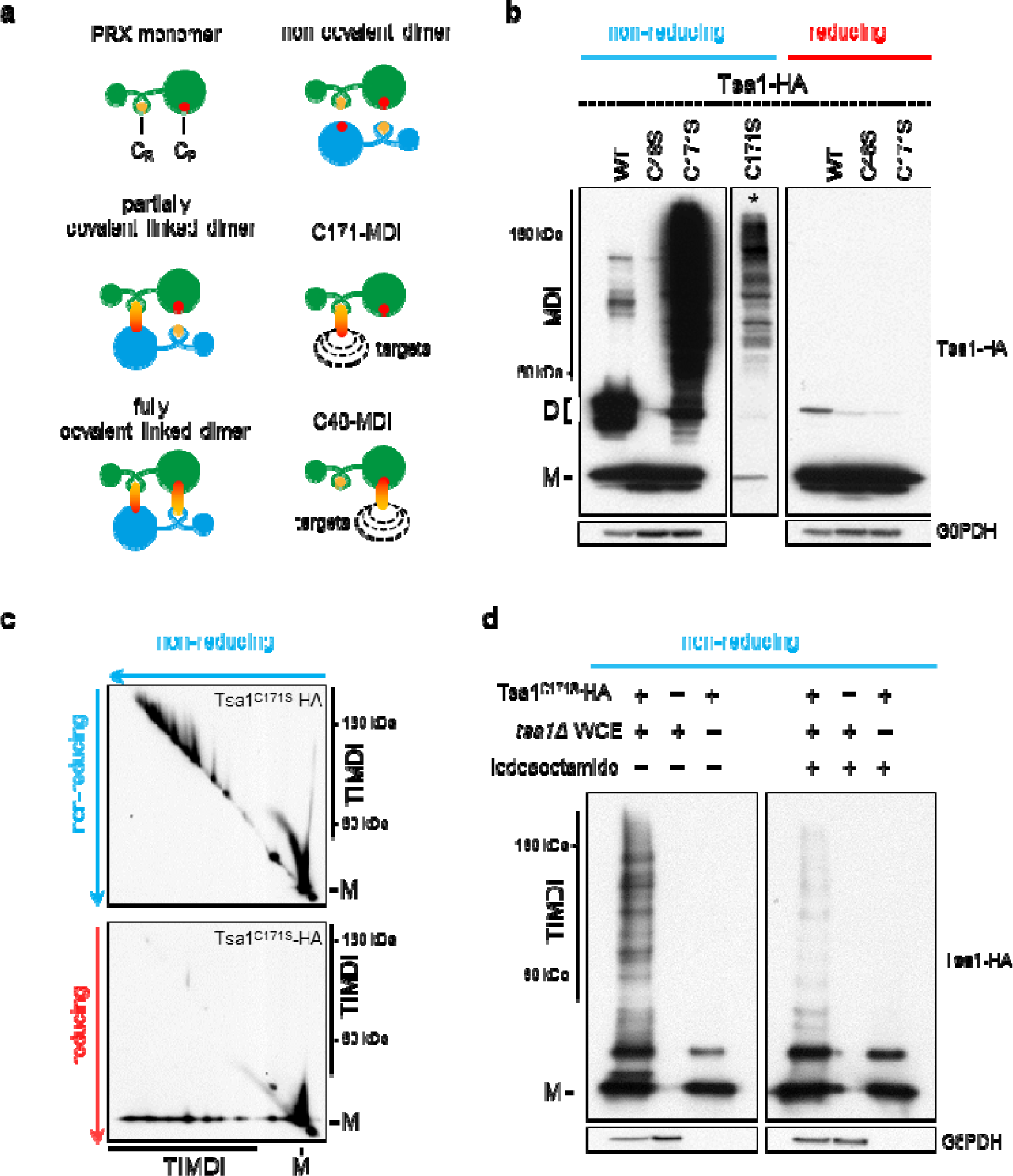
Tsa1 forms widespread covalent association relying on its peroxidatic cysteine. **a,** Schematic display of covalent cysteine dependent interaction of 2-Cys peroxiredoxins. Peroxiredoxins are displayed in green and blue. The peroxidatic cysteine (C_P_) is shown as filled red circle, the resolving cysteine (C_R_) is shown as filled orange circle and disulfide bridges as fused circles. Potential targets of mixed disulfide intermediates are indicated as dashed black circles. Experimental evidence for the different dimeric forms can be found in Supplementary Fig. 1e. **b,** Immunoblotting of whole cell extracts prepared in non-reducing and reducing conditions of TSA1^WT^-HA, the peroxidatic cysteine mutant TSA1^C48S^-HA and the resolving cysteine mutant TSA1^C171S^-HA to detect mixed disulfide intermediates. Lower exposure (*) of the Tsa1^C171S^-HA lane shows the complex high molecular weight pattern. Shorter exposure blots and experiments with untagged Tsa1 can be found in Supplementary Fig. 1d. **c,** High molecular weight adducts incorporate Tsa1. Two dimensional SDS-PAGE and immunoblotting for HA confirm the redox-sensitive presence of Tsa1^C171S^-HA in the high molecular weight adducts. Schematic outline of the 2D-SDS PAGE is found in Supplementary Fig. 1f. **d,** *in vitro* Tsa1-Induced Mixed Disulfide Intermediate (TIMDI) formation. Whole cell extracts from *tsa1*Δ cells were prepared and incubate with or without the thiol blocking agent iodoacetamide. After dialysis to remove the blocking agents equal amounts of non-blocked/blocked whole cell extracts was incubated with recombinant Tsa1^C171S^-HA. **b,c,d,** *M* refers to the Tsa1 monomer, *D* indicates the fully and partially disulfide linked Tsa1 homodimer. *MDI/TIMDI* indicates the mixed disulfide intermediates.

To recapitulate the *in vivo* results, we established an *in vitro* system and monitored direct binding of recombinant Tsa1 to proteins from a *tsa1*Δ yeast extract. Recombinant wild-type Tsa1 forms rapidly disulfide-linked homodimers in the absence of a reducing system, thus we used Tsa1^C171S^-HA which cannot form homo-dimeric C_P_-C_R_ linked disulfides. In the assay, recombinant Tsa1^C171S^-HA was mixed with whole cell extract from *tsa1*Δ cells in the presence of the reducing agent DTT. Dialyzing DTT from the reaction led to the formation of TIMDIs *in vitro*. Correspondingly, the blocking of free cysteines in whole cell extracts with iodoacetamide strongly reduces TIMDI formation (Fig. 1d). Thus, Tsa1 has the potential to covalently attached to many proteins, by the formation of disulfide bridges via its peroxidatic cysteine and presumably cysteine on its targets. Therefore, redox signaling relays show strong evolutionary conservation and could be a more widespread mechanism than expected^21^. Intriguingly, mixed disulfide intermediates with peroxiredoxins in redox signaling relays are of transient nature due to rapid intramolecular disulfide exchange by a structurally close cysteine, releasing the reduced peroxiredoxin and leaving the oxidized target protein behind. This type of disulfide exchange would also apply to C171S-MDIs and one would therefore expect an increase in oxidized target proteins instead of PRX-target MDIs. Thus, the extent of TIMDIs in the Tsa1^C171S^-HA mutant challenge this view and qualifies the covalent attachment of peroxiredoxin via its peroxidatic cysteine to target proteins as a potential post-translational modification of cysteine such as gluthathionylation or oxidations^9^.

### Tsa1-induced Mixed Disulfide Intermediates are induced upon H_2_O_2_ stress

In contrast to the C171S mutant, TIMDI formation of wild-type Tsa1 is low under standard growth conditions. In mammals 2-Cys peroxiredoxins have been established as sensitive and abundant forwarders of H_2_O_2_-derived oxidizing equivalents, and thus as highly efficient enablers of protein thiol oxidation and redox signaling^21,24^. To this end we assessed whether H_2_O_2_ stimulation *in vivo* induced TIMDIs formation. We treated cultures of Tsa1^WT^-HA with H_2_O_2_ from zero to 1 mM for 10 minutes. Those parameters were chosen since within 40 minutes more than 50% of H_2_O_2_ is scavenged in rich media cultures (referring to a 0.2 mM starting concentration), within 10 minutes newly synthesized antioxidants e.g. catalases do not matter yet and concentrations higher than 1 mM can be considered severe oxidative stress^33–35^. A slight TIMDI increase is observed at a physiological sub-lethal H_2_O_2_ concentrations (0.031 and 0.062mM). A significant burst of TIMDIs was observed when H_2_O_2_ increased to levels considered permissive for redox signaling (0.125 and 0.25mM), but not sufficient to cause substantial Tsa1 peroxidatic cysteine hyperoxidation (C48-SO_2/3_) nor lethality in the wild type (Fig. 2a,b,c and Supplementary Fig. 2a)^36^. Once H_2_O_2_ reached levels that lead to irreversible oxidation of the peroxidatic cysteine, and the wild type shows affected growth, TIMDIs became strongly impaired confirming the requirement of the peroxidatic cysteine to get engaged in mixed disulfide formation with targets (Figs. 2a,b,c and Supplementary Fig. 2a,b). Likewise, strains with untagged Tsa1 show a comparable TIMDI and peroxidatic hyperoxidation dynamics (Supplementary Fig. 1c and 2c).

**Fig. 2.**
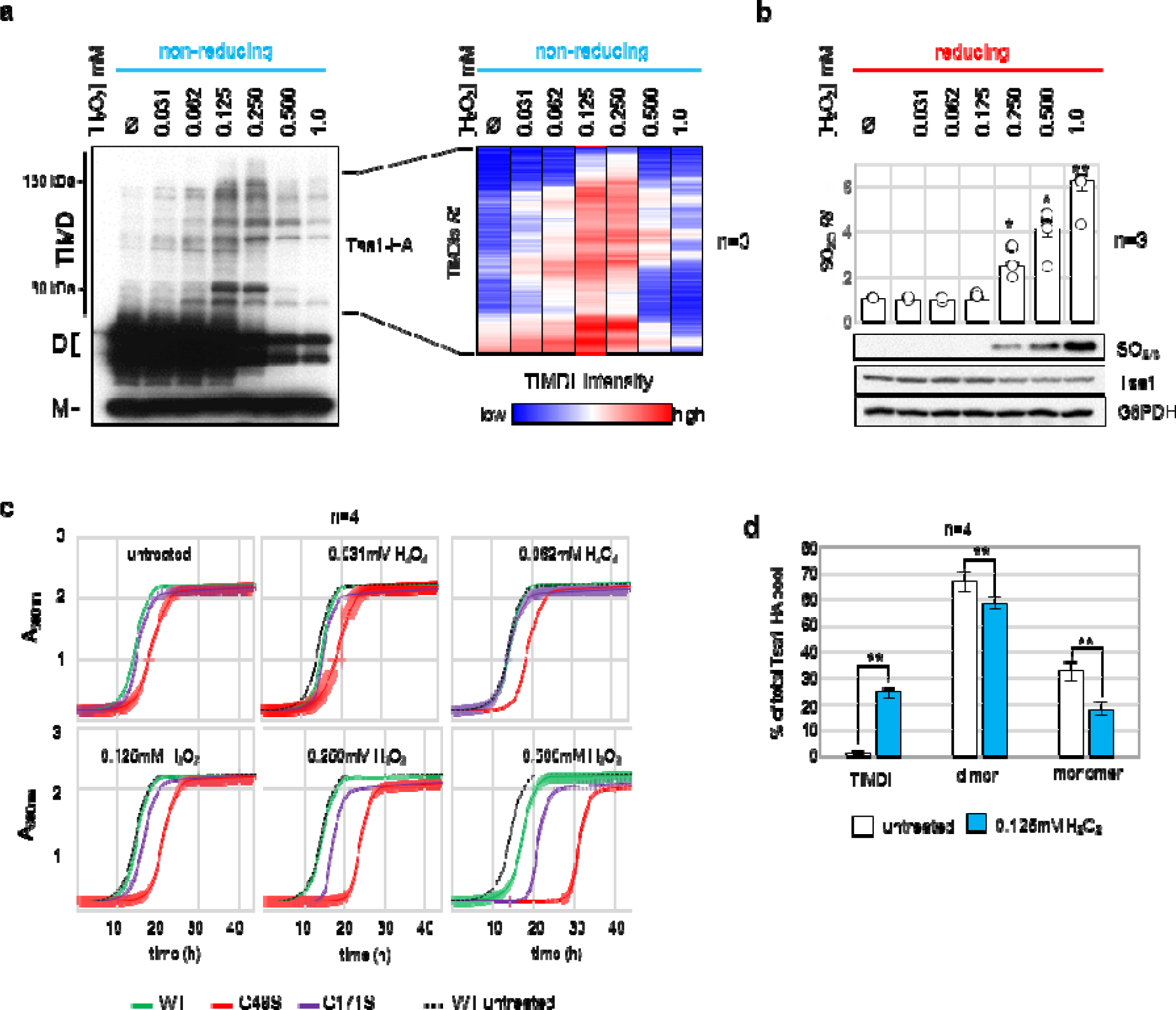
Tsa1-induced mixed disulfide increases in a dose response to H_2_O_2_. **a,** Hydrogen peroxide dose response of TIMDI induction. Tsa1^WT^-HA cultures were treated for 10’ with the indicated H_2_O_2_ concentration and non-reducing whole cell extracts were separated by a dual 10%15% SDS-PAGE. Western blot membranes were probed with anti-HA (Tsa1). Lower exposure of the blot and change in the Tsa1 monomer and homodimer ratio can be found in Supplementary Fig. 2a. Heat map of relative TIMDI intensity values of the indicated region was generated from biological triplicates. Relative TIMDI intensities (*RI*) are shown with lowest in blue and highest intensity in red. Log_2_-fold change and statistical analysis can be found in Supplementary Fig. 2b. **b,** Cysteine sulfinylation/sulfonylation (SO_2/3_) was assessed on same H_2_O_2_ dose response as in Fig. 2a. Relative SO_2/3_ intensities (*RI*) were quantified from biological triplicates, median values ± SD and single data points (circles) are shown. Two tailed Students test were performed, * = p-value < 0,05, ** = p-value < 0,005. Representative western blot membranes probed for anti-SO_2/3_, anti-HA (Tsa1) and anti-G6PDH (loading control) are shown. **c,** Growth curves of cells carrying Tsa1^WT^-HA (green), Tsa1^C48S^-HA (red) and Tsa1^C171S^-HA (purple) in unstressed conditions and at the indicated H_2_O_2_ concentrations. A_600_ of liquid cultures was measured every 10’ to follow cellular growth. The unstressed Tsa1^WT^-HA curve (dashed black line) depicted in each graph is meant for comparison to the H_2_O_2_ response. Experiment was performed as n= 4, median values ± SD is shown. **e,** The percentage of TIMDIs, monomer and dimer of the Tsa1^WT^-HA pool in unstressed and H_2_O_2_ treated conditions was determined by western blot quantification in triplicates. Two tailed Students test were performed, ** = p-value < 0,005. Linear range for quantification was measured in pixel intensities and graph can be found in Supplementary Fig. 2e.

Additionally, to exclude that TIMDI induction is an artificial effect, various thiol-blocking agents were compared and similar results were obtained (Supplementary Fig.2d). For a quantitative estimation of the TIMDI extend we measured the amount of TIMDIs, Tsa1 homo-dimer and monomer analyzing the plot profile of Tsa1^WT^-HA lanes of western blot analysis in a linear range (Supplementary Fig. 2e). Under normal conditions, TIMDIs are barely detectible and upon low H_2_O_2_ stress TIMDI significantly increased to more than 20% of the Tsa1 pool (Fig. 2d). Since Tsa1 is one of the highest expressed proteins and TIMDI formation is a dynamical process the extent of TIMDI formation upon oxidative stress is to be expected to be of physiological relevance.

### TIMDIs protect against protein damage upon stress

Exogenous stimulation with H_2_O_2_ affects directly the oxidation state of Tsa1 and induces redox relays^21^. The extent of potential targets suggested by the C171S trap mutant led us to speculate that the observed TIMDIs could regulate processes beyond H_2_O_2_ signaling relays. Tsa1 act as a ribosome-associated chaperone to protect newly synthesized proteins and prevent protein aggregation upon oxidative stress^20,27,37,38^. Therefore we first tested if TIMDI formation upon H_2_O_2_ could be a result of errors in *de novo* protein synthesis. Cycloheximide-induced block of translation did not prevent TIMDI formation in cells treated with 0.125mM H_2_O_2_ (Fig. 3a). Incorporation of the amino acid analogue AZC (L-azetidine-2-carboxylic acid) leads to the accumulation of translationally misfolded proteins^39^. We followed cytoplasmic foci formation of endogenously GFP-tagged Tsa1 and the disaggregase Hsp104, indicative of accumulation of translationally aggregated proteins. Addition of 5 mM AZC to the medium lead to the formation of foci, which were prevented by the addition of cycloheximide (Fig. 3b and Supplementary Fig.3a). No induction of TIMDIs by AZC was observed (Fig. 3c). Thus the formation of TIMDIs takes place at moderate H_2_O_2_ levels that do not saturate the peroxiredoxin system and does not rely on *de novo* protein production nor formation of the hyperoxidized chaperone.

**Fig. 3.**
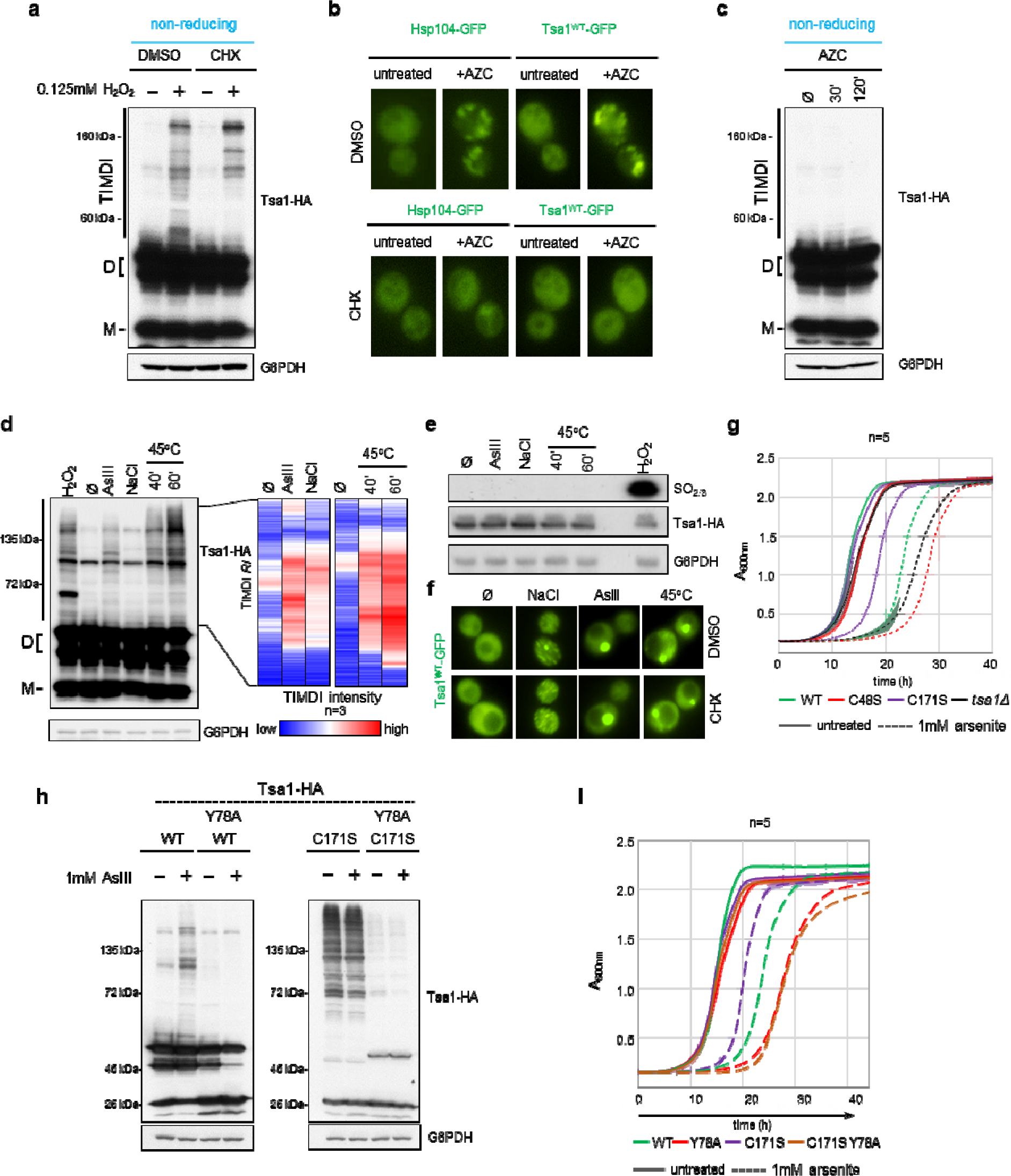
TIMDI is induced upon environmental stress, protects against arsenite toxicity and is independent of *de novo* protein production. **a,** *de novo* protein production does not contribute to TIMDI formation. Cells were pretreated with 100 μg/ml cyclohexamide for 15’ followed by ± 1 mM H_2_O_2_ for 10’. Immunoblots were probed with anti-HA (Tsa1) and anti-G6PDH (loading control). **b,** Aggregation of *de novo* proteins upon proline analog L-azetidine-2-carboxylic acid treatment (AZC). Cycloheximide-sensitive foci formation of Hsp104-GFP and Tsa1-GFP were assessed by microscopy after 60’ 5mM AZC treatment. See Supplementary Fig. 3a for untreated controls. **c,** Protein aggregation does not induce TIMDIs. Tsa1^WT^-HA cells were treated with 5mM AZC for the indicated time points. Non-reducing whole cell extracts were separated by dual 10%15% SDS-PAGE and immunoblots were probed with anti-HA (Tsa1) and anti-G6PDH (loading control). **d,** Various stresses induce TIMDIs. Non-reducing whole cell extracts of Tsa1^WT^-HA strains treated with the indicated stresses was separated by dual 10%15% SDS-PAGE and membranes were probed for anti-HA (Tsa1) and anti-G6PDH (loading control). Heat map of relative TIMDI intensities of Tsa1^WT^-HA in unstressed versus 10’ 1 mM AsIII, 10’ 0.8 M NaCl and 45°C heat shock for 40’ and 60’ of the indicated region was generated from biological triplicates. 10’ treatment with 0.125mM H_2_O_2_ was included as control. Relative TIMDI intensity (*RI*) are shown with lowest in blue and highest intensity in red. Log_2_-fold change and statistical analysis can be found in Supplementary Fig. 3c. See Supplementary Fig. 3b for concentration and heat shock intensity estimation for this experiment. **e,** Cysteine sulfinylation/sulfonylation of the peroxidatic cysteine (C48) of Tsa1-HA was assessed for the indicated stresses and compared to 1 mM H_2_O_2_ by western blotting. Membranes were probed with anti-SO_2/3_, anti-HA (Tsa1) and anti-G6PDH (loading control). **f,** Formation of Tsa1-GFP foci was investigated for the indicated stresses with 15’ pretreatment with DMSO or CHX by microscopy. Same experiment with Hsp104-GFP can be found in Supplementary Fig. 3d. **g,** TIMDIs protect against sodium arsenite induced stress. Growth of the indicated strains was assayed in liquid media ± 1mM sodium arsenite and A_600nm_ was measured every 10’ to follow growth. Experiment was performed as n= 5, median values ± SD is shown. **h,** Sodium arsenite induced TIMDIs depends on Tsa1 Y78. Whole cell extracts of the indicated strains and conditions were prepared in non-reducing conditions and separated by dual 10%15% SDS-PAGE. After transfer, membranes were probed with anti-HA (Tsa1) and anti-G6PDH (loading control). **i,** Cell fitness upon sodium arsenite stress depends on TIMDI induction. Growth of the indicated strains was assayed in liquid media ± 1mM sodium arsenite and A_600nm_ was measured every 10’ to follow growth. Experiment was performed as n= 5, median values ± SD is shown.

Next, we asked whether other stresses that affect protein homeostasis and the cellular redox balance lead to the formation of TIMDIs. Analogous to the H_2_O_2_ dose-response, non-reducing whole cell extracts of cells treated with various stresses (arsenite, sodium chloride and heat) were analyzed by non-reducing SDS-PAGE. Strikingly, especially arsenite and heat stress and to lesser extent high osmotic stress induced significant amounts of TIMDIs in an intensity dependent manner (concentration or temperature) (Fig.3d and Supplementary Fig.3b,c). Importantly, none of these stresses led to Tsa1 hyperoxidation, underscoring the mechanistic difference to the hyperoxidized Tsa1 chaperone (Fig. 3e). Of note, arsenite, osmotic and heat stress induced Tsa1-GFP foci that were not affected by cycloheximide induced translational block (Fig. 3f). Similar foci of Hsp104-GFP were observed upon osmotic stress that largely remained unaffected by cyclohexamide. In contrast, upon heat and arsenite stress foci formed by Hsp104-GFP appear more dispersed than those of Tsa1-GFP and were considerably affected by cycloheximide (Supplementary Fig.3d). This observation indicates that Hsp104 is not involved in TIMDI formation and supports that TIMDIs are mechanistically different from the C48-oxidation dependent Tsa1 chaperone. Furthermore, the cycloheximide sensitive phenotype raises the possibility that Hsp104 is involved in *de novo* protein protection while Tsa1 acts on mature proteins in the tested stress situations.

Arsenite is an important environmental toxic pollutant and the strong induction of TIMDIs upon arsenite stress led us to investigate this stress in more detail^40^. We found that the *tsa1*Δ deletion and the Tsa1^C48S^-HA strains showed an increased sensitivity to sodium arsenite compared to the wild type. Contrasting, the Tsa1^C171S^-HA mutant grew much better suggesting a direct role for TIMDI formation in stress adaptation (Fig. 3g). The sodium arsenite resistance phenotype directly correlates with the extent of TIMDI formation, and when we introduced an Y78A mutation, which prevents TIMDIs formation, in Tsa1^WT^-HA and Tsa1^C171S^-HA basal and arsenite stress induced TIMDIs were strongly reduced (Fig. 3h)^41^. Strikingly, the Y78A mutation led not only to a complete reversal of the Tsa1^C171S^-HA arsenite resistance, but also increased the arsenite sensitivity of the Tsa1^WT^-HA strain, while under unstressed conditions no survival deficiency was observed (Fig. 3i). Overall, peroxiredoxinylation acts independently of the classical hyperoxidized Tsa1 chaperone and stress-induced TIMDI formation is not only a hallmark of stresses that affect the cellular redox state, but contribute to stress protection.

### Identification of the redox-sensitive Tsa1 interactome in *S. cerevisiae*

Cysteine mutant interactor traps have been previously used successfully to identify novel interactors for thioredoxins^42,43^. We employed the C171S mutant as a Tsa1-modification interactor trap to decipher general TIMDI interactors independent of the need of stress induction. To gain a comprehensive understanding of the potential targets of peroxiredoxinylation, we identified and compared the redox-sensitive interactome of C171S and wild-type Tsa1 by mass spectrometry (Fig. 4a). TAP-tagged Tsa1^WT^ or Tsa1^C171S^ cultures were split in two, and proteins were extracted either in the presence of the thiol blocking agent MMTS (non-reducing) or of DTT (reducing). Tsa1 and its interactors were purified using IgG binding beads and high stringency washes were applied. After this, beads were washed several times with MMTS containing buffer to block thiol groups that were freed in the reducing conditions. Tsa1 and its interactors were released from the beads using TEV protease and subjected to a second round of immunoaffinity purification using calmodulin beads. After elution, Tsa1 and its covalent and non-covalent interactors were identified by mass spectrometry. Considering candidates that have been identified in both duplicates and applying a 5% FDR for peptides, we found 211 and 599 interactors with the Tsa1^WT^-TAP and Tsa1^C171S^-TAP mutant, respectively (Fig. 4b and Supplementary Data 1). The majority of binding partners showed a redox-sensitive behavior and an enrichment in non-reducing conditions (Fig. 4b,c). Of the Tsa1^WT^-TAP interactors 55.5% and of the Tsa1^C171S^-TAP interactors 64.3% were found exclusively in the non-reducing condition (NR-specific) supporting the idea of a redox-sensitive interactome of Tsa1. A position-frequency blot displayed no enrichment of a particular linear sequence suggesting a different mode of target selection of Tsa1 than a specific linear motif (Supplementary Fig. 4a). Neither the proximity of cysteines in the linear sequence of targets is an indicator of interaction (Supplementary Fig. 4b). In case of redox relays, for example, the proximity of cysteines in the tertiary structure could be determining. As expected for mixed disulfide mediated intermediates, interactors that are specific for non-reducing conditions showed an overrepresentation of proteins with more than five to eight cysteines (Fig. 4d and Supplementary Data 2). In the same line, only one percent of the non-reducing specific interactors had no cysteine, which is ten times less than expected from the overall proteome (Fig. 4d and Supplementary Data 2). Overall, this indicates that general Tsa1-target interaction can be independent on (particular) cysteines, but redox-sensitive and covalent interaction are favored by the presence of cysteines. Binding of HSP70 (Ssa1 or Ssa2) is an example for a known non-covalent interaction with Tsa1 and accordingly this interaction is found to be redox-insensitive in our Tsa1 interactome (Supplementary Data 1)^30^.

**Fig. 4.**
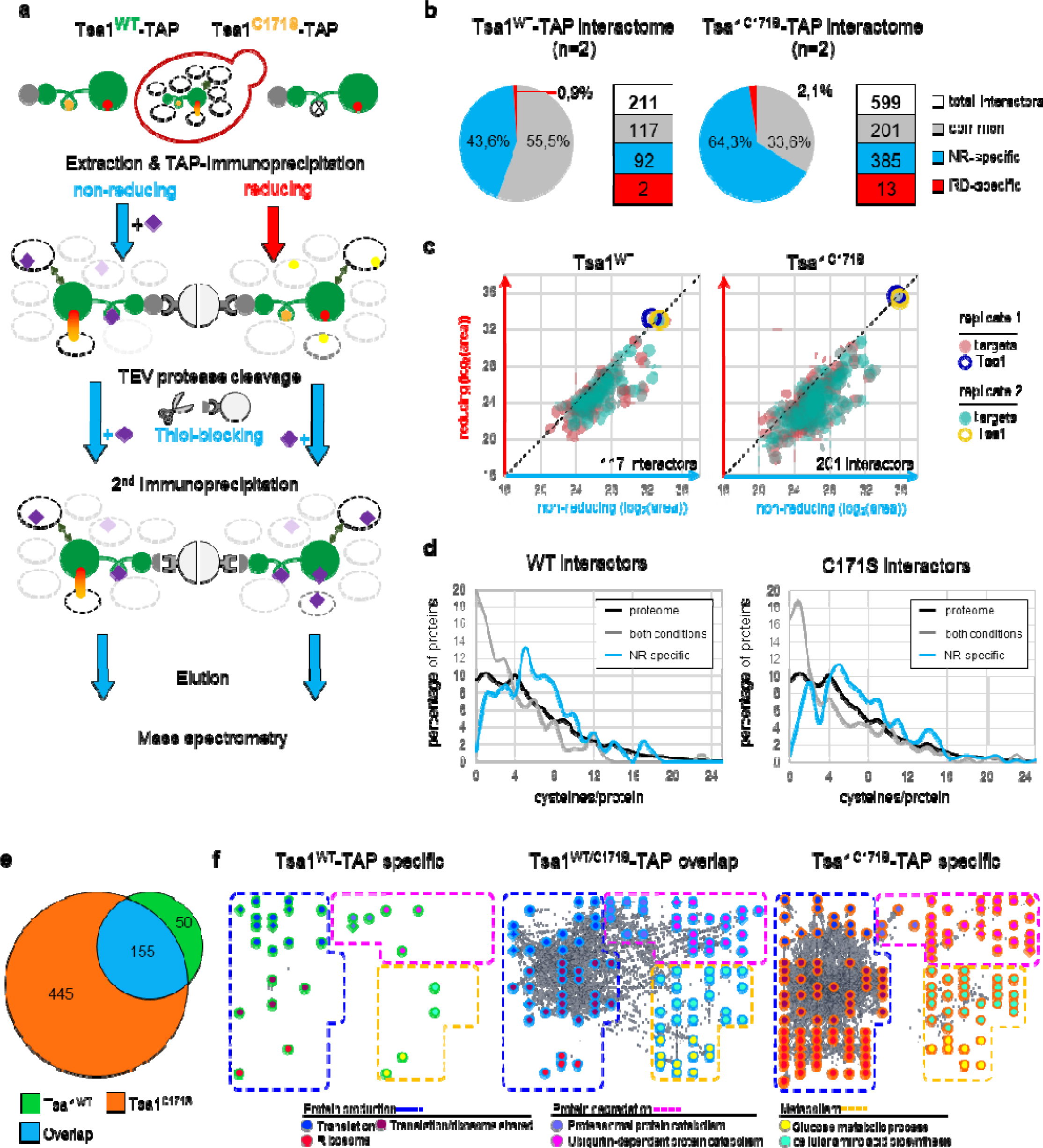
Tsa1 interacts with a large portion of the proteome in a redox-dependent manner. **a,** Schematic workflow of the mass spectrometry experiment to identify the redox-sensitive interactome of Tsa1^WT^-TAP and Tsa1^C171S^-TAP. Tsa1 and its cysteines are depicted as in Fig. 1a. Target cysteines are indicated as yellow filled circles. The C171S mutation is indicated by a circle with an *x*. The thiol-blocker MMTS is depicted as purple rhombus. **b,** Pie diagram of percentage of interactors that are found in both, non-reducing and reducing, conditions (*common*, grey), or are specific for one condition: non-reducing (*NR-specific*, blue) and reducing (*RD-specific*, red). Absolute numbers of total, common, NR-specific and RD-specific interactors are indicated in the Figure**. c,** The log_2_(area) values of Tsa1^WT^-TAP and TSA1^C171S^-TAP interactors were plotted as reducing (y-axis) against non-reducing (x-axis). Binders that were detected in both, non-reducing and reducing, conditions (*common*) are represented. The bait Tsa1 is shown as hollow circles (blue = replicate 1 and yellow = replicate 2). The interactors of the duplicates are shown as filled circles (green = replicate 1, red = replicate 2). Numbers next to the x-axis represents the numbers of interactors specifically identified in non-reducing conditions. Data are found in Supplementary Data 1. **d,** Percentage of proteins (y-axis) containing the indicated cysteine content/protein (x-axis). Interactors, which are shared in non-reducing and reducing conditions (*common*), are shown in grey, non-reducing-specific are shown in blue and the proteome wide distribution as black line. Due to the low number, reducing-specific candidates have been omitted from the analysis. Data are found in Supplementary Data 2. **e,** Overlap (blue) of interactors between the Tsa1^WT^-TAP (green) and Tsa1^C171S^-TAP (orange) interactome. Numbers of the interactors are indicated within the sections. **f,** Representative GO terms revealed by the PANTHER GO enrichment analysis. Full list of enriched GO terms with FDR and p-values is found in Supplementary Data 2.

The significant overlap of interactors (> 73%) between Tsa1^WT^ and Tsa1^C171S^ indicates that the resolving mutant indeed captures a bigger fraction of Tsa1 interactors (Fig. 4e). To discover the overrepresented biological processes of the interactors, we performed a PANTHER Overrepresentation Test^44^. Interactors in the redox-sensitive Tsa1 interactome included carbohydrate metabolism, as well as protein turnover processes such as translation, ribosomal biogenesis, chaperone function and proteolysis. Additionally, cellular metabolic processes such as amino acid biosynthesis, ATP generation as well as glucose metabolism were significantly enriched (Fig. 4f, Supplementary Fig. 4c and Supplementary Data 2). Supporting our findings, proteomics studies exploring cysteine accessibility and oxidations highlight cellular protein production and metabolism as oxidation sensitive processes^45,46^(Supplementary Fig. 4d). Proteins found in those processes generally are highly expressed and most of the interactors found belong to the 25% highest expressed proteins (Supplementary Fig. 4e). As aforementioned, the dynamic nature of TIMDIs might lead to an underestimation of Tsa1 targets. The use of the C171S mutant as an interactor trap not only confirms the majority of wild-type Tsa1 interactors, but strongly extents our target list to lesser-expressed proteins (Fig. 4e and Supplementary Fig. 4e). Importantly, while the majority of Tsa1 targets – and Tsa1 itself - are highly expressed the ability to form TIMDI adducts is not necessarily dependent on protein abundance. A 10-fold reduction of Tsa1 (by exchanging the endogenous Tsa1 promoter to a low-level expressing Rsp5 promoter) does not prevent TIMDI formation, but strongly reduces the extent of TIMDI induction upon oxidative stress (Supplementary Fig. 4f). On the other side Tdh3, a Glyceraldehyde-3-phosphate dehydrogenase, which is expressed to comparable levels as Tsa1 and also contains two cysteines in its catalytic core, does not show mixed disulfide induction upon 0.125 mM H_2_O_2_ like observed with Tsa1 (Supplementary Fig. 4g)^47^. Thus, features of 2-Cys peroxiredoxins such as their sensitivity to peroxidatic cysteine oxidation, their high steady state expression levels and their dual functions as peroxidase and chaperone, render them prone to form biological active adducts.

We demonstrated that the majority of Tsa1 interactions are redox sensitive and the C171S mutant is a valid tool to retrieve Tsa1 interactors. In a second, complementary approach, we aimed to specifically enrich covalent and peroxidatic cysteine dependent interactors by comparing the binding partners that have been DTT-eluted from a non-reducing immunopurification of Tsa1^C171S^-HA and Tsa1^C48S^-HA strains (Fig. 5a). We used iodoacetamide in the first thiol blocking step and N-ethylmaleimide in the second to differentially label accessible cysteines and those involved in mixed disulfide intermediates. Performing this experiment in biological triplicates, we found that indeed more than 2/3 of proteins were eluted specifically from Tsa1^C171S^ compared to the peroxidatic mutant (Fig.5b and Supplementary Data 3). Consistent with the observed redox sensitivity of most Tsa1 interactors, candidates common for Tsa1^C171S^ and Tsa1^C48S^ showed an enrichment tendency towards the resolving mutant (Fig. 5b). Considering only candidates that were found in triplicates as Tsa1^C171S^ interactors, we found 552 proteins (Fig. 5c). 60% of the cysteines in the corresponding peptides were modified by iodoacetamide, indicating that they were in the thiol form in their native state. Close to twenty (19.5 %) and a bit more than twenty (20.4 %) were N-ethylmaleimide modified or unmodified, respectively (Fig. 5c). There was a clear enrichment for the CxxC motif in common Tsa1^C171S^/Tsa1^C48S^ eluted interactors (Supplementary Fig. 5a). In line, the CxxC motif also was found to be prominent in peroxiredoxin-1 and −2 interacting partners^48^. In contrast, C171S-specific interaction partners did not show this enrichment indicating a different mechanism of interaction. Of note, the number of cysteines per protein follows the same trend as for the Tsa1^WT^-TAP and Tsa1^C171S^-TAP redox-sensitive interactors with C171S interactors having a slightly higher numbers of cysteines per protein, despite there is again no correlation to linear distance between cysteines or an amino acid motif surrounding the cysteines (Fig. 5d and Supplementary Data 4). Moreover, almost half of the C171S DTT-eluted interactors were also found in the Tsa1^C171S^ interactome (Fig. 5e). A gene ontology analysis, performed as we did for the Tsa1^WT^-TAP and Tsa1^C171S^-TAP interactors, revealed again that the oxidative sensitive cellular processes related to protein production and carbohydrate metabolism were enriched (Fig. 5f and Supplementary Data 4). Comparing the two complementary mass spectrometry approaches revealed a vast redox-sensitive interaction network of Tsa1 within the proteome.

**Fig. 5.**
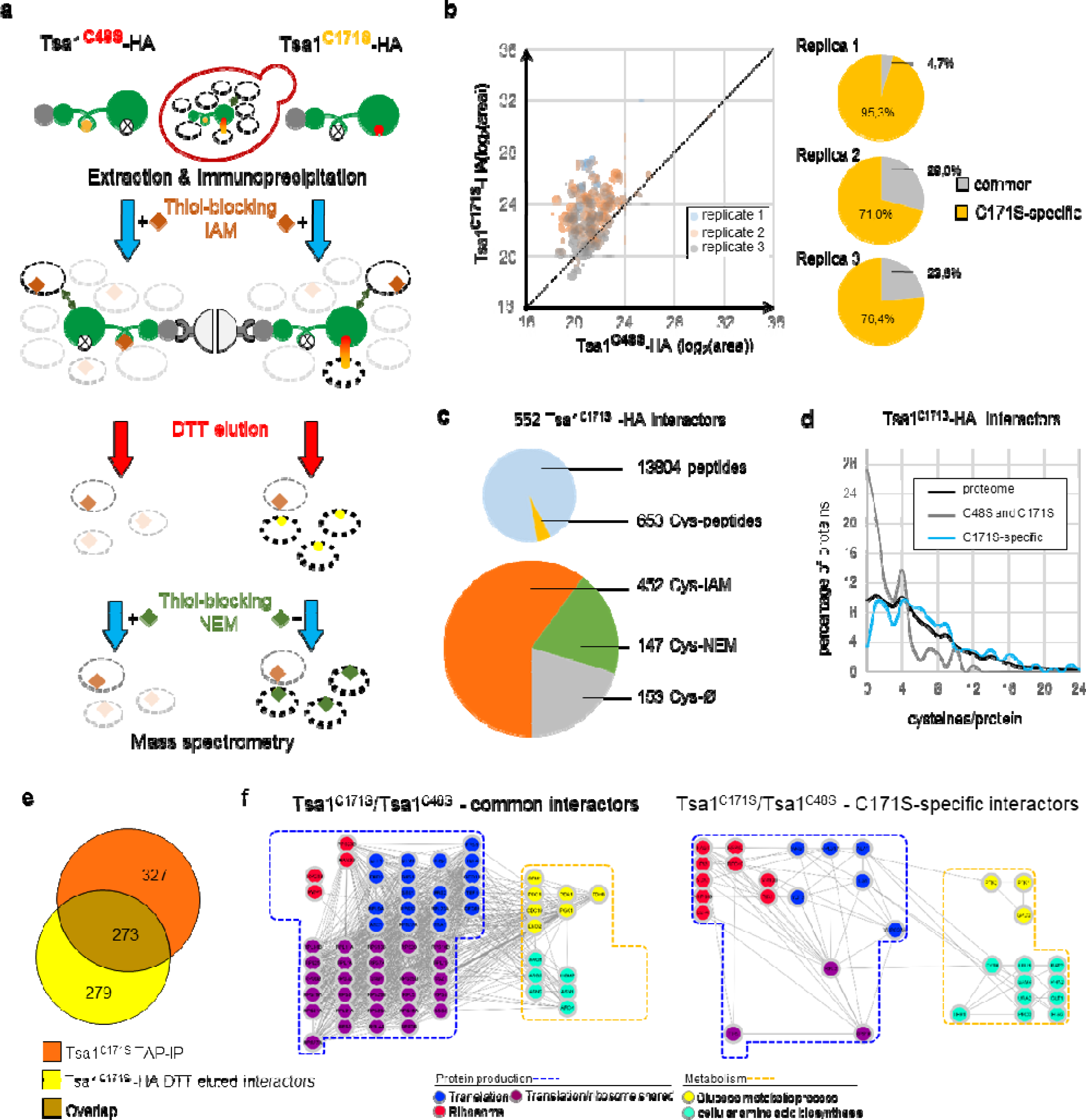
Sequential cysteine labeling identifies potential covalently associated targets to Tsa1. **a,** Schematic workflow of the mass spectrometry experiment to identify redox sensitive Tsa1^C171S^-HA and Tsa1^C48S^-HA interactors. Symbols are the same as used in Fig. 4a. C48S and C171S mutants are depicted by replacing the respective cysteine symbol by a circle with an *x*. Differential thiol blocking agents are indicated by a brown rhombus (iodoacetamide) or a green rhombus (N-ethylmaleimide). **b,** Representation of the interactors identified by mass spectrometry performed in triplicates. DTT-eluted candidates identified in both mutants are plotted Tsa1^C171S^-HA versus Tsa1^C48S^-HA (log_2_ area values). Pie diagrams for the three replicates show common candidates (identified in Tsa1^C171S^-HA and Tsa1^C48S^-HA) in grey and interactors that are specific for Tsa1^C171S^-HA in orange. The data set can be found in Supplementary Data 3. **c,** 552 interactors have been identified in all triplicates in the Tsa1^C171S^-HA DTT-elutions. Upper pie diagram shows the number of overall and cysteine containing peptides. The lower pie diagram indicates the number of cysteines that are carbamylated (Cys-IAM, modified by iodoacetamide during lysis), modified by N-ethylmaleimide (Cys-NEM, modified by N-ethylmaleimide after DTT-elution) or unmodified (Cys-Ø). Absolute numbers are next to the pie charts. **d,** Percentage of proteins (y-axis) containing the indicated cysteine content/protein (x-axis). DTT-eluted interactors that are found in both mutants (C48S and C171S) are shown in grey, C171S-specific interactors are shown in blue and the proteome wide distribution as black line. Data are found in Supplementary Data 4. **e,** Overlap (brown) of interactors identified in the Tsa1^C171S^-TAP immunoprecipitations (orange, Fig. 4) and DTT-eluted Tsa1^C171S^-HA interactors identified in this experiment (yellow). Numbers of the interactors are indicated within the sections. **f,** Representative GO terms revealed by the PANTHER GO enrichment analysis. Tsa1 interactors are visualized in interaction networks using the Cytoscape platform. Full list of enriched GO terms with FDR and p-values are found in Supplementary Data 4.

### Tsa1 buffers metabolism and associates with metabolic enzymes

Tsa1 regulates gluconeogenesis by interacting with the pyruvate kinase Cdc19, and metabolomics has revealed alterations in gluconeogenesis and the pentose phosphate pathway (PP pathway) in *tsa1*Δ*tsa2*Δ cells^49^. High molecular weight forms of Tsa1 and Cdc19 were observed, but it was not known whether they associated covalently^50^. Since peroxiredoxins appear to have a profound role in regulating metabolism we explored the effect of cysteine point mutations in Tsa1 alone were sufficient to alter the metabolomic profile by an unbiased metabolomics approach. We introduced the *TSA1^C48S^*and *TSA1^C^*^171^*^S^*mutations into a FY4 background (completely auxotrophic strain), grew cells in complete synthetic media in standard conditions and analyzed changes in their metabolome by mass spectrometry (Fig. 6a, Supplementary Fig. 6a,b and Supplementary Data 5). In general, changes were subtle, which is not surprising given the largely unaffected cellular growth of the *TSA1^C48S^*and *TSA1^C^*^171^*^S^*mutants in standard growth conditions (Supplementary Fig. 6c). Indeed, mutation of either cysteine leads to several metabolic alterations. As an expected result of a disturbed redox balance of the Tsa1 mutants, the redox pair glutathione and glutathione disulfide were both increased. Interestingly, while the GSH/GSSG ratio is not significantly affected in the Tsa1^C48S^ mutant, there was a clear decrease in the Tsa1^C171S^ mutant, indicative of a more oxidizing environment^51,52^. This is of interest since the increased oxidative environment could contribute to the high TIMDI levels in the Tsa1^C171S^ mutant. Of note, while the majority of metabolites of the “upstream” glucose metabolism showed an increase in both point mutants, phosphoenolpyruvate (PEP) decreased. Nevertheless, the alteration of the metabolome of both cysteine mutant confirm important roles for both, the peroxidative as well as the resolving mutant, in maintaining correct functioning metabolism. Thus, the global changes in metabolism observed in the Tsa1 point mutants suggested a key role for Tsa1-interaction in the regulation of metabolic enzymes as predicted from the Tsa1-interactome analyses.

**Fig. 6.**
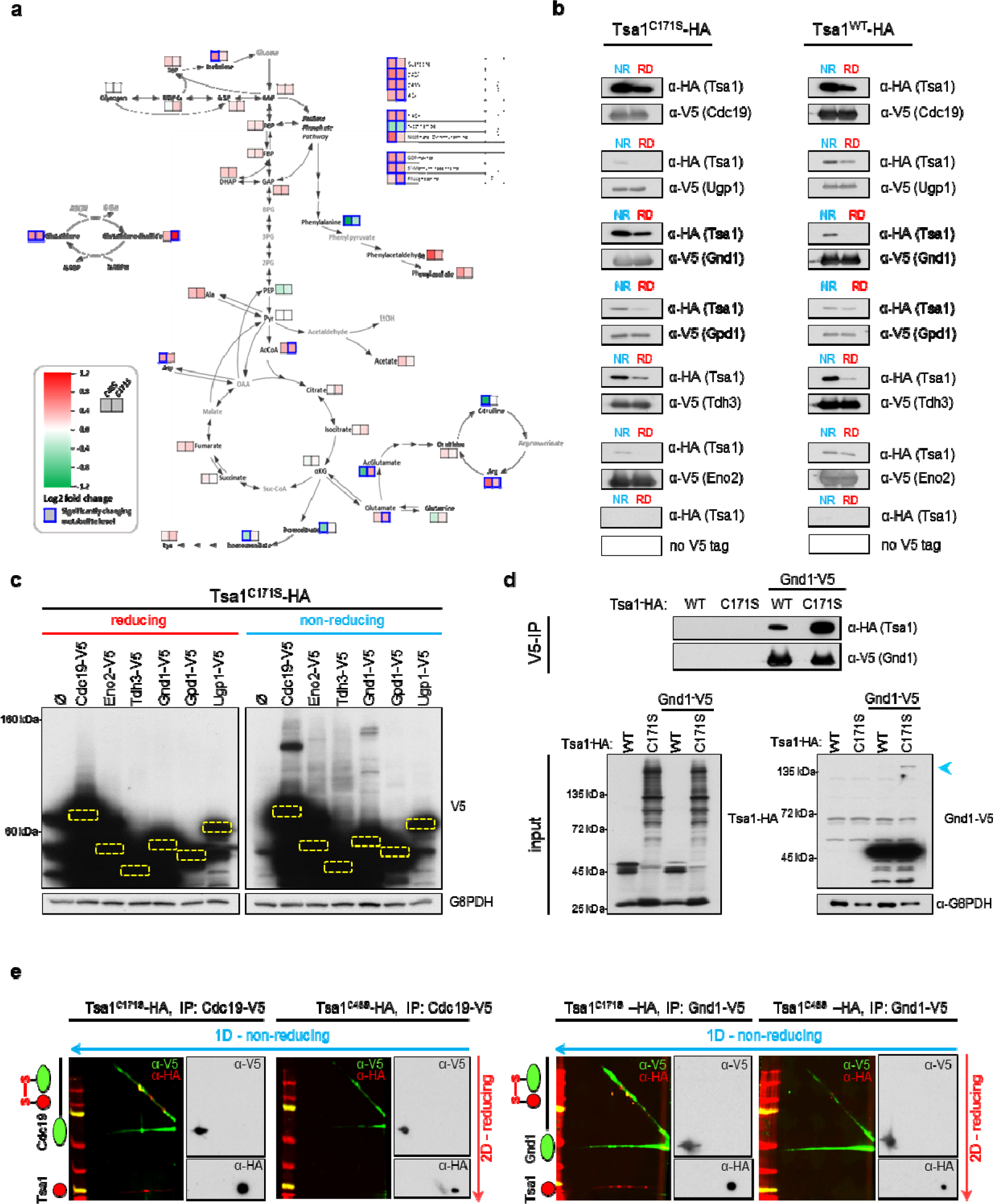
Tsa1 targets redox-sensitivity metabolic enzymes and buffers metaboli. **a,** Untargeted metabolomics reveals differences in glucose metabolism and – related - the TCA cycle of Tsa1^WT^-HA, Tsa1^C48S^-HA and Tsa1^C171S^-HA cells. Changes in metabolites in the C48S and C171S relative to the wild type are shown. The data set can be found in Supplementary Data 5. Analysis of statistical significance can be found in Supplementary Fig. 6a,b. **b,** Tsa1 physically interacts with metabolic enzymes. Single V5 tagged metabolic enzymes were immunoprecipitated under reducing and non-reducing conditions from Tsa1^WT^-HA and Tsa1^C171S^-HA strains. Immunoblots were probed for anti-HA (Tsa1) and anti-V5 (metabolic enzymes). Corresponding blots of whole cell extracts can be found in Fig. 6c and Supplementary Fig 6d,e. **c,** Whole cell lysate samples from the same Tsa1^C171S^-HA cultures used for the co-immunoprecipitation in Fig. 6b were separated by 10%15% SDS-PAGE and immunoblots probed for the V5-tagged metabolic enzymes. Probing the same membranes with anti-G6PDH was used as loading control. Dashed-line yellow squares indicates the location of the monomeric metabolic enzymes (see Supplementary Fig. 6e for shorter exposure). **d,** Tsa1^C171S^-HA shows increased binding to Gnd1-V5 compared to Tsa1^WT^-HA. Tsa1^WT^-HA and Tsa1^C171S^-HA strains ± Gnd1-V5 were used to immunoprecipitate Gnd1-V5 under non-reducing conditions. Immunoprecipitations were separated by reducing SDS-PAGE and immunoblots were probed for anti-HA (Tsa1) and anti-V5 (Gnd1). Inputs were separated by non-reducing dual 10%15% SDS-PAGE and were probed for anti-HA (Tsa1), anti-V5 (Gnd1) and anti-G6PDH as loading control. Mixed disulfide of Gnd1-V5 is indicated by blue arrowhead. **e,** Western blot membranes of 2D-SDS-PAGE of V5-immunoprecipitation of Cdc19-V5 and Gnd1-V5. Immunoprecipitations were separated under non-reducing conditions in the first dimension (right to left). Gel pieces were treated for 5’ with 5 mM DTT to achieve partial reduction and run in the second dimension (top down). Region of high molecular weight adducts /TIMDIs was probed simultaneously with mouse anti-V5 (Gnd1-V5 or Cdc19-V5) and rabbit anti-HA (Tsa1-HA). Blots were than developed with the LI-COR Odyssey^®^ system, using anti-mouse (red) and anti-rabbit (green) secondary fluorescence labelled antibodies. Membrane parts containing monomeric metabolic enzymes and Tsa1 were probed by same primary antibodies, but developed sequentially using conventional HRP secondary antibodies.

To validate the interaction of Tsa1 with carbohydrate metabolism enzymes, we performed co-immunoprecipitation experiments with V5-tagged potential target proteins (i.e., Cdc19, Gnd1, Tdh3, Gpd1, Ugp1 and Eno2) under reducing and non-reducing conditions. We found that all of them physically interacted with Tsa1^WT^-HA and Tsa1^C171S^-HA (Fig. 6b). Corresponding whole cell extracts separated by SDS-PAGE revealed DTT-sensitive high molecular weight adducts of the enzymes specifically in the non-reducing conditions (Fig. 6c and Supplementary Fig. 6d,e). In addition to the confirmation CoIPs we made a direct comparison of the Gnd1-V5 interaction with wild-type and C171S mutant Tsa1 (Fig 6d). As expected Tsa1^C171S^ mutant has a stronger interaction with Gnd1-V5 than Tsa1^WT^-HA (Fig. 6d). As expected from the low amount of TIMDIs, HMW adducts of the metabolic enzymes are barely detectible under standard growth conditions. In line with TIMDI increase upon environmental impacts, metabolic enzymes HMW adducts are increased upon stress (Supplementary Fig. 6f).

To confirm that the observed mixed disulfide intermediates of metabolic enzymes correspond to TIMDIs we performed a two-dimensional SDS-PAGE analysis of Gnd1-V5 (6-phosphogluconate dehydrogenase) and Cdc19-V5. To this aim Gnd1-V5 and Cdc19-V5 were immunopurified under no-reducing conditions from a Tsa1^C171S^-HA and Tsa1^C48S^-HA strain. After separation on a SDS-PAGE in the first dimension, the lane was cut and reduced by DTT. Subsequently the gel fragment was separated in the 2^nd^ dimension. Indeed, high molecular weight adducts of both, Gnd1-V5 and Cdc19-V5, co-localize with Tsa1^C171S^-HA and could be split by reduction to the monomeric V5-tagged enzymes and Tsa1^C171S^-HA (Fig. 6e). Of note, the V5 tag of Gnd1 and Cdc19 does not affect survival nor catalytic activity of these enzymes and CoIPs with Gnd1-V5 or untagged Cdc19 with endogenous Tsa1 yielded the same results (Supplementary Fig. 7a-f). Thus, the global changes in metabolism observed in the Tsa1 point mutants suggested a key role for Tsa1 in the regulation of metabolic networks as predicted from the Tsa1-interactome analyses.

### Cys460 is an important residue for of Gnd1 activity which is targeted by Tsa1

Gnd1 is a central enzyme of the oxidative branch of the pentose phosphate pathway that has a strong redox-sensitive interaction with Tsa1 (Fig. 6b-e). The pentose phosphate pathway fuels the cell with NADPH, a key metabolite in redox reactions and oxidative stress protection^53,54^. Gnd1 forms more than a single high molecular weight species in the Tsa1^C171S^-HA background, indicating that more than one cysteine could be involved in the covalent interaction with Tsa1 (Fig. 6c and 6e). In contrast, in a Tsa1 wild-type strain, Gnd1-V5 forms a single predominant high molecular weight species that was induced by H_2_O_2_ and depend on the peroxidatic cysteine (Fig. 7a,d and Supplementary Fig. 8a). Moreover, covalent attachment to the preferred target protein cysteines to the peroxidatic cysteine of Tsa1 seems to prevent undirected mixed disulfide formation, judged by the increase of Gnd1-V5 adducts in the Tsa1^C48S^-HA mutant (Fig.7a and Supplementary Fig. 8a). To understand the nature of this covalent interaction, we created a series of mutants in all cysteines of Gnd1-V5 in the genomic context. While mutations in some residues showed a reduction of TIMDIs, others resulted in a strong increase (Fig. 7b,c). Importantly, independently of an increase or decrease, the high molecular weight adducts of all GND1-V5 variants were dependent on Tsa1 since they disappeared in a *tsa1*Δ deletion background (Supplementary Fig. 8b). The Gnd1^C460S^ mutation in the Tsa1^C171S^-HA background completely abolished the prevailing Gnd1 high molecular weight adduct and introducing the C460S mutation in Gnd1 in a Tsa1^WT^-HA background led to the loss of the main Gnd1 TIMDI, making C460 the primary cysteine targeted by Tsa1 (Fig. 7b-e). In addition to the loss of the predominant Gnd1-TIMDI species, Gnd1^C460S^ led to an increase in mixed disulfide intermediates of various sizes (Fig. 7e and Supplementary Fig. 8c).

**Fig. 7.**
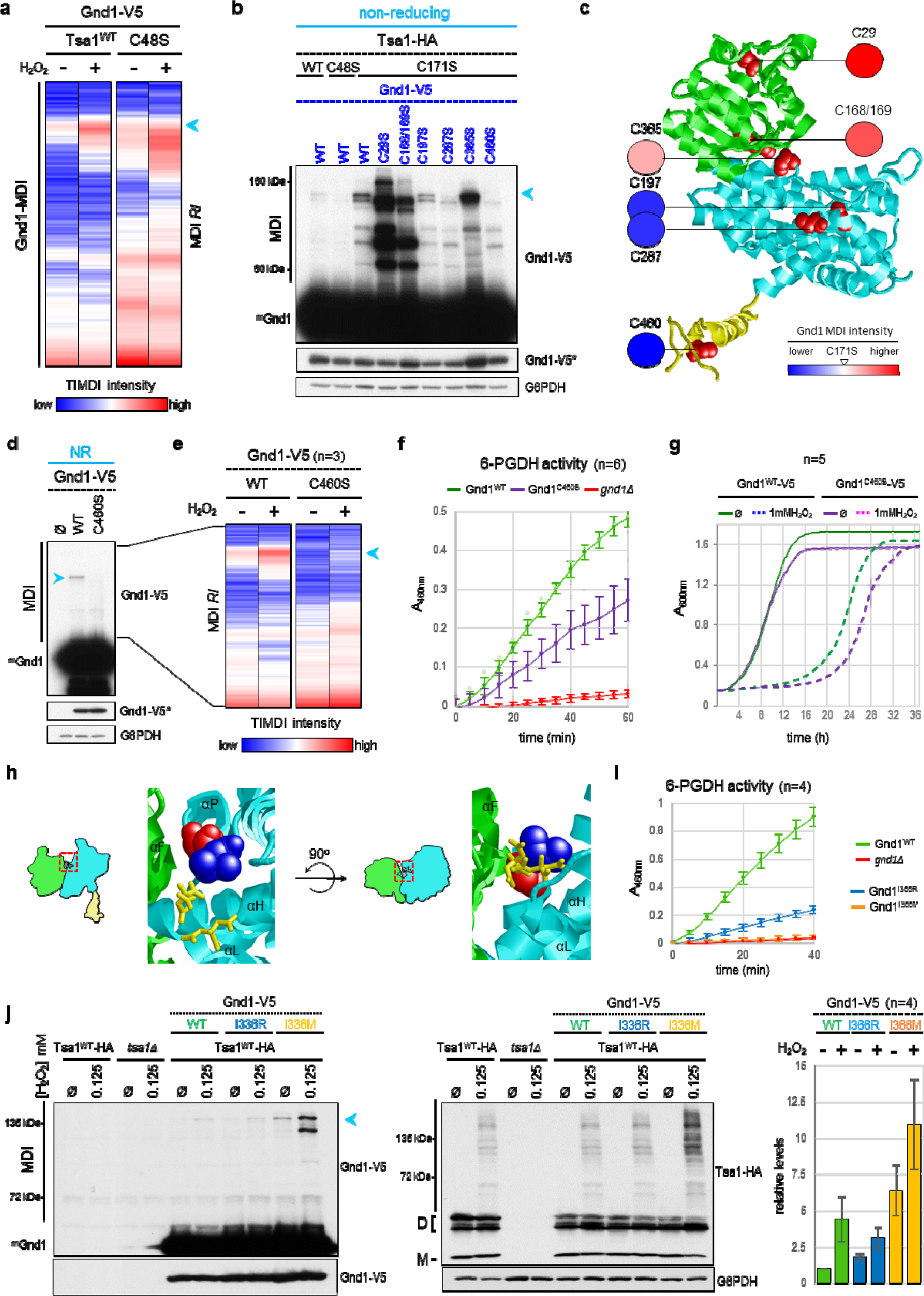
Gnd1 C460 a cysteine relevant for enzymatic activity and survival upon oxidative stress is targeted byTsa1. **a,** Heat map of relative Gnd1-MDI intensity values was generated from biological triplicates. Relative MDI intensity are shown with lowest in blue and highest intensity in red. Log_2_-fold change and statistical analysis can be found in Supplementary Fig. 8a. The predominant adduct of Gnd1^WT^-V5 is indicated by a blue arrowhead. **b,** Cysteines in Gnd1-V5 were mutated one by one (with the exception of the di-cysteine motif C168/C169) in the Tsa1^C171S^-HA background. Non-reducing whole cell extracts were separated on dual 10%15% SDS-PAGE and immunoblots were probed with anti-V5 (Gnd1 versions; long exposure to visualize Gnd1-V5 MDIs and short exposure (*) to visualize monomeric Gnd1-V5) and anti-G6PDH (loading control). Western blots for the same strains in a *tsa1*Δ can be found in Supplementary Fig 8b. **c,** Gnd1 crystal structure (2P4Q) with the N-terminal A domain (NADP+ binding, green), the central B domain (substrate binding, blue), and the C-terminal extension (lid formation, yellow). Structure is shown as ribbons with the exception of the cysteines as space fill models (red). Position of the cysteines is indicated. Filled circles show the high molecular weight adduct changes relative to the Gnd1^WT^-V5 in the Tsa1^C171S^-V5 background (lower in blue, equal in white, increase in red, relative values) **d,** Non-reducing SDS-PAGE and subsequent western blotting of whole cell extracts of strains harboring a Gnd1^C460S^-V5 mutant or wild-type Gnd1-V5 introduced in the Tsa1^WT^-HA background. Membranes were probed for anti-V5 (Gnd1) and anti-G6PDH (loading control). **e,** Heat map of relative MDI intensities of Gnd1^WT^-V5 and Gnd1^C460S^-V5 in a Tsa1^WT^-HA background of the indicated region in Fig.7d was generated from biological triplicates. Relative TIMDI intensity are shown with lowest in blue and highest intensity in red. Log_2_-fold change and statistical analysis can be found in Supplementary Fig. 8c. Cells were unstressed or treated 10’ with 0.125 mM H_2_O_2_ (n=3). The predominant adduct of Gnd1^WT^ is indicated by an blue arrowhead. **f,** Gnd1^C460S^-V5 has reduced 6-phosphogluconate dehydrogenase activity. 6-PGDH activity in extracts of Gnd1^WT^-V5, Gnd1^C460S^-V5 and *gnd1*Δ cells was measured (n=6). Relative activity corrected for background is shown as median values ± SD. Single data points are depicted as circles. **g,** Growth curves of the Gnd1^WT^-V5 and Gnd1^C460S^-V5 strains in liquid media ± 1 mM H_2_O_2_. Circles show A_600_ measurements every 10’ to follow cellular growth. **h,** Location of amino acid residues C365 (space fill red) and I366 (space fill blue) at the Gnd1 domain A (green) and B (cyan) interface. Location of the two citrate molecules in the substrate binding pocket is shown as stick model in yellow. **i,** Mutation of the I366 residues reduces 6-phosphogluconate dehydrogenase of Gnd1. 6-PGDH activity in extracts from Gnd1^WT^-V5, Gnd1^I366R^-V5, Gnd1^I366M^-V5 and *gnd1*Δ cells grown in YPD was measured. **j,** Non-reducing whole cell extract of Gnd1^WT^-V5 and I366 mutants with and without 0.125 mM H_2_O_2_ treatment were separated on on-reducing SDS-PAGE and western blot membranes was probed for V5 (Gnd1) and HA (Tsa1) to visualize the respective high molecular weight adducts. G6PDH is displayed as loading control.Tsa1^WT-^HA and tsa1Δ strain were included as control. Left panel shows the quantification of the blue arrowhead indicated Gnd1-V5 MDI in triplicates. **b,d,j,** mGnd1 indicates the monomeric Gnd1-V5. Peroxiredoxin and their cysteines are depicted as in Fig. 1a.

The C-terminus of Gnd1 is indispensable for its activity, stabilizes the dimer and forms a lid on the substrate-binding pocket^55^. Thus, C460 might be an important residue for Gnd1’s catalytic activity and therefore we assayed Gnd1 enzymatic activity of this mutant comparing it to the wild type. Extracts from wild-type Gnd1-V5, the Gnd1^C460S^-V5 mutant and *gnd1Δ* cells were assessed in a colorimetric enzymatic assay that measures the ability of cell extracts to metabolize NADP+ to NADPH using 6-phosphogluconate as substrate. Accordingly, we found that the Gnd1^C460S^-V5 mutant showed reduced 6-phosphogluconate dehydrogenase activity (Fig. 7f). Remarkably, this mutant showed a clear reduction in cell fitness in the presence of oxidative stress (Fig. 7g). Thus, we have identified C460 of Gnd1 as an important residue for phosphogluconate dehydrogenase activity by deciphering Gnd1 peroxiredoxinylation. Modification by Tsa1 could protect Gnd1 from unwanted MDI formation and thereby regulate enzyme function in a redox manner.

In contrast to Gnd1 C460S, the C365S mutant showed strongly increased high molecular weight adducts in the Tsa1^C171S^-HA background (Fig. 7b,c). Cys365 is located at the domain A/B interface in the substrate-binding groove and we speculated that misfolding or enzymatic defects would lead to covalent attachment of Tsa1 as a quality control mechanism. To test this hypothesis, we selected isoleucine 366, a neighboring residue of Cys365 that is not involved in substrate binding, but resides in the binding groove (Fig. 7h) and mutated it to arginine or methionine. Indeed, Gnd1^I366R^ and Gnd1^I366M^ abolished or strongly reduced 6PGDH activity, respectively (Fig. 7i). Consistently, those mutants have increased Gnd1 HMW adducts that were abolished in the *tsa1*Δ deletion mutant (Fig. 7j and Supplementary Fig.8d). There is a direct correlation between activity and the degree of high molecular adduct formation, with the GND1^I366M^ having a stronger increase in adducts than the GND1^I366R^ mutant (Fig. 7j). Thus, while C460 of Gnd1 is a direct target of peroxiredoxinylation, mutations in residues critical for folding or catalytic activity could lead to Tsa1 attachment.

### Thioredoxins regulate the extent of peroxiredoxinylation

Low basal TIMDI levels could be due to a redox environment that does not favor their formation, or negative regulation. In canonical peroxiredoxin-regulated redox signaling an unstable PRX-target mixed disulfide is catalyzed to a disulfide bond within the target protein^26^. The absence of a (structural) vicinal cysteine to Gnd1 C460, and the formation of mixed disulfide intermediates through multiple cysteine residues in Gnd1 mutants challenge the canonical redox relay as observed e.g. in Bcy1^22^. Could a target peroxiredoxinylation mark that does not subsequently result in an oxidized target protein act as a recruiting signal for the protein folding machinery? In aged cells and upon oxidative stress, Tsa1-SO_2/3_ binds to the Hsp70 chaperone and recruits the Hsp104 disaggregase to aggregated proteins. In this way, protein aggregations are resolved and oxidized Tsa1 is recycled via the sulfiredoxin Srx1^30^.

We identified Hsp70 (Ssa1/2), Hsp104, Srx1 and thioredoxins (Trx1/2) in our Tsa1 interactome (Supplementary Data 1 and 3). Deletion of Ssa1, Ssa2, Hsp104 and Srx1 did not result in an altered basal TIMDI pattern. Noteworthy, while the specific Srx1-Tsa1 covalent adduct disappears general TIMDIs are unaffected in the *srx1*Δ mutant upon challenge with 0.125 mM H_2_O_2_ (Supplementary Fig. 9a). In marked contrast, the deletion of thioredoxins (*trx1*Δ*trx2*Δ) resulted in a burst of TIMDIs (Fig. 8a and Supplementary Fig. 9a,b). Following TIMDI formation from 0.031 to 1mM H_2_O_2_ shows an increase of TIMDIs in *trx1*Δ*trx2*Δ double mutant at all concentrations (Supplementary Fig. 9c). Like in the Tsa1 C171S mutant, the *trx1*Δ*trx2*Δ double mutant shows strongly impaired peroxidatic cysteine hyperoxidation (Supplementary Fig. 9d). We speculate that while in the C171S mutant Tsa1 oxidized dimers cannot form and the sulfenic form rapidly reacts with cysteines on interactors, in the *trx1*Δ*trx2*Δ mutant part of the sulfenic Tsa1 forms TIMDIs and part further oxidizes to the fully oxidized dimer, as observed in Supplementary Figure 9a. Importantly, introducing a plasmid with the genomic region encoding *TRX1* or *TRX2* do partially rescue basal as well as 0.125mM induced TIMDI formation (Supplementary Fig. S9e). Thus, both cytoplasmic thioredoxins in yeast are able to resolve TIMDIs and most likely for full control both genes have to be functionally present. Additionally, the absence of thioredoxins led to strong mixed disulfide induction of Gnd1-V5, which was dependent on its C460 (Fig. 8b).

**Fig. 8.**
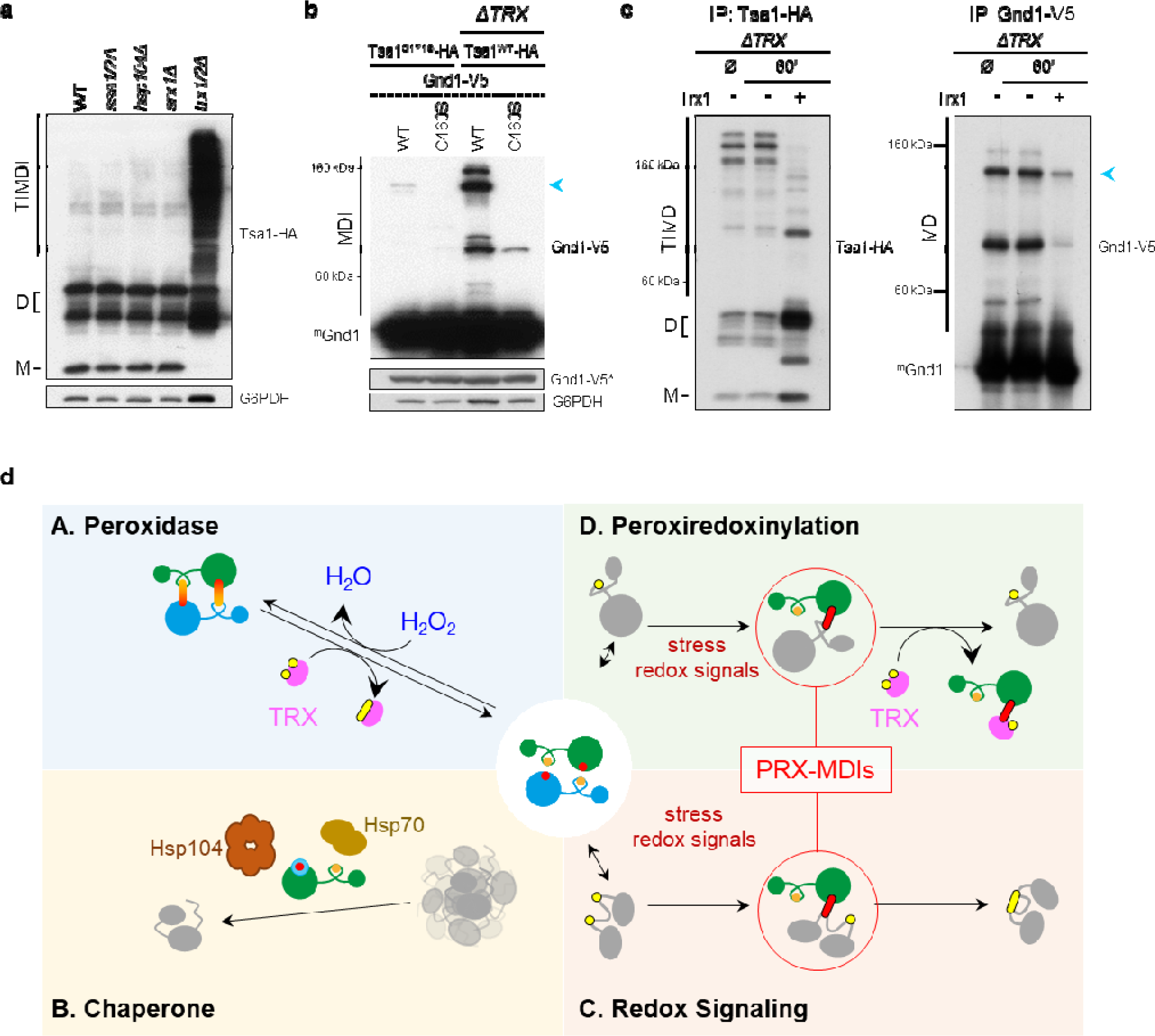
Thioredoxins directly resolve Tsa1-induced mixed disulfides. **a,** Deletion of thioredoxins leads to high basal TIMDI levels. Non-reducing whole cell extracts of *hsp70*Δ (*ssa1*Δ*ssa2*Δ), *hsp104*Δ, sulfiredoxin (*srx1*Δ) and thioredoxin (*trx1*Δ*trx2*Δ) strains having Tsa1^WT^-HA were separated by dual 10%15% SDS-PAGE and immunoblots were probed for anti-HA (Tsa1) and G6PDH (loading control). Quantification of the TIMDI extent of *trx1*Δ*trx2*Δ mutants is found in Supplementary Fig. 9b. **b,** Gnd1-V5 high molecular weight adducts in a thioredoxin deletion (*trx1*Δ*trx2*Δ) strain. Non-reducing whole cell lysate of Gnd1^WT^-V5 and Gnd1^C460S^-V5 in Tsa1^C171S^-HA or *trx1*Δ*trx2*Δ Tsa1^WT^-HA background were separated on a SDS-PAGE and immunoblots were probed by anti-V5 (Gnd1) and anti-G6PDH (loading control). Gnd1-V5 monomer is visualized by a lower exposure (*). Blue arrowhead indicate the major Gnd1-V5 MDI band. **c,** Thioredoxins efficiently remove peroxiredoxinylation *in vitro*. Immunopurification of Tsa1^WT^-HA (left panel) or Gnd1-V5 (right panel) from a thioredoxin deletion background were incubate ± recombinant Trx1-HIS for 60’. Blots were probed for anti-HA (Tsa1) or anti-V5 (Gnd1). Blue arrowhead indicate the major Gnd1-V5 MDI band. Probing membranes for Tsa1-Trx1 adducts as a product of TIMDI resolution can be found in Supplementary Fig. 9f. **d,** Mechanistically model of TIMDIs in the context of peroxiredoxin function. (A) The core catalytic redox cycle of the peroxiredoxin peroxidase function. Hydrogen peroxide oxidizes peroxiredoxins (PRX) which are subsequently reduced by the thioredoxins (TRX). Not shown is the recycling of TRX by the thioredoxin reductase. (B) Outlines the chaperone formation of peroxiredoxin as a result of peroxidatic cysteine hyperoxidation upon oxidative stress. Severe oxidative stress insults can lead to protein aggregation and hyperoxidation (SO_2/3_) of peroxiredoxins. Hyperoxidized PRX localizes to the aggregates and recruits the disaggregase Hsp104 via Hsp70 interaction. Sulfiredoxin (SRX) subsequently recycles PRX-SO_2_ and thereby protect it from irreversible oxidation to SO_3_ and degradation. (C,D) Tsa1-Induced Mixed Disulfide Intermediate (TIMDI) formation as a result of redox imbalance caused by H_2_O_2_ or other stresses. Tsa1 forms covalent disulfide bridges with thiol groups of target proteins as a result of moderate H_2_O_2_ and other stresses. Those adducts can be resolved by (C) a disulfide exchange reaction by a vicinal cysteine in the target protein leading to a recycled peroxiredoxin and an oxidized target that act e.g. as an effector of redox signaling or (D) thioredoxins in the absence of a vicinal cysteine. In this case the target protein is released in the reduced thiol form, not changing the redox state of the protein and an PRX-TRX adduct that can be resolved by the thioredoxin reductase. C48-SO_2/3_ is shown as a blue, red-filled circle. Target proteins are shown in grey. important cysteines in target proteins and thioredoxin are shown as yellow filled circles. Disulfide bridges are indicated as fused circles.

Comparable to the C171S mutant, a more oxidizing environment likely contributes to high TIMDI levels in the *trx1*Δ*trx2*Δ double mutant. Additionally, in support for a direct, enzymatic targeting of thioredoxins in TIMDI resolution, we found that recombinant Trx1-HIS resolved general TIMDIs, as well as Gnd1-Tsa1 TIMDIs *in vitro* (Fig. 8c). Furthermore, in *in vitro* assays, we detected only Tsa1-HA - Trx1-HIS adducts, indicating that TIMDIs are resolved by transferring the attached Tsa1 to Trx1 (Supplementary Fig. 9f). *in vivo* the Tsa1-Trx1 adduct would be subsequently resolved by an intramolecular disulfide formation in the thioredoxin, resulting in a reduced peroxiredoxin and a oxidized thioredoxin. The oxidized thioredoxin can subsequently be resolved by the thioredoxin reductase (TRR) system. In conclusion, thioredoxins and not the Hsp70/Hsp104 complex are required to target Tsa1-mixed disulfide intermediates for resolution.

## Discussion

Peroxiredoxins were described as bifunctional enzymes that carry H_2_O_2_-detoxifying peroxidase and molecular chaperone functions^17,56,57^. It is now clear that these two functions merely provide the frame work for the diversity of the cellular roles of peroxiredoxins such as redox signaling ^58,59^. For the redox functions, it is important to activate the peroxidatic thiol group into a reactive sulfenic species, that either directly forms a mixed disulfide intermediate with a redox signaling protein or forms a C_P_-C_R_ disulfide homodimer intermediate. In both cases, two structurally close cysteines subsequently form a disulfide, resulting in an oxidized redox regulator or oxidized homodimer. Studying the major yeast peroxiredoxin Tsa1, we found that it is able to form widespread Tsa1-induced mixed disulfides that depend on the peroxidatic cysteine. Similar mixed disulfides have been observed for several organisms arguing for an evolutionary conserved process^21,24,60,61^. Here, we show that mutating the resolving cysteine C171 - which abrogates the C_P_-C_R_ disulfide linked dimeric form - does not impede TIMDI formation and even strongly enhance them. This implies that a C_P_-C_R_ disulfide exchange with a target cysteine is not necessary in TIMDI formation. We found that at H_2_O_2_ stress conditions that do not lead to hyperoxidation of the peroxidatic cysteine result in a strong induction with about 20% of the cellular Tsa1 pool incorporated into Tsa1-Induced Mixed disulfides (TIMDIs). Importantly the formation of covalent Tsa1 adducts does not involve *de novo* protein production nor requires the Hsp70/Hsp104 system, two processes Tsa1 has been described to be involved to prevent protein aggregation^20,30^. Instead, we found that mature proteins are targets for TIMDI formation and deletion of the thioredoxins result in a strong increase of TIMDIs, involving up to 60% of the cellular Tsa1 pool. While our results do not exclude that a change to a more oxidizing cellular environment contribute to TIMDI formation in the Tsa1^C171S^ and thioredoxin deletion mutant, we clearly show that *in vitro* assays that Tsa1^C171S^ can directly form TIMDIs with mature proteins and that TIMDIs can be efficiently resolved by thioredoxins supporting a direct role in TIMDI resolution by thioredoxins.

In wild-type cells we found a strong TIMDI formation upon stresses that can directly affect protein function and elevate the basal oxidative stress level, such as arsenite stress, osmotic stress or heat shock^62–64^. Interestingly, TIMDI formation and cycloheximide resistant foci formation correlate in those stresses. Thus one possibility is that target proteins and peroxiredoxins are concentrated in such foci, which could be the trigger for rapid TIMDI formation. Strikingly, while the wild type and the peroxidatic cysteine mutant (C48S) are sensitive to arsenite stress, the C171S mutant has a marked growth advantage under this condition. In line with the model that increased TIMDI can protect against stress a mutant on the N-terminal lobe of Tsa1, Y78A, that strongly impairs decamer formation concomitantly reduces TIMDIs and renders C171S sensitive - and cells with otherwise wild-type Tsa1 even more susceptible - to arsenite stress.

Since this is the first report that shows that the C171S has a pronounced beneficial role in a certain stress conditions we deciphered the redox-sensitive interactome of *S. cerevisiae* Tsa1^WT^ and Tsa1^C171S^. Here we show that peroxiredoxins interact with hundreds of target proteins in a redox sensitive manner. The peroxidatic cysteine is crucial for many of those interactions and we find that peroxiredoxinylation could be a widespread process involving proteins in key cellular processes, including translation and metabolism. A critical point is if the high TIMDI formation of Tsa1^C171S^ is simply an artifact generated by the resolving cysteine mutation. We argue with several observation against this possibility: First, there is a strong overlap of interactors of Tsa1^WT^ and Tsa1^C171S^ discovered by mass spectrometry. Second, we confirmed the interaction of various glycolytic enzymes interactions of Tsa1 with Tsa1^WT^ and Tsa1^C171S^. Next, while the C171S mutant has no large structural alterations, a shift in equilibrium to the locally unfolded/destabilized state is observed^65^. Thus, we reason that Tsa1^C171S^ adopts a conformation *in vivo* that is distinct of the hyperoxidized chaperone, but favors covalent thiol attachment to interaction partners. This locally unfolded state can explain the hyperoxidation resistance of Tsa1^C171S^ and would allow the formation of mixed disulfides. Subsequently we speculate that environmental conditions that result in an increase of TIMDIs elicit a state of Tsa1^WT^ that at least partially correspond to that of the C171S mutant.

Considering the general low reactivity of most cysteines in proteins and the need to be selective in context of redox signaling, we asked which proteins are targeted and how the target cysteines were selected. Besides ribosomal proteins and proteins of protein turnover, processes related to glucose metabolism were strongly enriched in the Tsa1 interactome. Cdc19 has been reported to interact in a redox-sensitive fashion with Tsa1 and we showed here that this association is indeed via the formation of a covalent disulfide link between Cdc19 and Tsa1. Supporting for the role in carbohydrate metabolism, we also showed that in non-challenged conditions, large alterations of the metabolome is found in both Tsa1 cysteine mutants (Tsa1^C171S^ and Tsa1^C48S^). While our metabolomic studies does not allow to conclude on changes in fluxes, using Gnd1 (6-phosphogluconate dehydrogenase), we clearly showed that Tsa1 targets this enzyme preferentially at a residue that is important for catalytic activity and important for survival upon oxidative stress. Thus, we present the following model for peroxiredoxins function (Figure 8d): (A) in their peroxidase function peroxiredoxins detoxify hydrogen peroxide of exogenous (atmosphere) and endogenous (metabolism) origin. Oxidized, disulfide-bridge linked C_P_-C_R_ dimers are resolved by the thioredoxins, forming the core of the catalytic redox cycle. (B) Severe oxidative stress leads to the formation of the C48-hyperoxidized (SO_2_ and SO_3_) chaperone^17,57^. In this peroxiredoxin function Hsp70 and Hsp104 are recruited to refold damaged, aggregated proteins^30^. Subsequently sulfiredoxin is required to revert the sulfonylation of the peroxidatic cysteine. (C,D) However, at modest H_2_O_2_ stress and stresses that affect the redox balance and folding of mature proteins, the non-oxidized chaperone function of Tsa1 leads to increased interaction with potential targets with accessible cysteines to form covalent adducts (peroxiredoxinylation^24^), which we named Tsa1-induced mixed disulfide intermediates, TIMDIs. (C) In the case of the canonical redox relay a structural close cysteine can now attack the target-S-S-Tsa1 disulfide bond, releasing Tsa1 and leaving an oxidized redox regulator behind^19,22,66^. (D) In the absence of a vicinal cysteine, we showed that thioredoxin is sufficient and necessary to target TIMDIs, resulting in a non-modified thiol group on the target protein and a peroxiredoxins--thioredoxin linked mixed disulfide intermediate. This has several important implications and consequences. First, peroxiredoxinylation does not need an enzymatic ligation system such as seen for ubiquitin ligases or kinases (phosphorylation). The high expression levels do not only make peroxiredoxins important H_2_O_2_ detoxifiers but also increase the likelihood to interact with their targets - especially taking into account local concentration increases by foci formation upon stress. The properties of the peroxidatic cysteine allows the formation of sulfenic/sulfinic Tsa1 at low H_2_O_2_ concentration and subsequently the reaction with target thiols at moderate stress condition. In the absence of TIMDI formation, random mixed disulfides can occur that crosslink proteins that are damaging to the cell. In conclusion, we present evidence that peroxiredoxins may act as guardians of thiol reactivity on a proteome-wide scale and that peroxiredoxinylation is a bona fide post-translational modification on cysteine residues that, together with thioredoxins, controls the formation of protein mixed disulfide intermediates in the cytoplasm. In this context a signaling function by peroxiredoxin attachment to a stress-activated kinase has been described recently^23^. The large extent of peroxiredoxinylation that we observed in this study is therefore an indication that new, redox-sensitive mechanisms of peroxiredoxins are likely to be discovered in the future.

## Methods

### Yeast Cultures

Standard procedures were used for the growth and genetic manipulation of *Saccharomyces cerevisiae*. Cells were grown at 30°C in YPD or in synthetic complete medium ^59^ with 2% glucose (metabolomics). For yeast transformation (deletion or knock in mutants using a PCR product as template for homologues recombination) a 50ml culture was grown to OD_600_=0.6 - 0.8. Cells were harvested by centrifuging at 3000 rpm for 3’. Cell pellet was washed with 500μl 10mM Tris-HCl, pH 8.0, supplemented with 1mM EDTA and 100mM Lithium acetate and centrifugation step was repeated. Pellet was resuspended in 1ml of the same buffer. 100μl of cell suspension was added to a microfuge tube containing a mixture of 10μl 10mg/ml heat denatured ssDNA with 1μg of PCR product and mixed well by flicking. 600μl of 40 % PEG in 10 mM Tris-HCl, pH 8.0, with 1 mM EDTA and 100mM lithium acetate was added, tube was inverted five times and mix was incubated at 30°C for 45’. Then 70μl of DMSO was added, suspension was vortexed and heat shock at 45°C was applied for 15’. Cells were pelleted by centrifuging at 3000rpm for 1’. Pellet was resuspended in 400μl 10 mM Tris-HCl, pH 8.0 with 1 mM EDTA and was plated onto agar plates containing the auxotrophy selection medium. In case of antibiotic selection, cells were plated on YPD and replicated to antibiotic containing YPD plates using velvets 12h after transformation. GFP-tagged *S. cerevisiae* strains were obtained from the *ThermoFisher* Yeast GFP Clone Collection ^60^. The Tsa1-TAP strains was obtained from the *Horizon* Yeast TAP Tagged ORFs Collection ^61^. The single C-terminally HA-tagged strains were generated by first inserting an auxotrophic marker 100bp 3’ of the Tsa1 stop codon in the endogenous locus in a BY4741 background to generate strain YGS347. Using genomic DNA from YGS347 as template PCR cassettes for homologous integration were amplified using *TSA1 mHA f01* (forward primer containing a single HA followed by a stop codon) and *Ctag TSA1 r03* (reverse primer) oligos. This cassette has been transformed into BY4741 to generate an endogenously single HA-tagged strain, YGS447. To generate endogenously single HA-tagged C171S (YGS448) and C48S (YGS449) mutants of Tsa1 cassettes for homologous recombination were amplified using oligos *C171S wt f01* and *C48S f02*, respectively, and genomic YGS447 DNA as template. For single V5-tagging, the respective oligos found in Document S7 were used to generate PCR cassettes for homologous recombination using the pYM16 plasmid as template. To replace the promoter of Tsa1 with the promoter of Tdh3 or Rsp5 first an URA auxotrophic marker was inserted by homologous recombination in the 5’ genomic region of TDH3 (YGS1735) or RSP5 (YGS338). Genomic DNA from YGS1735/YGS338 was used as a template to generate a cassette to replace endogenous Tsa1 promoter of YGS447 strains. All *S. cerevisiae* strains generated or used in this study are listed in the Supplementary Data 7.

### Growth curves

*S. cerevisiae* strains were maintained in log-phase at least 12h before the experiment. For the experiment cells were diluted to OD_600_=0.1 and grown for 3h, then diluted to OD_600_=0.01 30’ before the start of the growth curve. Yeast cells were mixed with YPD or YPD containing stressors at 1:1 to a final volume of 200μl in 96-well plates. Plates were incubated under orbital shaking at 30°C in a Synergy H1 (BioTek® Instruments) and OD_600_ was measured every 10’.

### Recombinant protein production

TSA1^C171S^ coding sequence was amplified from genomic YGS448 DNA (see “*S. cerevisiae* strains generated in this study”) and cloned into pGEX-6P-1 using BamHI-HF and NotI-HF. The resulting plasmid was transformed into BL21 and expression was induced using IPTG for 3 h. After purification using glutathione sepharose, GST-tagged protein was eluted with L-glutathione. To remove L-glutathione buffer was exchanged to 50mM Tris_HCl, pH 7.0 supplemented with 150mM NaCl, 1mM EDTA, 1mM DTT, 0.01% NP40 using centrifugal filter columns. The GST-tag was removed by PreScission Protease. To this aim, 0.3μl of PreScission protease was added per 1μl buffer-exchanged recombinant protein and incubated at 12-16h. To the cleavage reaction 5x the volume of 50mM Tris-HCL pH8.0, 2mM DTT and 1.5x volume of the beads that were used for the initial GST-protein extraction was added and rotated for 1h at 4°C. Beads were removed by centrifugation and supernatant was concentrated using centrifugal filter columns.

TRX1 coding sequence was PCR-amplified from genomic BY4741 DNA and cloned into pETM10 using NcoI and NotI-HF. The resulting plasmid was transformed into BL21 and expression was induced using IPTG for 3 h. His-tagged Trx1 was purified on a Ni2+ sepharose resin following the manufactureŕs instructions (cOmplete™ His-Tag Purification Resin, Merck).

### *In vitro* TIMDI resolution by thioredoxins

Gnd1-V5 or TSA1-HA were immunopurified from a Δ*trx1*Δ*trx2* mutant background using 10 ml culture of OD_600_= 0.8 per condition under non-reducing conditions. All subsequent steps were conducted in an N_2_ atmosphere: 2 μg of His-tagged Trx1 was added to beads suspended in reaction buffer (50 mM Tris-HCl, 1 mM EDTA, 150 mM NaCl, pH 7.5) and incubated at 30°C under shaking at 600 rpm. The reaction was stopped by adding the same volume of reaction buffer supplemented with 20 mM N-ethylmaleimide. Non-reducing sample buffer was added to a final concentration of 50mM Tris-HCl, 2% SDS, 10% glycerol, and 0.05% bromophenol blue. After samples were heated to 95°C for 2’ they were separated by SDS-PAGE and transferred to PVDF membranes.

### *In vitro* peroxiredoxinylation

*tsa1*Δ cells were harvested at an OD_600_=0.5 and washed once in ice-cold 1xPBS, and whole-cell extracts were obtained by glass bead lysis in 50 mM Tris-HCl pH 8.0 and 2 mM DTT. For cysteine-blocked whole-cell extracts, 20mM N-ethylmaleimide was added to the buffer during extraction, followed by incubation for 30 min on ice, and buffer exchange to 50mM Tris-HCl pH 8.0 containing 2mM DTT was done using centrifugal filter columns. For the peroxiredoxinylation reaction, 400ng of whole cell extracts and 200ng of recombinant Tsa1^C171S^ was mixed and dialyzed 3x against a 10kDa centrifugal filter column. After incubating the mix for 15 min at 30°C, the reaction was stopped by adding a double volume of 20mM iodoacetamide and dialyzing it two-times against a 10kDa centrifugal filter column at room temperature. Non-reducing sample loading buffer (final concentration of 50mM Tris-HCl, 2% SDS, 10% glycerol, and 0.05% bromophenol blue) was added to the samples, which were analyzed by immunoblotting.

### Immunoprecipitations

Cells were harvested at OD600 = 0.6 - 0.8 and rapidly frozen in liquid N_2_. Proteins were extracted by glass bead lysis in Buffer A (50mM Tris-HCl, 15mM EDTA, 15mM EGTA, 150mM NaCl, 0.1% Triton X100 complemented with 1mM benzamidine, 1mM PMSF, 100μg leupeptin and 100μg pepstatin). N-ethylmaleimide, S-methyl methanethiosulfonate or iodoacetamide were used as thiol blocking agents at a concentration of 20mM during the lysis for non-reducing extraction. For reducing extraction 20 mM DTT was used. Lysates were diluted 1:10 with Buffer A and immunoprecipitation resin (beads) was added for 1 h on a rotating wheel at 4°C. After incubation with the sample, beads were washed by sequentially increasing the NaCl concentration (0.15/0.3/0.6/0.8 M) in Buffer A. For western blot analysis, beads were resuspended in sample loading buffer, boiled at 95°C for 2’ and loaded. Immunoprecipitation resin (beads) used in this study: rabbit IgG-Agarose (0.12μl slurry/ml of extract), V5-Trap™ Magnetic Agarose (0.24μl/ml of extract), and HA-magnetic beads PierceTM (0.4μl/ml of extract). All beads were blocked in BSA (5μl of 2mg/ml stock in 1ml) supplemented 1xTBST for 30’ at 4°C. Before use, beads were equilibrated by washing 3x the beads with 500μl extraction buffer.

### Tandem affinity purification

Strains for the tandem affinity purification are based on the Yeast TAP Tagged ORFs Collection (Ghaemmaghami et al., 2003). 1000 ml of yeast culture (OD_60_0 = 0.8) was harvested per replicate and frozen in liquid N_2_. Subsequently, the pellet was resuspended in 24 ml of Buffer A and 4 ml of resuspension was transferred to a 50-ml vial containing 2 ml of glass beads. Cells were lysed by vortexing 10 x 1 min with 1 min rest on ice between vortexing steps. Samples were cleared by centrifuging at 5000 rpm for 15 min (Allegra® X-15R, Beckman-Coulter). Cleared lysates were pooled and 150μl of rabbit IgG-Agarose was added and incubated for 1h. Samples were further processed following the TAP purification protocol of the Gingras laboratory ^63^.

### Mass Spectrometry

Sample preparation. Samples were digested with endoproteinase LysC (1:10 w:w, 37°C, ON), and trypsin (1:10 w:w, 37°C, 8h). After digestion, peptide mix was acidified with formic acid and desalted with a MicroSpin C18 column (The Nest Group, Inc) prior to LC-MS/MS analysis.

Chromatographic and mass spectrometric analysis. i) TSA1^WT^ and TSA1^C171S^ experiments. Samples were analyzed using a LTQ-Orbitrap Velos Pro mass spectrometer (Thermo Fisher Scientific, San Jose, CA, USA) coupled to an EASY-nLC 1000 (Thermo Fisher Scientific (Proxeon), Odense, Denmark). Peptides were loaded onto the 2-cm Nano Trap column with an inner diameter of 100 μm packed with C18 particles of 5 μm particle size (Thermo Fisher Scientific) and were separated by reversed-phase chromatography using a 25-cm column with an inner diameter of 75 μm, packed with 1.9 μm C18 particles (Nikkyo Technos Co., Ltd. Japan). Chromatographic gradients started at 93% buffer A and 7% buffer B with a flow rate of 250 nl/min for 5 minutes and gradually increased 65% buffer A and 35% buffer B in 60 min. After each analysis, the column was washed for 15 min with 10% buffer A and 90% buffer B. Buffer A: 0.1% formic acid in water. Buffer B: 0.1% formic acid in acetonitrile.

The mass spectrometer was operated in positive ionization mode with nanospray voltage set at 2.1 kV and source temperature at 300°C. Ultramark 1621 for the was used for external calibration of the FT mass analyzer prior the analyses, and an internal calibration was performed using the background polysiloxane ion signal at m/z 445.1200. The instrument was operated in data-dependent acquisition (DDA) mode and full MS scans with 1 micro scans at resolution of 60,000 were used over a mass range of m/z 350-2000 with detection in the Orbitrap. Auto gain control (AGC) was set to 1E6, dynamic exclusion (60 seconds) and charge state filtering disqualifying singly charged peptides was activated. In each cycle of DDA analysis, following each survey scan, the top twenty most intense ions with multiple charged ions above a threshold ion count of 5000 were selected for fragmentation. Fragment ion spectra were produced via collision-induced dissociation (CID) at normalized collision energy of 35% and they were acquired in the ion trap mass analyzer. AGC was set to 1E4, isolation window of 2.0 m/z, an activation time of 10 ms and a maximum injection time of 100 ms were used. All data were acquired with Xcalibur software. Digested bovine serum albumin was analyzed between each sample to avoid sample carryover and to assure stability of the instrument. Acquired spectra were analyzed using the Proteome Discoverer software suite (v1.4, Thermo Fisher Scientific) and the Mascot search engine (v2.5, Matrix Science). The data were searched against a Swiss-Prot yeast database plus a list of common contaminants and all the corresponding decoy entries. For peptide identification the precursor ion mass tolerance was set to 7 ppm, trypsin was chosen as enzyme and up to three missed cleavages were allowed. The fragment ion mass tolerance was set to 0.5 Da. Oxidation of methionine and N-terminal protein acetylation were used as variable modifications whereas dithiomethane was set as variable modification on cysteines. False discovery rate (FDR) in peptide identification was set to a maximum of 5%.

ii) Dual IAM-NEM experiments. Samples were analyzed using a Orbitrap Eclipse mass spectrometer (Thermo Fisher Scientific, San Jose, CA, USA) coupled to an EASY-nLC 1200 (Thermo Fisher Scientific (Proxeon), Odense, Denmark). Peptides were loaded directly onto the analytical column and were separated by reversed-phase chromatography using a 50-cm column with an inner diameter of 75 μm, packed with 2 μm C18 particles spectrometer (Thermo Scientific, San Jose, CA, USA).

Chromatographic gradients started at 95% buffer A and 5% buffer B with a flow rate of 300 nl/min and gradually increased to 25% buffer B and 75% A in 79 min and then to 40% buffer B and 60% A in 11 min. After each analysis, the column was washed for 10 min with 100% buffer B. Buffer A: 0.1% formic acid in water. Buffer B: 0.1% formic acid in 80% acetonitrile.

The mass spectrometer was operated in positive ionization mode with nanospray voltage set at 2.4 kV and source temperature at 305°C. The acquisition was performed in data-dependent adquisition (DDA) mode and full MS scans with 1 micro scans at resolution of 120,000 were used over a mass range of m/z 350-1400 with detection in the Orbitrap mass analyzer. Auto gain control (AGC) was set to 4E5 and injection time to 50ms. In each cycle of data-dependent acquisition analysis, following each survey scan, the most intense ions above a threshold ion count of 10000 were selected for fragmentation. The number of selected precursor ions for fragmentation was determined by the “Top Speed” acquisition algorithm and a dynamic exclusion of 60 seconds. Fragment ion spectra were produced via high-energy collision dissociation (HCD) at normalized collision energy of 28% and they were acquired in the ion trap mass analyzer. AGC was set to 1E5, and an isolation window of 1.4 m/z and a maximum injection time of 200 ms were used.

Digested bovine serum albumin was analyzed between each sample to avoid sample carryover and to assure stability of the instrument. Acquired spectra were analyzed using the Proteome Discoverer software suite (v2.0, Thermo Fisher Scientific) and the Mascot search engine (v2.6, Matrix Science (4)). The data were searched against a Swiss-Prot Yeast database plus a list of common contaminants and all the corresponding decoy entries. For peptide identification a precursor ion mass tolerance of 7 ppm was used for MS1 level, trypsin was chosen as enzyme, and up to three missed cleavages were allowed. The fragment ion mass tolerance was set to 0.5 Da for MS2 spectra. Oxidation of methionine and N-terminal protein acetylation were used as variable modifications whereas carbamidomethylation and N-ethyl maleimide on cysteines was set as a fixed modification. False discovery rate (FDR) in peptide identification was set to a maximum of 5%.

The raw proteomics data have been deposited to the PRIDE repository with the dataset identifier PXD034411 ^64^.

### Immunoblotting

Protein lysates for the evaluation of TIMDIs were prepared in lysis buffer (50 mM Tris-HCl, 2% SDS, 10% glycerol, and 0.05% bromophenol blue) containing 2 mM S-Methyl methanethiosulfonate or 2 mM 1,4-dithiothreitol. Proteins were separated by dual Tris-glycine SDS-PAGE (lower quarter 15%, upper ¾ 10% acrylamide gel and transferred to a P-type PVDF membrane using the BioRad wet transfer system. The following primary antibodies were used for western blotting: anti-HA (1:100), anti-V5 (1:2000), anti-G6PDH (1:10’000), and anti-Peroxiredoxin-SO_2/3_ (1:2000). Anti-mouse IgG HRP-linked whole antibody (1:10’000) and anti-rabbit IgG from donkey whole antibody HRP (1:10’000) were used as secondary antibodies. Membranes were imaged using the Clarity Western ECL substrate on medical X-ray films using an automated developing machine (Hyperprocessor, Amersham Pharmacia).

For infrared fluorescence detection using the LI-COR Odyssey^®^ Infrared Imaging System SDS-PAGE gel was transferred to a LF-type PVDF membrane and blocked with Intercept® blocking buffer. Primary antibodies and secondary antibodies were also diluted in the same blocking buffer.

### 2D SDS-PAGE

For the 1^st^ dimension, samples were separated on a 1-mm SDS-PAGE. The cut gel piece was incubated in 50 mM Tris-HCl 2% SDS with or without 5 mM DTT at 45°C for 10 min and rinsed in stacking gel mix before being placed horizontally on a 1.5-mm SDS-PAGE. The gel piece was overlaid with a stacking gel mix and electrophoresis was performed.

### Image analysis

Films were scanned in TIF format at a resolution of 300 dpi using a (EPSON PERFECTION 4990 PHOTO, EPSON). Quantification for the TIMDI heat map was done using the FIJI software ^65^.

### Metabolomics Studies

Metabolite extraction, sample measurement and data processing for untargeted metabolomics was done according to Raguz Nakic et al., 2016. In brief, strains were grown as quadruplicates in Verduyn media supplemented with 2 g/l glucose as carbon source in deep well plates. At OD595 0.60 +/-0.08, metabolites were extracted using a hot extraction protocol.

Ions within a mass/charge ratio range of 50–1000 were measured by direct flow double injection of extracts on an Agilent 6550 series quadrupole TOF MS with the aid of a GERSTEL MPS2 autosampler. Ion annotation was performed using the S. cerevisiae reactants defined in the KEGG database. All further analysis with ions corresponding to deprotonated metabolites was conducted using Matlab (The Mathworks, Natick).

### 6-phosphogluconate dehydrogenase activity measurements

Cells were grown to an OD600 = 0.5, and 1 ml/condition was harvested, washed once in 1xPBS and then frozen in liquid N_2_ until processing. Sample extraction and 6-PGDH activity measurements were performed using the 6-Phosphogluconate Dehydrogenase Activity Colorimetric Assay Kit (BioVision, K540), following the manufactureŕs protocol.

### Time lapse microscopy

Cells were grown for 12h to reach an OD_600_=0.4 in low fluorescence media (YNB, CSM-URA supplemented with 2% glucose and 20μg/ml uracil). 400μl of cells were seeded into 8 well microcopy chamber slide, previously coated with concanavalin A (1mg/ml in dH_2_O, wells were incubated with 300μl for 30’, washed twice with dH_2_O and air dried). Cells were allowed to settle for 30’, supernatant was removed and 250ul low fluorescence media was added. Cycloheximide and/or AZC were added in 100μl media to achieve a final concentration of 100μg/ml and 5mM, respectively.

Images were acquired at 60x magnification (Plan Apo VC 60x Oil objective) using a Nikon Eclipse T*i* inverted microscope and an ORCA digital camera (Hamamatsu) using the NIS elements AR software. CoolLED pE excitation system was used and GFP wide field fluorescence and bright field channels were used.

### Reagents and materials

ssDNA oligos, reagents and materials used in this study can be found in the Supplementary Data 6 and Supplementary Data 8, respectively.

## Data availability

S. cerevisiae strains and plasmids generated in this manuscript are available upon request. Sequences of ssDNA oligos used in this study are available in Document S7.

The raw proteomics data have been deposited to the PRIDE repository. Data are available via ProteomeXchange with identifier PXD034411.

Any additional information required to reanalyze the data reported in this paper is available from the corresponding author upon request.

## Code availability

All packages used for data analysis are publicly available. No custom code was generated for this study.

## Supporting information

Description of supplementary files

Supplementary Data 1 - Mass Spectrometry Data Tsa1^WT and Tsa1^C171S

Supplementary Data 2 - Cysteine Frequency and Distance and PANTHER GO enrichment for Tsa1-WT and Tsa1-C171S interactors

Supplementary Data 3 - Mass Spectrometry Data DTT-eluted interactors of Tsa1^C48S and Tsa1^C171S

Supplementary Data 4 - Cysteine frequency and distance and PANTHER GO enrichment for DTT-eluted interactors of Tsa1^C48S and Tsa1^C171S

Supplementary Data 5 - Metabolomics Data

Supplementary Data 6 - Sequence of ssDNA oligos used in this study

Supplementary Data 7 - S cerevisiae strains used in this study

Supplementary Data 8 - Materials and Reagents used in this study

## Acknowledgements

We thank Dr. P Latorre (IRB Barcelona) for help with some of initial statistical analyses and C. Stephan-Otto Attolini and A. Caballe of the IRB Biostatistics Core Facility for their support with the statistical analysis of data. We are grateful to Prof. Shusuke Kuge, Faculty of Pharmaceutical Sciences, Tohoku Medical and Pharmaceutical University, Sendai, Japan, for sharing antisera to detect endogenous Tsa1 and Cdc19. The laboratories of FP and EdeN are supported by a coordinated grant from the Ministry of Science, Innovation, and Universities (PID2021-124723NB-C21/C22 and FEDER), and the Government of Catalonia (2017 SGR 799). We gratefully acknowledge institutional funding from the Ministry of Science, Innovation and Universities through the Centres of Excellence Severo Ochoa Award, and from the CERCA Programme of the Government of Catalonia and the Unidad de Excelencia María de Maeztu, funded by the AEI (CEX2018-000792-M). FP and EdeN are recipients of ICREA Acadèmia awards (Government of Catalonia). GS was supported by an Advanced Postdoc. Mobility fellowship (P300P3_147895) by the Swiss National Science Foundation.

## Statistical information

Growth curves and enzyme kinetics were performed at least in triplicates (number of replicates is indicated in the figure, where n represents the number of biological replicates) and curves are depicted as median values ± standard deviation as indicated in the figure legends. In graphs of experiments with a replicate number of three, all data points are depicted in the figure.

Heat map for relative TIMDI intensities was generated from western blots. To this aim, a linear selection starting at the highest molecular weight to the monomeric (in case of Gnd1) and dimeric (in case of Tsa1) protein form was used. Magnitude of intensity of pixels along this line was recorded. Experiments were performed in triplicates. From the same selection differential intensity analysis between stress conditions and the basal conditions was performed. Data was log normalized before fitting a linear model for each row, considering the experimental condition in the model. The “lmFit” function from the limma package was used to estimate the model coefficients, followed by the “contrasts.fit” function to consider the comparisons of interest and the “eBayes” function to estimate statistical significance ^67^. P-value adjustment was done with the Benjamini-Hochberg method. Adjusted p-value lower than 0.05 was considered to for significant differences.

Growth curve parameters were estimated by fitting a logistic curve to the observed data of independent experiments using the function “SummarizeGrowth” of the growthcurver R package ^68^. In order to compare conditions, a linear model was fitted to the parameters of the curves considering the experimental condition in the model. The “glht” function from the multcomp R package and confint function were used to estimate the coefficients and significance of the contrasts of interest ^69^.

For enzyme activity curves comparison, background levels were subtracted from observed activity levels sample wise. Resulting values were further normalized by dividing by total protein levels. A linear model was fit to the normalized values including the combination of time and condition as covariate. The “glht” function was used to compare conditions at each time point. Furthermore, the slopes of the linear regime were estimated through a linear model including the interaction between the condition and the time variable. The experimental batch was also included to consider observed technical biases. An ANOVA analysis was performed to compare conditions. Finally, the “lstrends” function from the lsmeans R package was used to estimate the significance of the differences in slopes ^70^. When applicable, batch adjusted values were computed by substracting the corresponding coefficient estimated by the model.

In the metabolomics experiment, the ion-wise fold change was determined for each mutant against the wild type and a cutoff on the fold change was applied for removal of very small changes. For this purpose, the absolute log2 fold changes retrieved from replicate permutation of all wild-type ions were pooled, and the 99% quantile was determined to be at 0.3616. Accordingly, corresponding to a 30% change, the log2 fold change cutoff was defined at 0.3785. P-values were calculated by a two-sided 2-sample t-test assuming unequal variance and corrected for multiple testing as previously described ^71^. Ions with an absolute log2 fold change > 0.3785 and a corrected p-value < 10^-3^ were considered significantly changing.

## Author contributions

GS and ZR performed the experiments; EB, ES performed proteomic analysis; GS, US, EdeN, and FP participated in the design, data analysis and writing of the manuscript.

