## Supplementary material for "Peroxiredoxinylation buffers the redox state of the proteome upon cellular stress": Description of supplementary files

**Description of additional Supplementary Files**

**Supplementary Data 1**

Mass spectrometry Data of TAP-immunopurification of Tsa1WT-TAP and Tsa1C171S-TAP. Contains the search parameters, identified protein groups and peptides.

**Supplementary Data 2**

Cysteine Frequency and Distance and PANTHER GO enrichment for Tsa1^WT^-TAP and Tsa1^C171S^-TAP interactors. Contains the cysteine frequency and distance measurements. Contains the GO term annotation and enrichment analysis.

**Supplementary Data 3**

Mass spectrometry Data of HA-immunopurification of Tsa1^C48S^-HA and Tsa1^C171S^-HA. Contains the search parameters, identified protein groups and peptides.

**Supplementary Data 4**

Cysteine Frequency and Distance and PANTHER GO enrichment for Tsa1^C48S^-HA and Tsa1^C171S^-HA interactors. Contains the cysteine frequency and distance measurements. Contains the GO term annotation and enrichment analysis.

**Supplementary Data 5**

Untargeted metabolomics dataset of Tsa1^WT^-HA, Tsa1^C48S^-HA and Tsa1^C171S^-HA. Contains the full annotation of the metabolites and detected metabolites.

**Supplementary Data 6**

Sequence of ssDNA oligos used in this study.

**Supplementary Data 7**

*S. cerevisiae* strains used in this study.

**Supplementary Data 8**

Materials and Reagents used in this study.
