## Supplementary Figures for "Peroxiredoxinylation buffers the redox state of the proteome upon cellular stress"

### Slide 1
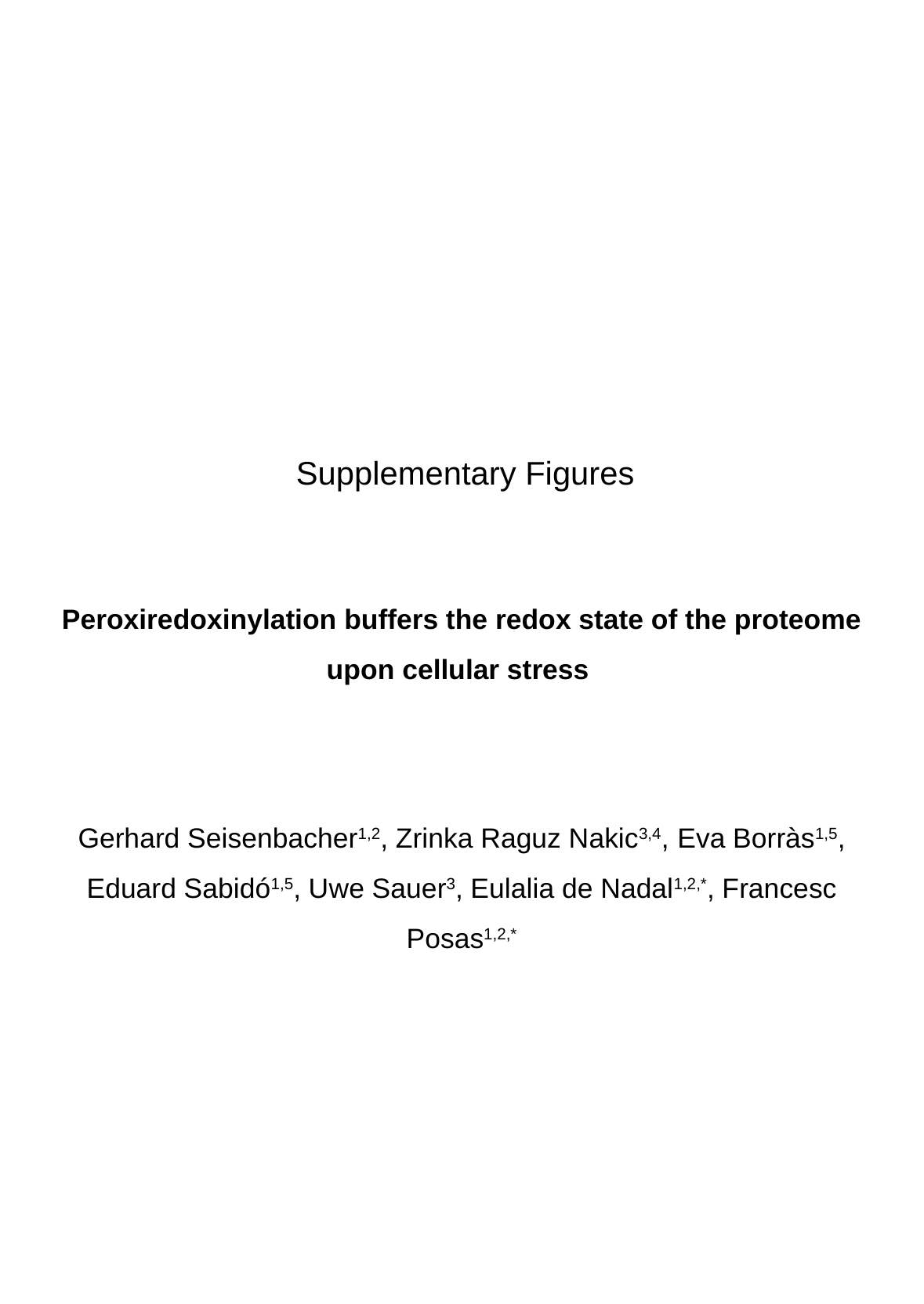

Supplementary Figures
Peroxiredoxinylation buffers the redox state of the proteome upon cellular stress
Gerhard Seisenbacher^1,2^, Zrinka Raguz Nakic^3,4^, Eva Borràs^1,5^, Eduard Sabidó^1,5^, Uwe Sauer^3^, Eulalia de Nadal^1,2,*^, Francesc Posas^1,2,*^

### Slide 2
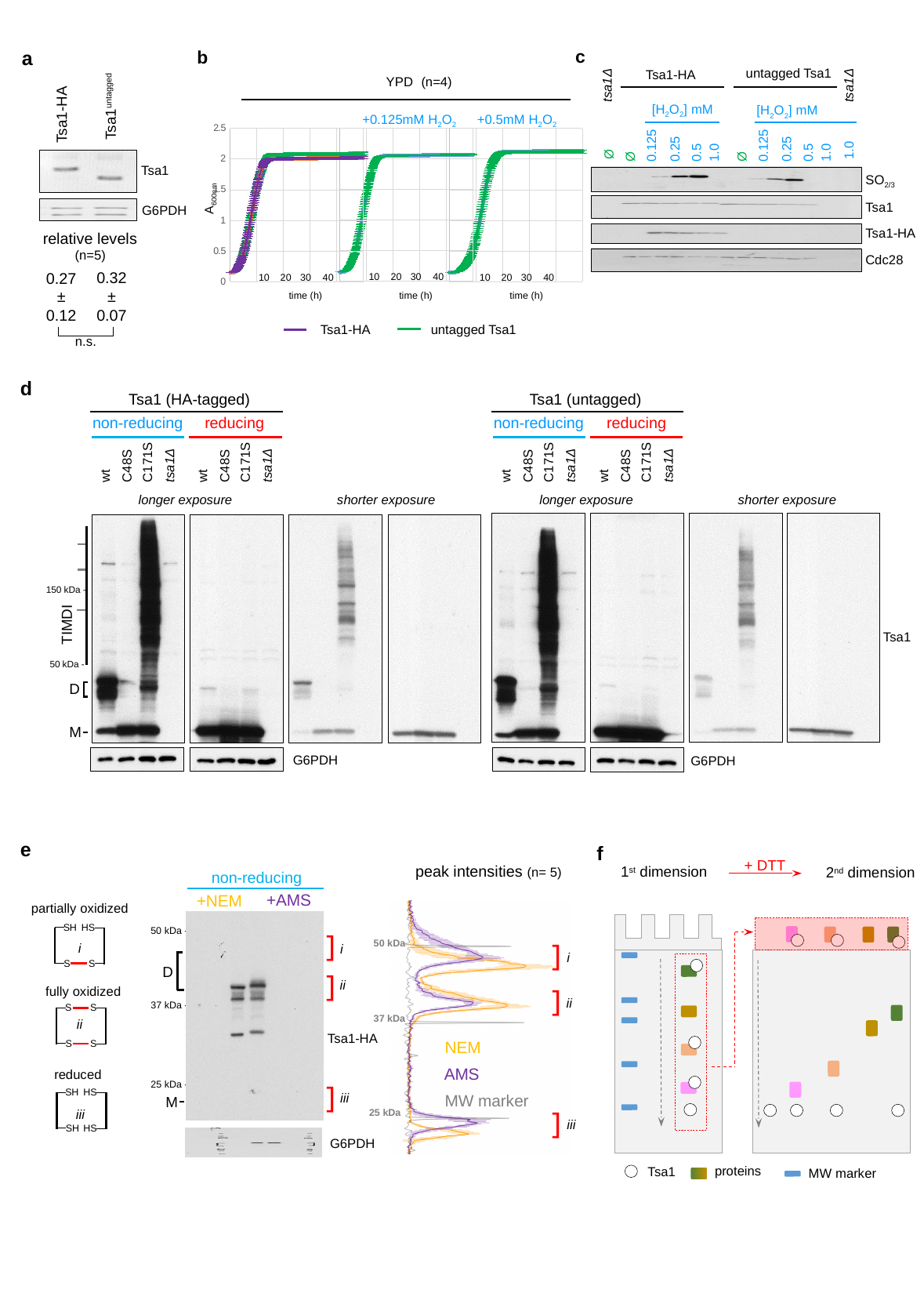

c
b
a
Tsa1^untagged^
Tsa1-HA
Tsa1
G6PDH
relative levels
(n=5)
0.32
±
0.07
0.27
±
0.12
n.s.
untagged Tsa1
Tsa1-HA
(n=4)
YPD
+0.125mM H_2_O_2_
+0.5mM H_2_O_2_
#### Chart
| Category | | |
|---|---|---|10
20
30
40
time (h)
#### Chart
| Category | | |
|---|---|---|10
20
30
40
time (h)
#### Chart
| Category | | |
|---|---|---|10
20
30
40
time (h)
A_600nm_
Tsa1-HA
untagged Tsa1
tsa1Δ
tsa1Δ
[H_2_O_2_] mM
[H_2_O_2_] mM
0.125
Ø
0.25
0.5
1.0
0.125
Ø
0.25
0.5
1.0
1.0
Ø
SO_2/3_
Tsa1
Tsa1-HA
Cdc28
d
Tsa1 (untagged)
non-reducing
C171S
tsa1Δ
C48S
wt
reducing
C171S
tsa1Δ
C48S
wt
longer exposure
shorter exposure
Tsa1 (HA-tagged)
non-reducing
C171S
tsa1Δ
C48S
wt
reducing
C171S
tsa1Δ
C48S
wt
longer exposure
shorter exposure
150 kDa -
50 kDa -
TIMDI
G6PDH
G6PDH
Tsa1
D
M
e
f
+ DTT
peak intensities (n= 5)
1^st^ dimension
2^nd^ dimension
non-reducing
+AMS
+NEM
50 kDa -
]
D
]
37 kDa -
25 kDa -
]
M
i
ii
Tsa1-HA
iii
G6PDH
partially oxidized
SH
SH
HS
HS
i
fully oxidized
SH
SH
HS
HS
ii
reduced
SH
SH
HS
HS
iii
50 kDa
]
i
]
ii
37 kDa
NEM
AMS
MW marker
]
25 kDa
iii
proteins
Tsa1
MW marker

### Slide 3
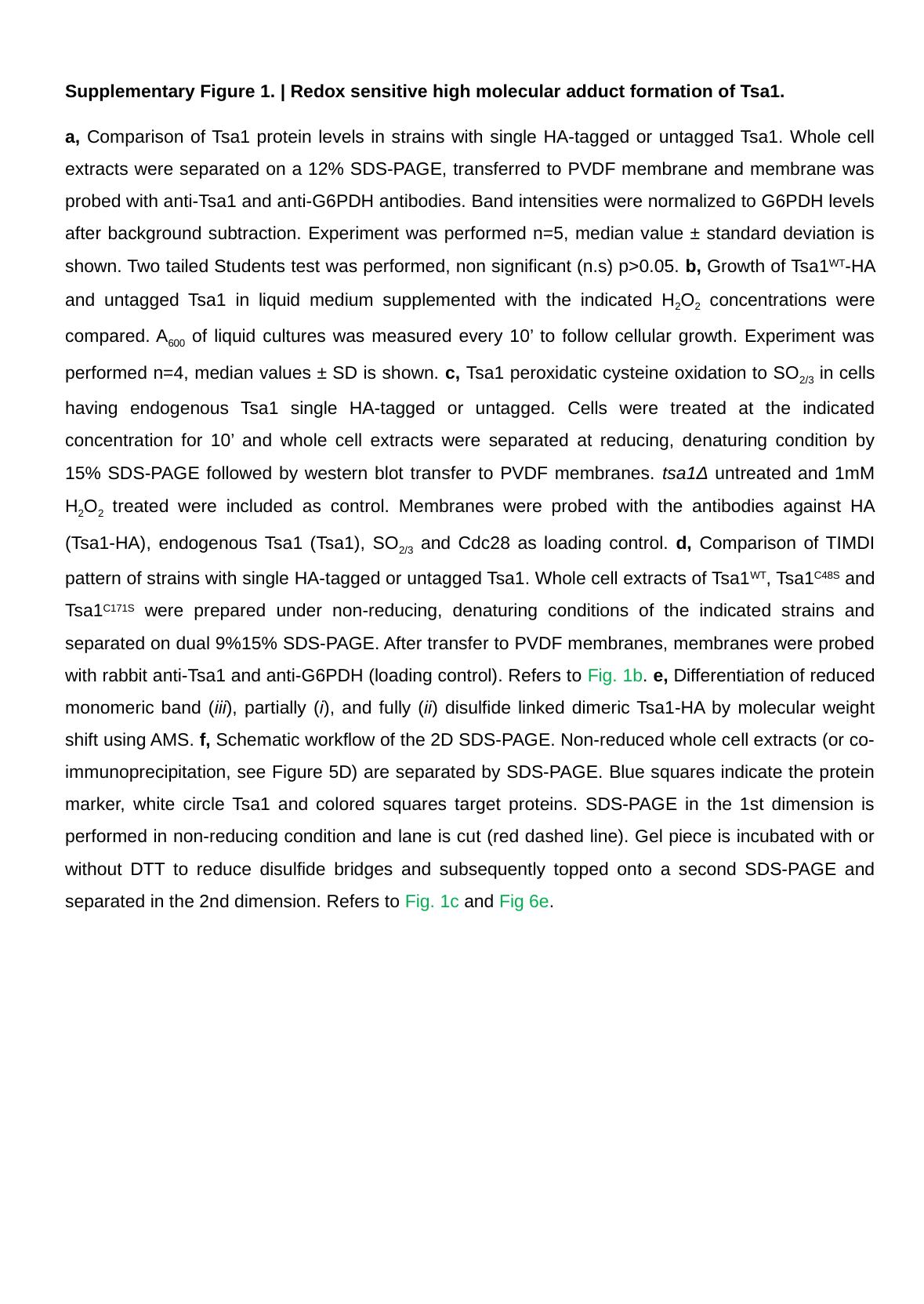

Supplementary Figure 1. | Redox sensitive high molecular adduct formation of Tsa1.
a, Comparison of Tsa1 protein levels in strains with single HA-tagged or untagged Tsa1. Whole cell extracts were separated on a 12% SDS-PAGE, transferred to PVDF membrane and membrane was probed with anti-Tsa1 and anti-G6PDH antibodies. Band intensities were normalized to G6PDH levels after background subtraction. Experiment was performed n=5, median value ± standard deviation is shown. Two tailed Students test was performed, non significant (n.s) p>0.05. b, Growth of Tsa1^WT^-HA and untagged Tsa1 in liquid medium supplemented with the indicated H_2_O_2_ concentrations were compared. A_600_ of liquid cultures was measured every 10’ to follow cellular growth. Experiment was performed n=4, median values ± SD is shown. c, Tsa1 peroxidatic cysteine oxidation to SO_2/3_ in cells having endogenous Tsa1 single HA-tagged or untagged. Cells were treated at the indicated concentration for 10’ and whole cell extracts were separated at reducing, denaturing condition by 15% SDS-PAGE followed by western blot transfer to PVDF membranes. tsa1Δ untreated and 1mM H_2_O_2_ treated were included as control. Membranes were probed with the antibodies against HA (Tsa1-HA), endogenous Tsa1 (Tsa1), SO_2/3_ and Cdc28 as loading control. d, Comparison of TIMDI pattern of strains with single HA-tagged or untagged Tsa1. Whole cell extracts of Tsa1^WT^, Tsa1^C48S^ and Tsa1^C171S^ were prepared under non-reducing, denaturing conditions of the indicated strains and separated on dual 9%15% SDS-PAGE. After transfer to PVDF membranes, membranes were probed with rabbit anti-Tsa1 and anti-G6PDH (loading control). Refers to Fig. 1b. e, Differentiation of reduced monomeric band (iii), partially (i), and fully (ii) disulfide linked dimeric Tsa1-HA by molecular weight shift using AMS. f, Schematic workflow of the 2D SDS-PAGE. Non-reduced whole cell extracts (or co-immunoprecipitation, see Figure 5D) are separated by SDS-PAGE. Blue squares indicate the protein marker, white circle Tsa1 and colored squares target proteins. SDS-PAGE in the 1st dimension is performed in non-reducing condition and lane is cut (red dashed line). Gel piece is incubated with or without DTT to reduce disulfide bridges and subsequently topped onto a second SDS-PAGE and separated in the 2nd dimension. Refers to Fig. 1c and Fig 6e.

### Slide 4
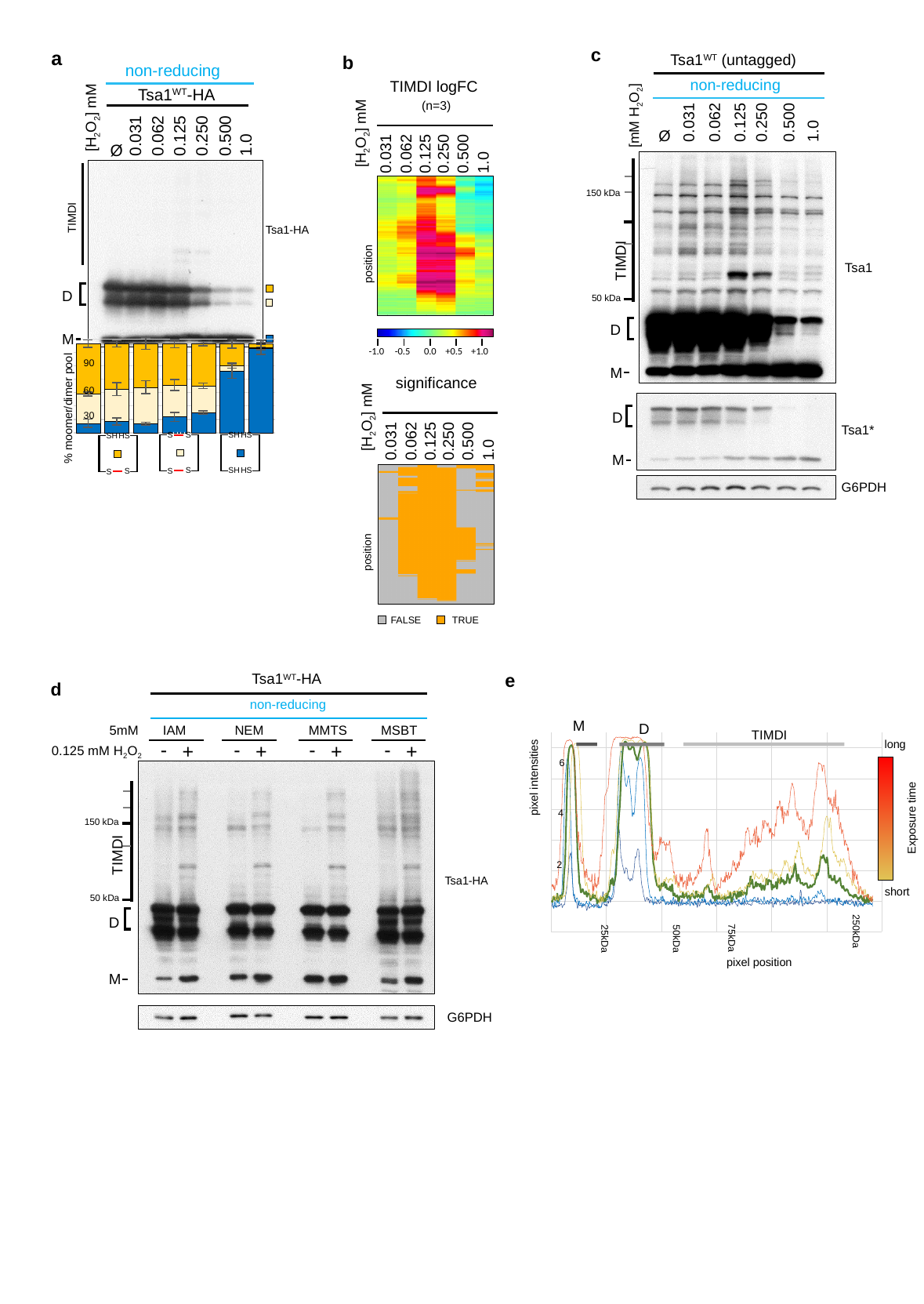

c
Tsa1^WT^ (untagged)
non-reducing
[mM H_2_O_2_]
0.062
0.125
0.250
0.500
1.0
0.031
Ø
150 kDa
50 kDa
TIMDI
Tsa1
Tsa1*
G6PDH
a
b
TIMDI logFC
 (n=3)
[H_2_O_2_] mM
0.062
0.125
0.250
0.500
0.031
1.0
position
-1.0
-0.5
0.0
+0.5
+1.0
non-reducing
[H_2_O_2_] mM
Ø
Tsa1^WT^-HA
0.031
0.062
0.125
0.250
0.500
1.0
TIMDI
Tsa1-HA
D
M
D
M
#### Chart
| Category | III | III | I |
|---|---|---|---|
| BG 0mM | 10.538446976725156 | 32.88394774404073 | 56.577605279234106 |
| BG 0.031mM | 12.566393359270686 | 36.26137601415085 | 51.172230626578454 |
| BG 0.062mM | 10.180641180508195 | 40.88662725650717 | 48.93273156298463 |
| BG 0.125mM | 17.766126085338488 | 35.722643038692794 | 46.511230875968735 |
| BG 0.25mM | 22.725095414004127 | 30.028765703633955 | 47.24613888236192 |
| BG 0.5mM | 69.16467248362112 | 6.431082441487384 | 24.40424507489151 |
| BG 1.0mM | 94.5973027835462 | 1.3486137047875448 | 4.0540835116662635 |
90
significance
[H_2_O_2_] mM
0.062
0.125
0.250
0.500
0.031
1.0
position
FALSE
TRUE
60
% moomer/dimer pool
D
M
30
HS
SH
HS
SH
HS
SH
HS
SH
HS
HS
SH
SH
e
TIMDI
#### Chart
| Category | | | | | | |
|---|---|---|---|---|---|---|long
6
pixel intensities
4
Exposure time
2
short
250kDa
75kDa
25kDa
50kDa
pixel position
Tsa1^WT^-HA
d
non-reducing
5mM
IAM
NEM
MMTS
MSBT
-
+
-
+
-
+
-
+
0.125 mM H_2_O_2_
150 kDa
50 kDa
M
D
TIMDI
Tsa1-HA
D
M
G6PDH

### Slide 5
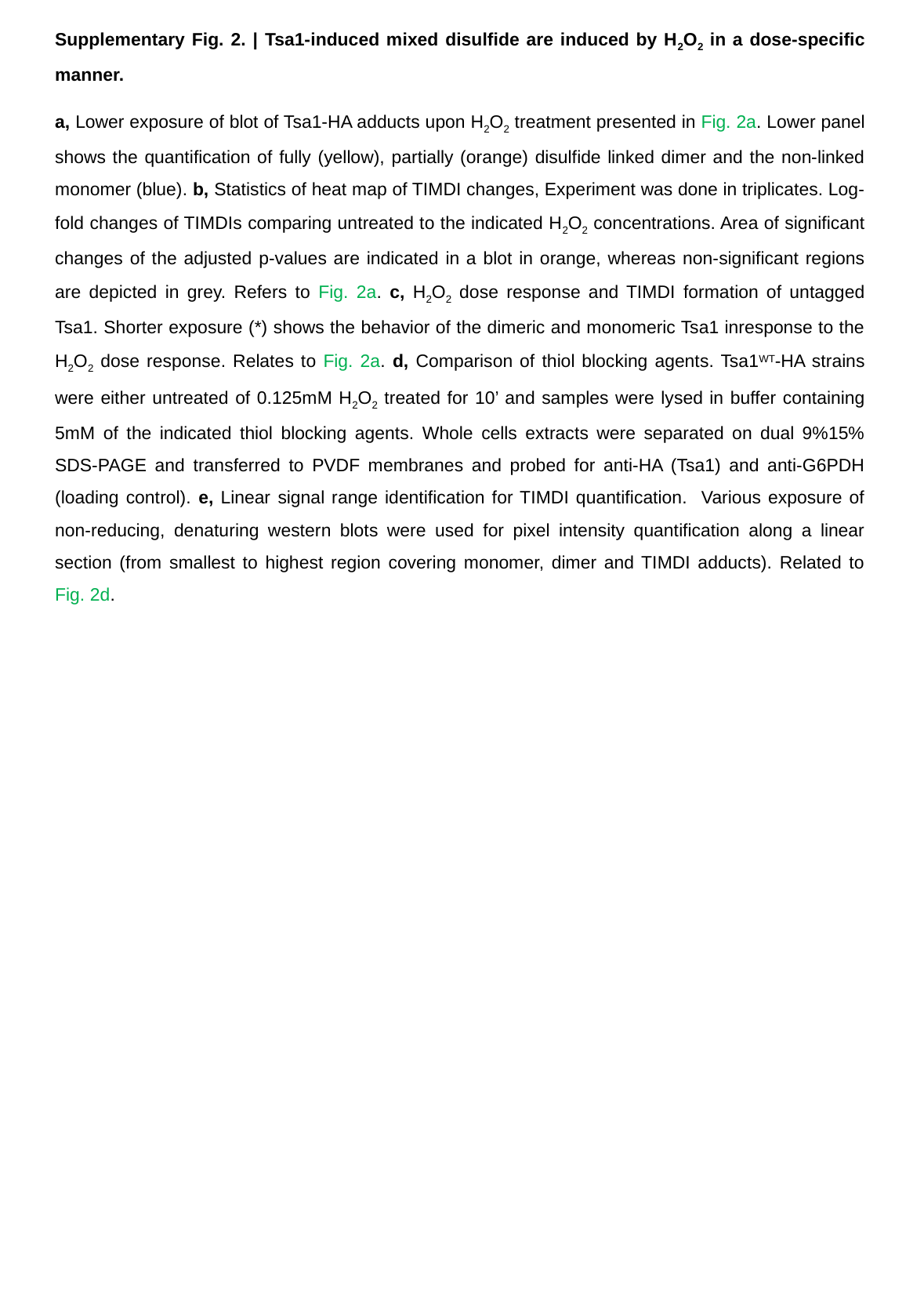

Supplementary Fig. 2. | Tsa1-induced mixed disulfide are induced by H_2_O_2_ in a dose-specific manner.
a, Lower exposure of blot of Tsa1-HA adducts upon H_2_O_2_ treatment presented in Fig. 2a. Lower panel shows the quantification of fully (yellow), partially (orange) disulfide linked dimer and the non-linked monomer (blue). b, Statistics of heat map of TIMDI changes, Experiment was done in triplicates. Log-fold changes of TIMDIs comparing untreated to the indicated H_2_O_2_ concentrations. Area of significant changes of the adjusted p-values are indicated in a blot in orange, whereas non-significant regions are depicted in grey. Refers to Fig. 2a. c, H_2_O_2_ dose response and TIMDI formation of untagged Tsa1. Shorter exposure (*) shows the behavior of the dimeric and monomeric Tsa1 inresponse to the H_2_O_2_ dose response. Relates to Fig. 2a. d, Comparison of thiol blocking agents. Tsa1^WT^-HA strains were either untreated of 0.125mM H_2_O_2_ treated for 10’ and samples were lysed in buffer containing 5mM of the indicated thiol blocking agents. Whole cells extracts were separated on dual 9%15% SDS-PAGE and transferred to PVDF membranes and probed for anti-HA (Tsa1) and anti-G6PDH (loading control). e, Linear signal range identification for TIMDI quantification. Various exposure of non-reducing, denaturing western blots were used for pixel intensity quantification along a linear section (from smallest to highest region covering monomer, dimer and TIMDI adducts). Related to Fig. 2d.

### Slide 6
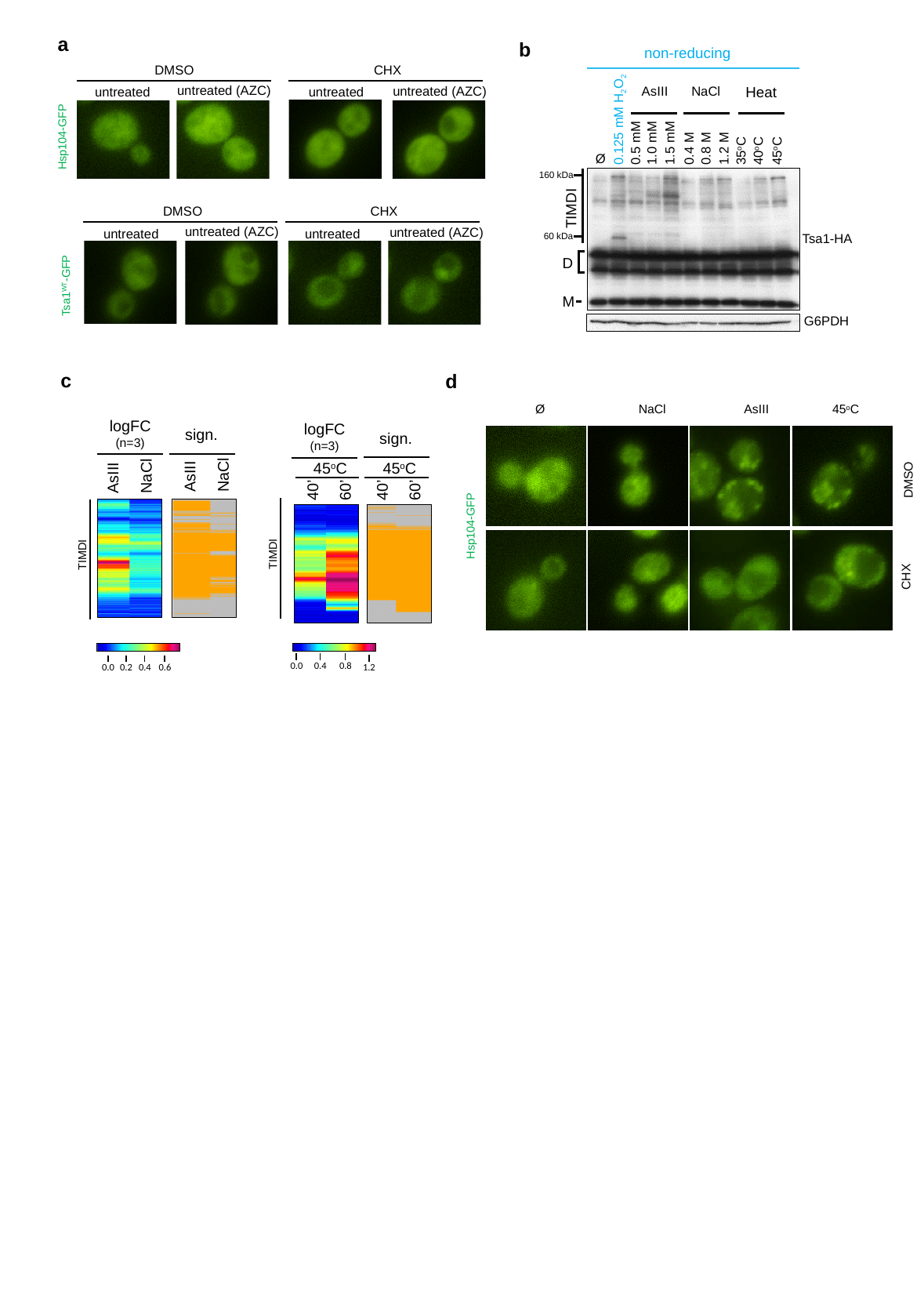

a
DMSO
CHX
untreated (AZC)
untreated
untreated
Hsp104-GFP
DMSO
CHX
untreated
untreated
Tsa1^WT^-GFP
b
0.125 mM H_2_O_2_
1.0 mM
0.5 mM
1.5 mM
0.4 M
0.8 M
1.2 M
35^o^C
40^o^C
45^o^C
Ø
non-reducing
AsIII
NaCl
Heat
160 kDa
TIMDI
60 kDa
Tsa1-HA
G6PDH
untreated (AZC)
untreated (AZC)
untreated (AZC)
D
M
c
d
Ø
45^o^C
NaCl
AsIII
logFC
(n=3)
logFC
(n=3)
sign.
45^o^C
40’
60’
45^o^C
40’
60’
sign.
NaCl
NaCl
AsIII
AsIII
DMSO
Hsp104-GFP
TIMDI
TIMDI
CHX
0.0
0.2
0.4
0.6
0.4
0.8
0.0
1.2

### Slide 7
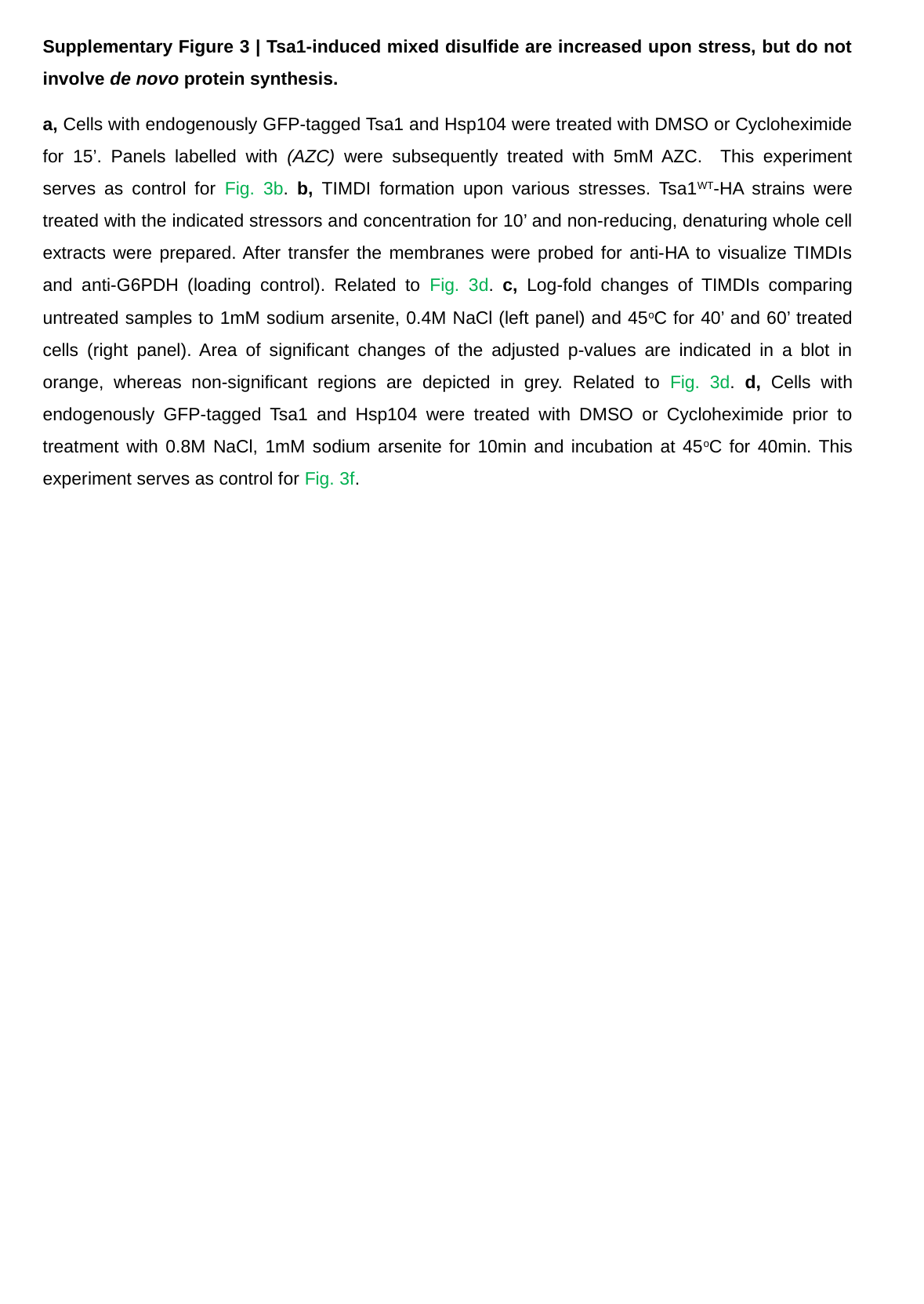

Supplementary Figure 3 | Tsa1-induced mixed disulfide are increased upon stress, but do not involve de novo protein synthesis.
a, Cells with endogenously GFP-tagged Tsa1 and Hsp104 were treated with DMSO or Cycloheximide for 15’. Panels labelled with (AZC) were subsequently treated with 5mM AZC. This experiment serves as control for Fig. 3b. b, TIMDI formation upon various stresses. Tsa1^WT^-HA strains were treated with the indicated stressors and concentration for 10’ and non-reducing, denaturing whole cell extracts were prepared. After transfer the membranes were probed for anti-HA to visualize TIMDIs and anti-G6PDH (loading control). Related to Fig. 3d. c, Log-fold changes of TIMDIs comparing untreated samples to 1mM sodium arsenite, 0.4M NaCl (left panel) and 45^o^C for 40’ and 60’ treated cells (right panel). Area of significant changes of the adjusted p-values are indicated in a blot in orange, whereas non-significant regions are depicted in grey. Related to Fig. 3d. d, Cells with endogenously GFP-tagged Tsa1 and Hsp104 were treated with DMSO or Cycloheximide prior to treatment with 0.8M NaCl, 1mM sodium arsenite for 10min and incubation at 45^o^C for 40min. This experiment serves as control for Fig. 3f.

### Slide 8
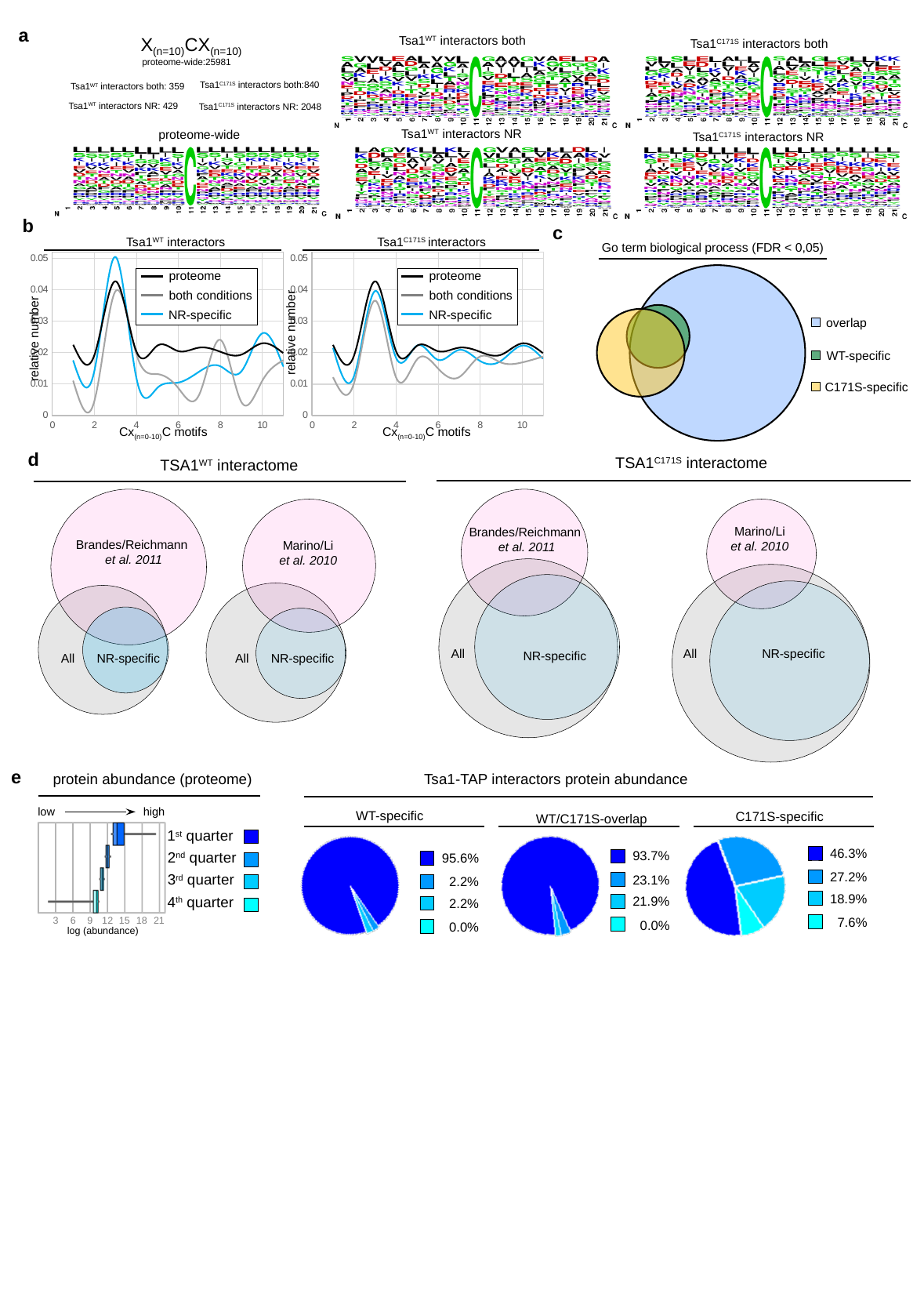

a
X_(n=10)_CX_(n=10)_
Tsa1^WT^ interactors both
Tsa1^C171S^ interactors both
proteome-wide:25981
Tsa1^C171S^ interactors both:840
Tsa1^WT^ interactors both: 359
Tsa1^WT^ interactors NR: 429
Tsa1^C171S^ interactors NR: 2048
Tsa1^WT^ interactors NR
proteome-wide
Tsa1^C171S^ interactors NR
b
Tsa1^WT^ interactors
Tsa1^C171S^ interactors
#### Chart
| Category |
|---|
#### Chart
| Category | | | |
|---|---|---|---|proteome
both conditions
NR-specific
proteome
both conditions
NR-specific
relative number
relative number
Cx_(n=0-10)_C motifs
Cx_(n=0-10)_C motifs
c
Go term biological process (FDR < 0,05)
overlap
WT-specific
C171S-specific
d
TSA1^C171S^ interactome
TSA1^WT^ interactome
Marino/Li
et al. 2010
Brandes/Reichmann
 et al. 2011
Brandes/Reichmann
 et al. 2011
Marino/Li
et al. 2010
All
All
NR-specific
NR-specific
All
NR-specific
All
NR-specific
e
protein abundance (proteome)
3
6
9
12
15
18
21
low
high
1^st^ quarter
2^nd^ quarter
3^rd^ quarter
4^th^ quarter
log (abundance)
Tsa1-TAP interactors protein abundance
WT-specific
C171S-specific
WT/C171S-overlap
46.3%
27.2%
18.9%
7.6%
93.7%
23.1%
21.9%
0.0%
95.6%
2.2%
2.2%
0.0%
#### Chart
| Category |
|---|

### Slide 9
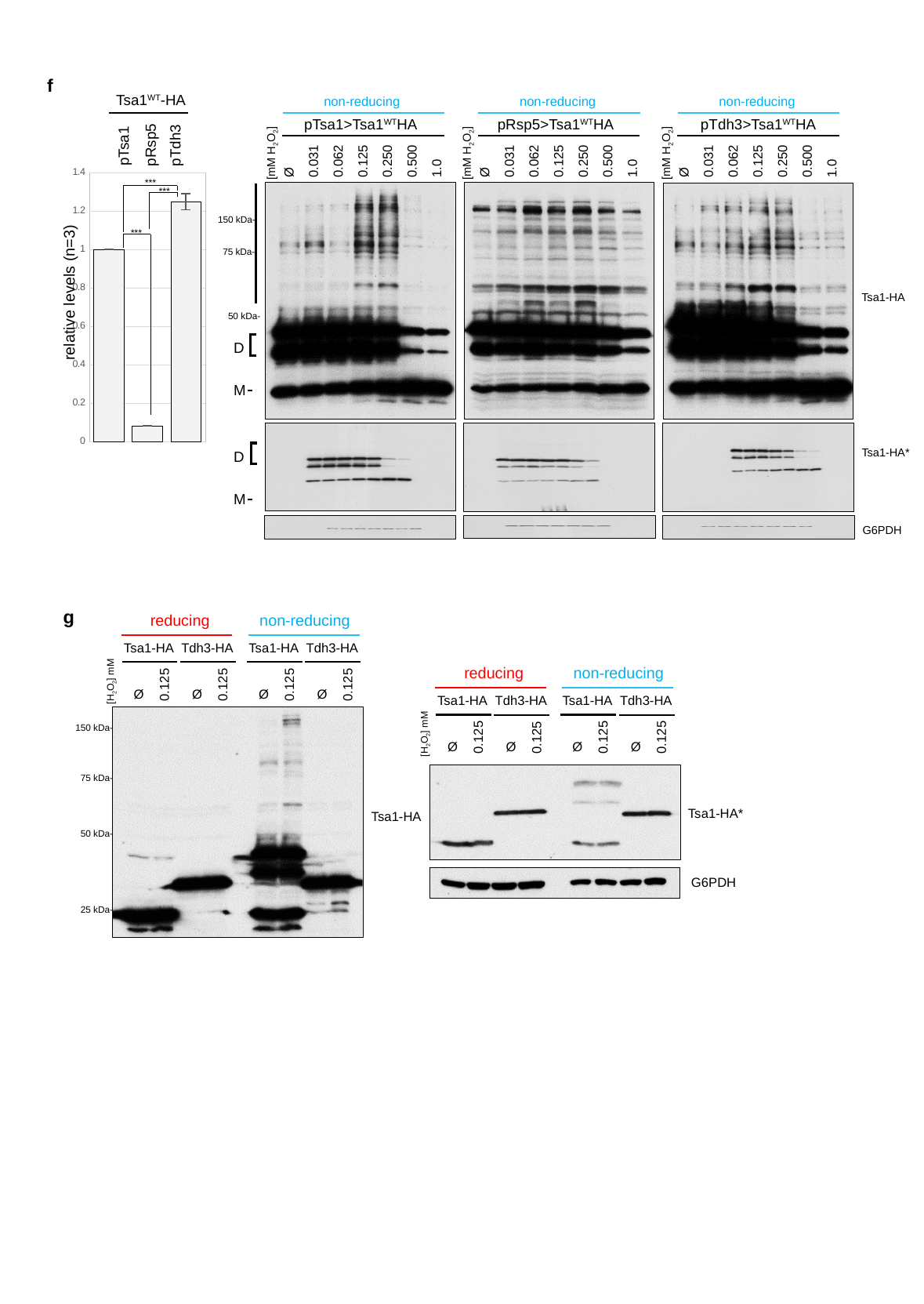

f
Tsa1^WT^-HA
non-reducing
non-reducing
pRsp5>Tsa1^WT^HA
0.031
0.062
0.125
0.250
[mM H_2_O_2_]
0.500
1.0
Ø
non-reducing
pTdh3>Tsa1^WT^HA
0.031
0.062
0.125
0.250
[mM H_2_O_2_]
0.500
1.0
Ø
Tsa1-HA
Tsa1-HA*
G6PDH
pTsa1>Tsa1^WT^HA
pRsp5
pTdh3
pTsa1
0.031
0.062
0.125
0.250
[mM H_2_O_2_]
0.500
1.0
Ø
#### Chart
| Category | |
|---|---|***
***
150 kDa-
***
75 kDa-
relative levels (n=3)
50 kDa-
D
M
D
M
g
reducing
non-reducing
Tsa1-HA
0.125
Ø
Tdh3-HA
0.125
Ø
Tsa1-HA
0.125
Ø
Tdh3-HA
0.125
Ø
reducing
non-reducing
[H_2_O_2_] mM
Tsa1-HA
0.125
Ø
Tdh3-HA
0.125
Ø
Tsa1-HA
0.125
Ø
Tdh3-HA
0.125
Ø
150 kDa-
[H_2_O_2_] mM
75 kDa-
Tsa1-HA*
Tsa1-HA
50 kDa-
25 kDa-
G6PDH

### Slide 10
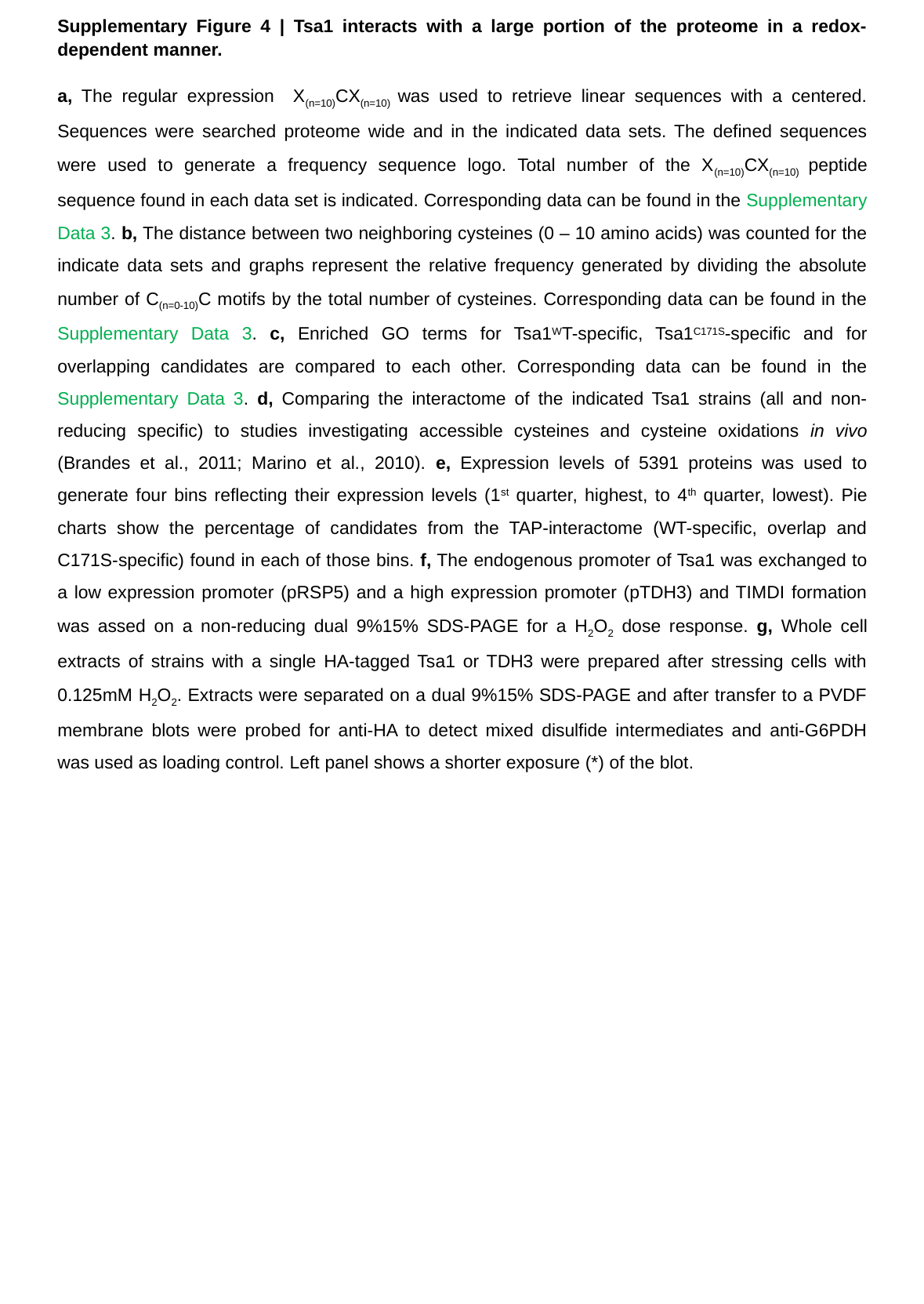

Supplementary Figure 4 | Tsa1 interacts with a large portion of the proteome in a redox-dependent manner.
a, The regular expression X_(n=10)_CX_(n=10)_ was used to retrieve linear sequences with a centered. Sequences were searched proteome wide and in the indicated data sets. The defined sequences were used to generate a frequency sequence logo. Total number of the X_(n=10)_CX_(n=10)_ peptide sequence found in each data set is indicated. Corresponding data can be found in the Supplementary Data 3. b, The distance between two neighboring cysteines (0 – 10 amino acids) was counted for the indicate data sets and graphs represent the relative frequency generated by dividing the absolute number of C_(n=0-10)_C motifs by the total number of cysteines. Corresponding data can be found in the Supplementary Data 3. c, Enriched GO terms for Tsa1^W^T-specific, Tsa1^C171S^-specific and for overlapping candidates are compared to each other. Corresponding data can be found in the Supplementary Data 3. d, Comparing the interactome of the indicated Tsa1 strains (all and non-reducing specific) to studies investigating accessible cysteines and cysteine oxidations in vivo (Brandes et al., 2011; Marino et al., 2010). e, Expression levels of 5391 proteins was used to generate four bins reflecting their expression levels (1^st^ quarter, highest, to 4^th^ quarter, lowest). Pie charts show the percentage of candidates from the TAP-interactome (WT-specific, overlap and C171S-specific) found in each of those bins. f, The endogenous promoter of Tsa1 was exchanged to a low expression promoter (pRSP5) and a high expression promoter (pTDH3) and TIMDI formation was assed on a non-reducing dual 9%15% SDS-PAGE for a H_2_O_2_ dose response. g, Whole cell extracts of strains with a single HA-tagged Tsa1 or TDH3 were prepared after stressing cells with 0.125mM H_2_O_2_. Extracts were separated on a dual 9%15% SDS-PAGE and after transfer to a PVDF membrane blots were probed for anti-HA to detect mixed disulfide intermediates and anti-G6PDH was used as loading control. Left panel shows a shorter exposure (*) of the blot.

### Slide 11
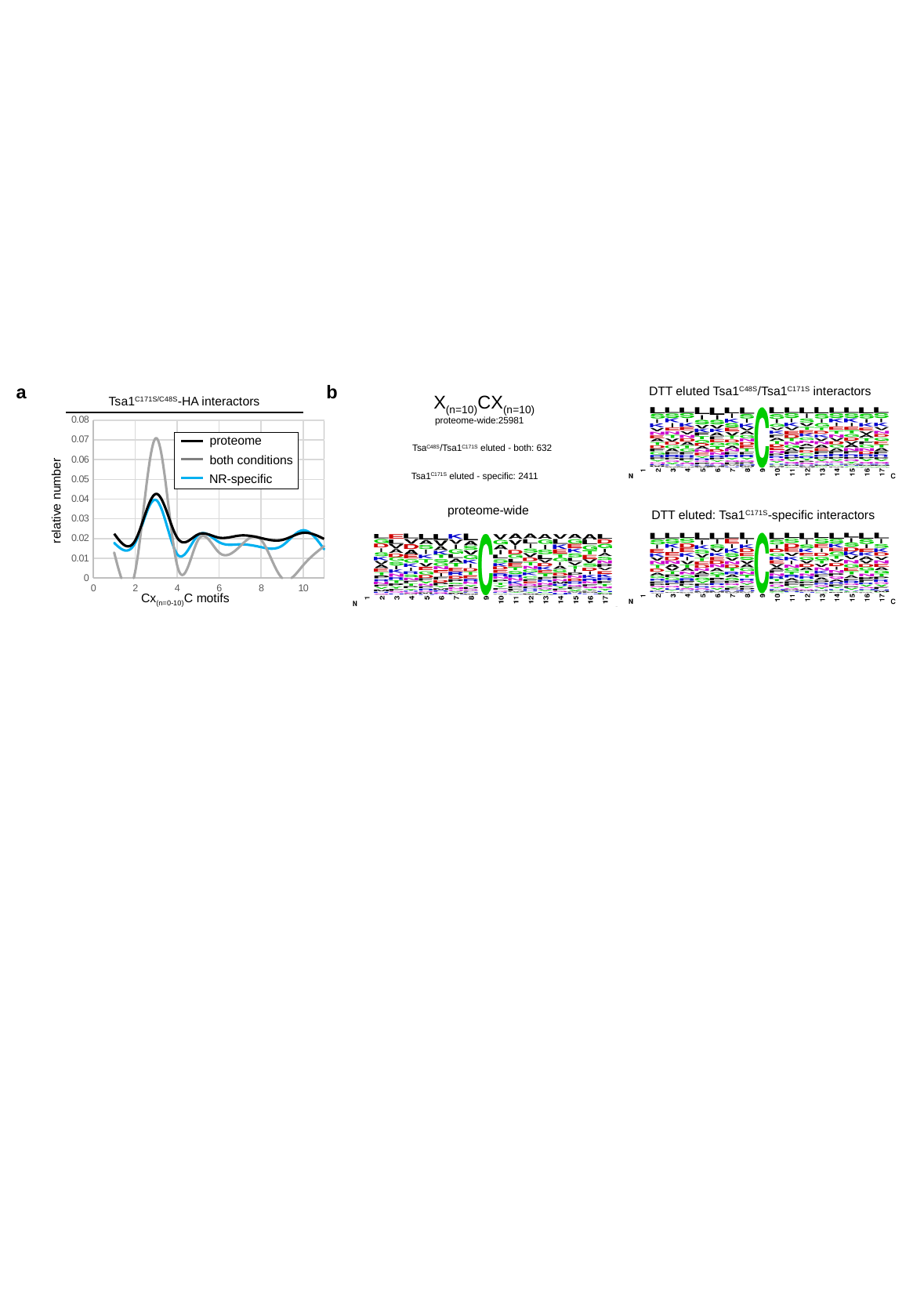

a
b
DTT eluted Tsa1^C48S^/Tsa1^C171S^ interactors
X_(n=10)_CX_(n=10)_
Tsa1^C171S/C48S^-HA interactors
proteome-wide:25981
#### Chart
| Category | | | |
|---|---|---|---|proteome
both conditions
NR-specific
Tsa^C48S^/Tsa1^C171S^ eluted - both: 632
Tsa1^C171S^ eluted - specific: 2411
relative number
proteome-wide
DTT eluted: Tsa1^C171S^-specific interactors
Cx_(n=0-10)_C motifs

### Slide 12
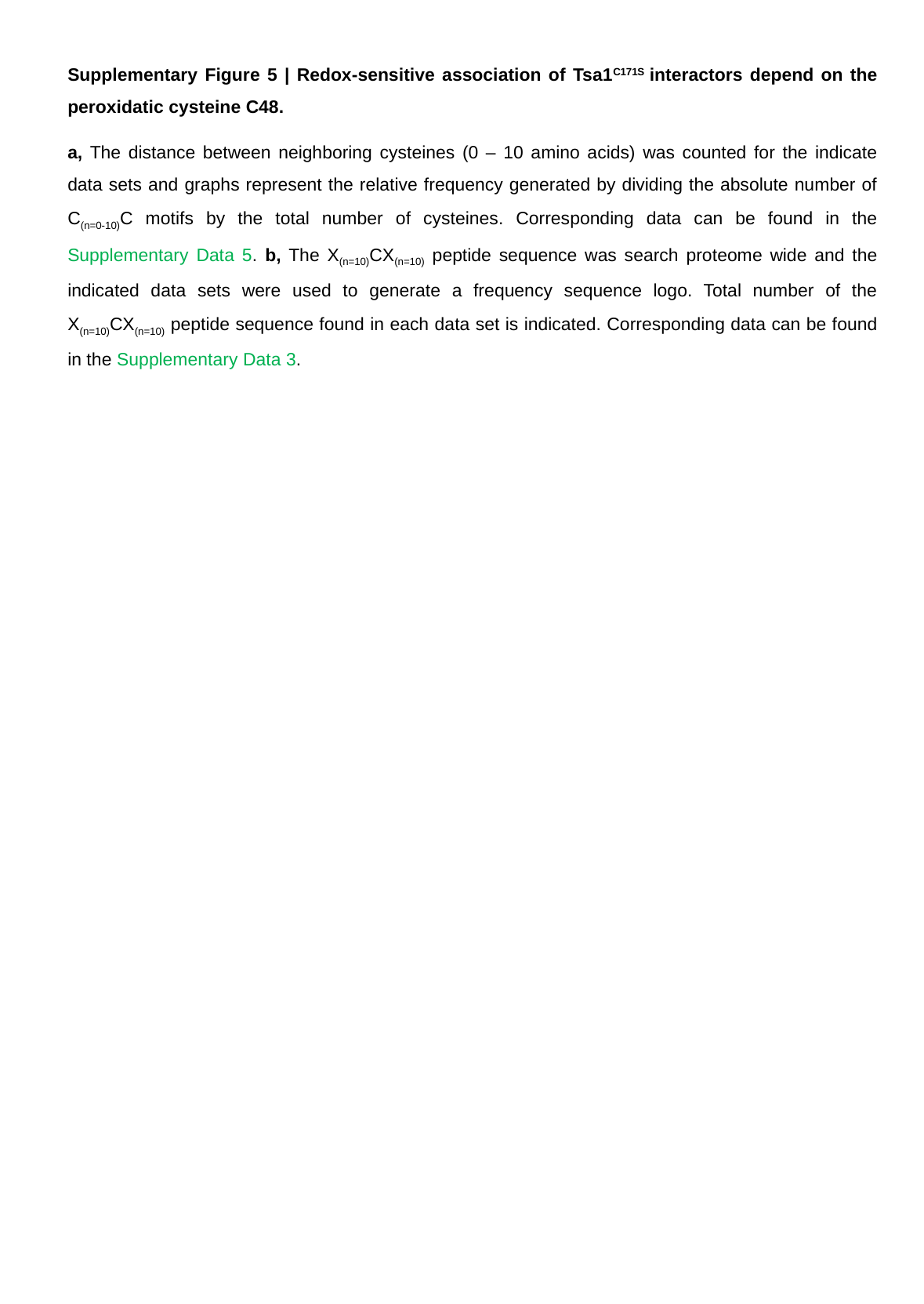

Supplementary Figure 5 | Redox-sensitive association of Tsa1^C171S^ interactors depend on the peroxidatic cysteine C48.
a, The distance between neighboring cysteines (0 – 10 amino acids) was counted for the indicate data sets and graphs represent the relative frequency generated by dividing the absolute number of C_(n=0-10)_C motifs by the total number of cysteines. Corresponding data can be found in the Supplementary Data 5. b, The X_(n=10)_CX_(n=10)_ peptide sequence was search proteome wide and the indicated data sets were used to generate a frequency sequence logo. Total number of the X_(n=10)_CX_(n=10)_ peptide sequence found in each data set is indicated. Corresponding data can be found in the Supplementary Data 3.

### Slide 13
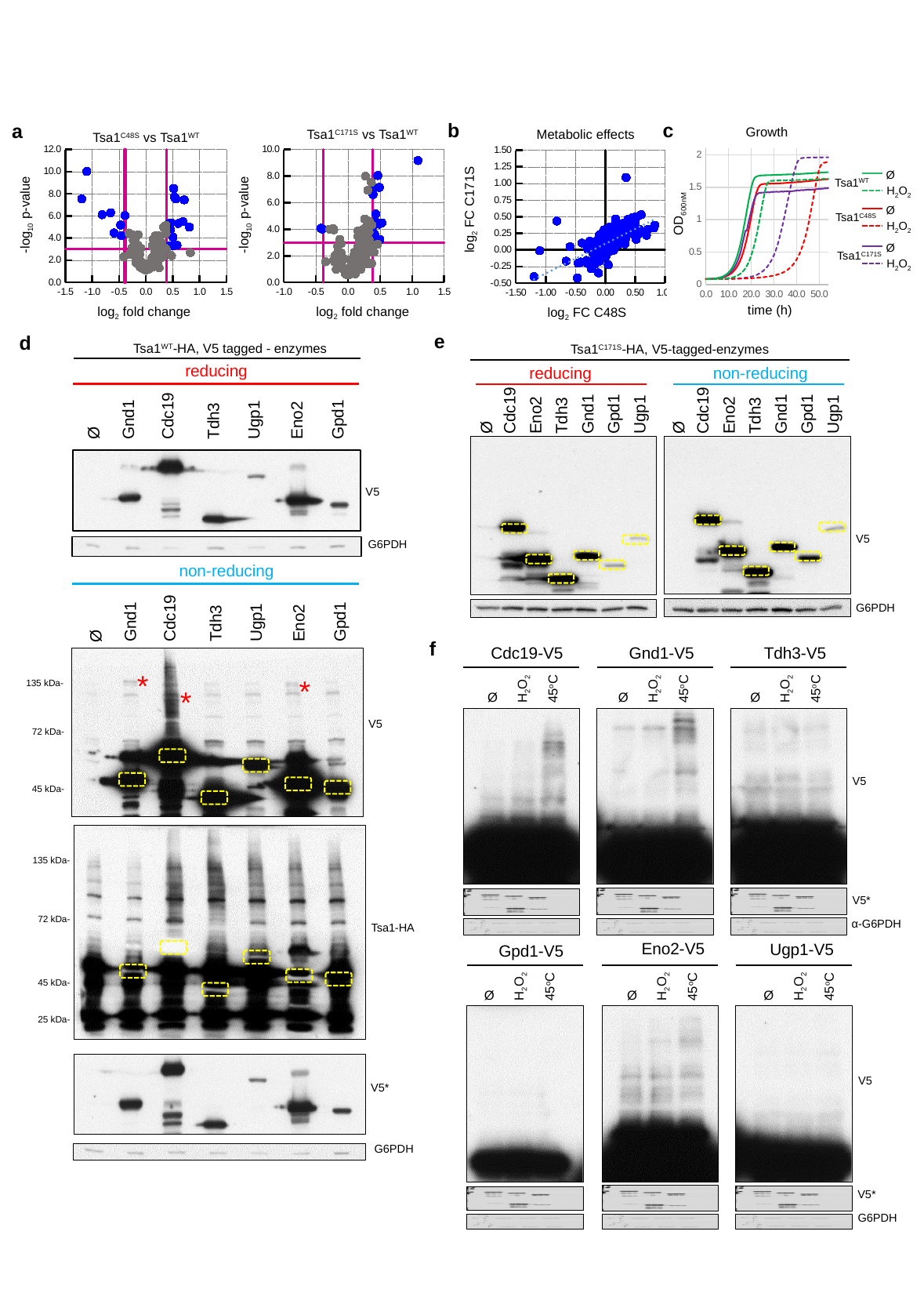

b
c
a
Growth
Metabolic effects
Tsa1^C171S^ vs Tsa1^WT^
Tsa1^C48S^ vs Tsa1^WT^
#### Chart
| Category |
|---|
#### Chart
| Category |
|---|
#### Chart
| Category |
|---|
#### Chart
| Category | | | |
|---|---|---|---|Ø
H_2_O_2_
Tsa1^WT^
Ø
H_2_O_2_
Tsa1^C48S^
Ø
H_2_O_2_
Tsa1^C171S^
-log_10_ p-value
log_2_ fold change
-log_10_ p-value
log_2_ fold change
log_2_ FC C171S
OD_600nM_
time (h)
log_2_ FC C48S
e
d
Tsa1^WT^-HA, V5 tagged - enzymes
Tsa1^C171S^-HA, V5-tagged-enzymes
reducing
reducing
non-reducing
Cdc19
Gnd1
Gpd1
Ugp1
Eno2
Tdh3
Cdc19
Gnd1
Gpd1
Ugp1
Eno2
Tdh3
Cdc19
Gnd1
Gpd1
Ugp1
Eno2
Tdh3
Ø
Ø
Ø
V5
V5
G6PDH
non-reducing
Cdc19
Gnd1
Gpd1
Ugp1
Eno2
Tdh3
Ø
G6PDH
f
Cdc19-V5
Gnd1-V5
Tdh3-V5
H_2_O_2_
45^o^C
Ø
H_2_O_2_
45^o^C
Ø
H_2_O_2_
45^o^C
Ø
*
*
135 kDa-
*
V5
72 kDa-
V5
V5*
α-G6PDH
45 kDa-
135 kDa-
72 kDa-
Tsa1-HA
Eno2-V5
Ugp1-V5
Gpd1-V5
H_2_O_2_
45^o^C
Ø
H_2_O_2_
45^o^C
Ø
H_2_O_2_
45^o^C
Ø
45 kDa-
25 kDa-
V5
V5*
G6PDH
V5*
G6PDH

### Slide 14
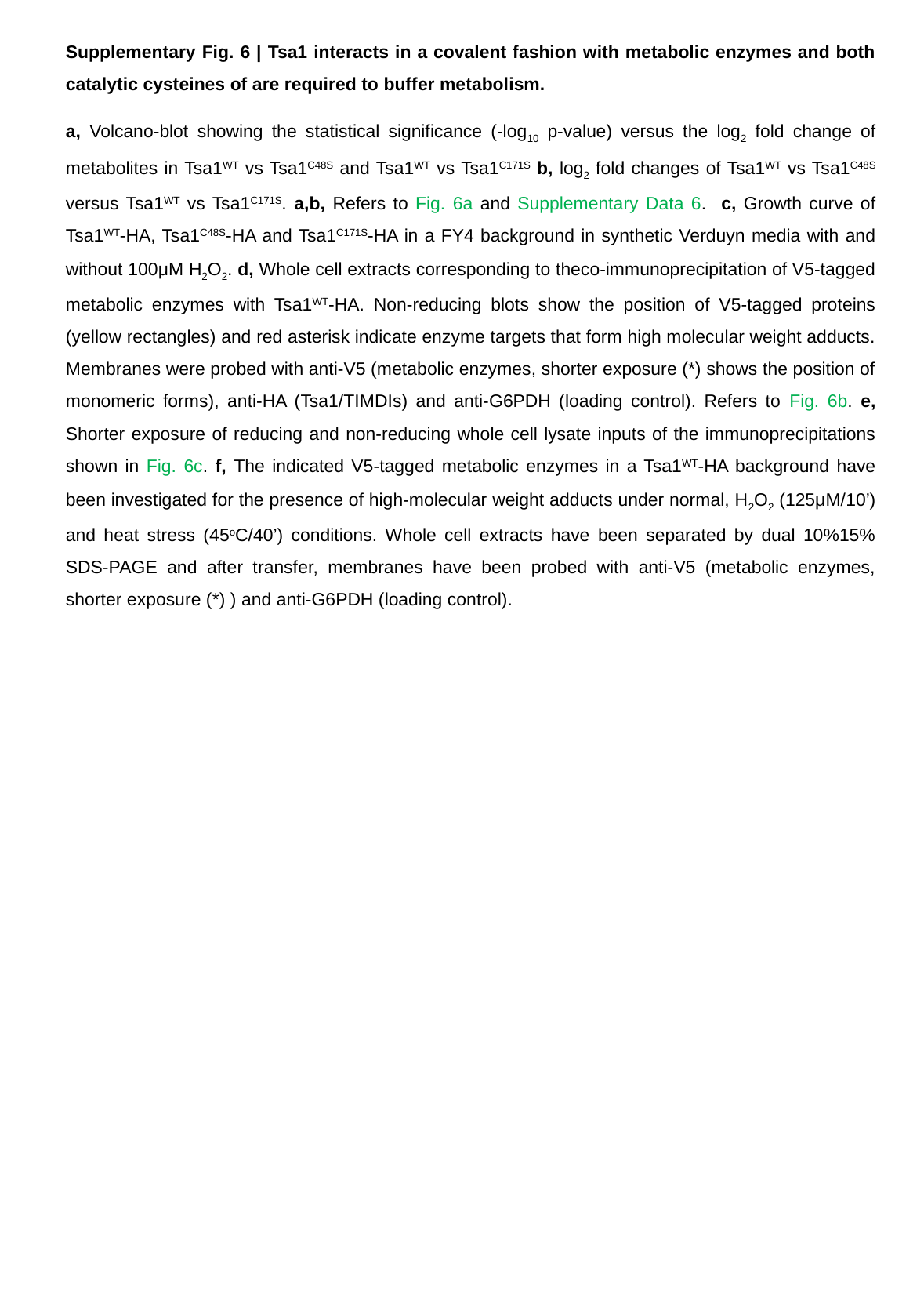

Supplementary Fig. 6 | Tsa1 interacts in a covalent fashion with metabolic enzymes and both catalytic cysteines of are required to buffer metabolism.
a, Volcano-blot showing the statistical significance (-log_10_ p-value) versus the log_2_ fold change of metabolites in Tsa1^WT^ vs Tsa1^C48S^ and Tsa1^WT^ vs Tsa1^C171S^ b, log_2_ fold changes of Tsa1^WT^ vs Tsa1^C48S^ versus Tsa1^WT^ vs Tsa1^C171S^. a,b, Refers to Fig. 6a and Supplementary Data 6. c, Growth curve of Tsa1^WT^-HA, Tsa1^C48S^-HA and Tsa1^C171S^-HA in a FY4 background in synthetic Verduyn media with and without 100μM H_2_O_2_. d, Whole cell extracts corresponding to theco-immunoprecipitation of V5-tagged metabolic enzymes with Tsa1^WT^-HA. Non-reducing blots show the position of V5-tagged proteins (yellow rectangles) and red asterisk indicate enzyme targets that form high molecular weight adducts. Membranes were probed with anti-V5 (metabolic enzymes, shorter exposure (*) shows the position of monomeric forms), anti-HA (Tsa1/TIMDIs) and anti-G6PDH (loading control). Refers to Fig. 6b. e, Shorter exposure of reducing and non-reducing whole cell lysate inputs of the immunoprecipitations shown in Fig. 6c. f, The indicated V5-tagged metabolic enzymes in a Tsa1^WT^-HA background have been investigated for the presence of high-molecular weight adducts under normal, H_2_O_2_ (125μM/10’) and heat stress (45^o^C/40’) conditions. Whole cell extracts have been separated by dual 10%15% SDS-PAGE and after transfer, membranes have been probed with anti-V5 (metabolic enzymes, shorter exposure (*) ) and anti-G6PDH (loading control).

### Slide 15
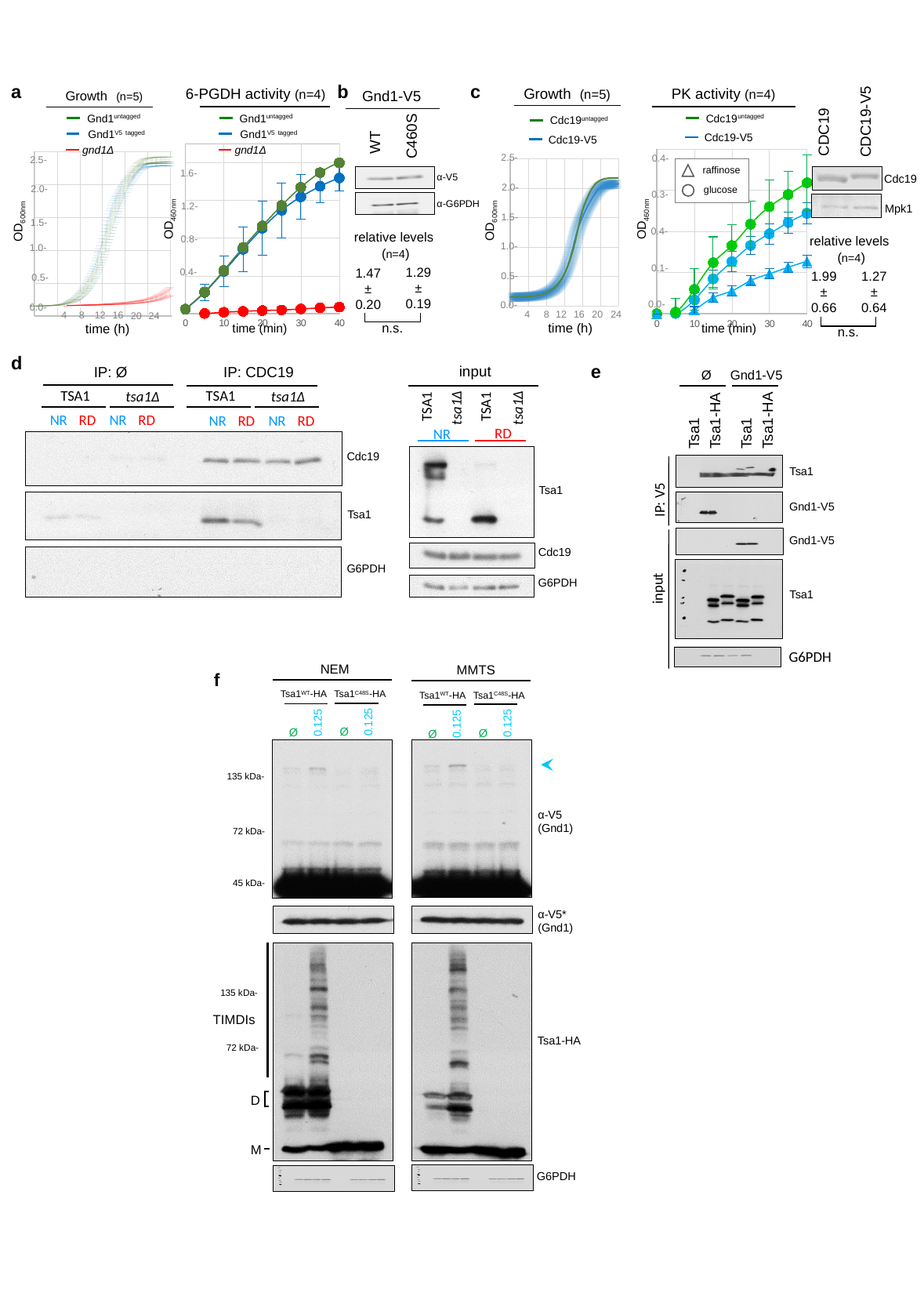

a
b
c
Growth (n=5)
6-PGDH activity (n=4)
 PK activity (n=4)
Growth (n=5)
Gnd1-V5
Cdc19^untagged^
Gnd1^untagged^
Gnd1^V5^ ^tagged^
gnd1Δ
Gnd1^untagged^
Gnd1^V5^ ^tagged^
gnd1Δ
CDC19-V5
Cdc19^untagged^
Cdc19-V5
CDC19
C460S
Cdc19-V5
WT
#### Chart
| Category |
|---|
#### Chart
| Category | 1020_average | 1026_average | Δgnd1_average |
|---|---|---|---|
#### Chart
| Category | | | |
|---|---|---|---|2.5-
0.4-
#### Chart
| Category | GND1 untag | GND1-V5 |
|---|---|---|2.5-
1.6-
1.2-
0.8-
0.4-
raffinose
α-V5
Cdc19
2.0-
2.0-
glucose
0.3-
α-G6PDH
Mpk1
1.5-
OD_460nm_
OD_460nm_
1.5-
OD_600nm_
OD_600nm_
0.4-
relative levels
relative levels
1.0-
1.0-
(n=4)
(n=4)
0.1-
1.29
±
0.19
1.47
±
0.20
1.99
±
0.66
1.27
±
0.64
0.5-
0.5-
0.0-
0.0-
0.0-
4
8
12
16
20
24
8
4
12
16
20
24
n.s.
time (h)
time (min)
time (min)
time (h)
n.s.
d
e
input
IP: CDC19
IP: Ø
Ø
Gnd1-V5
TSA1
tsa1Δ
TSA1
tsa1Δ
TSA1
TSA1
tsa1Δ
tsa1Δ
NR
RD
NR
RD
NR
RD
NR
RD
Tsa1-HA
Tsa1-HA
RD
Tsa1
Tsa1
NR
Cdc19
Tsa1
Tsa1
IP: V5
Gnd1-V5
Tsa1
Gnd1-V5
Cdc19
G6PDH
G6PDH
input
Tsa1
G6PDH
NEM
Tsa1^WT^-HA
0.125
Ø
Tsa1^C48S^-HA
0.125
Ø
MMTS
Tsa1^WT^-HA
0.125
Ø
Tsa1^C48S^-HA
0.125
Ø
α-V5
(Gnd1)
α-V5*
(Gnd1)
Tsa1-HA
G6PDH
f
135 kDa-
72 kDa-
45 kDa-
135 kDa-
TIMDIs
72 kDa-
D
M

### Slide 16
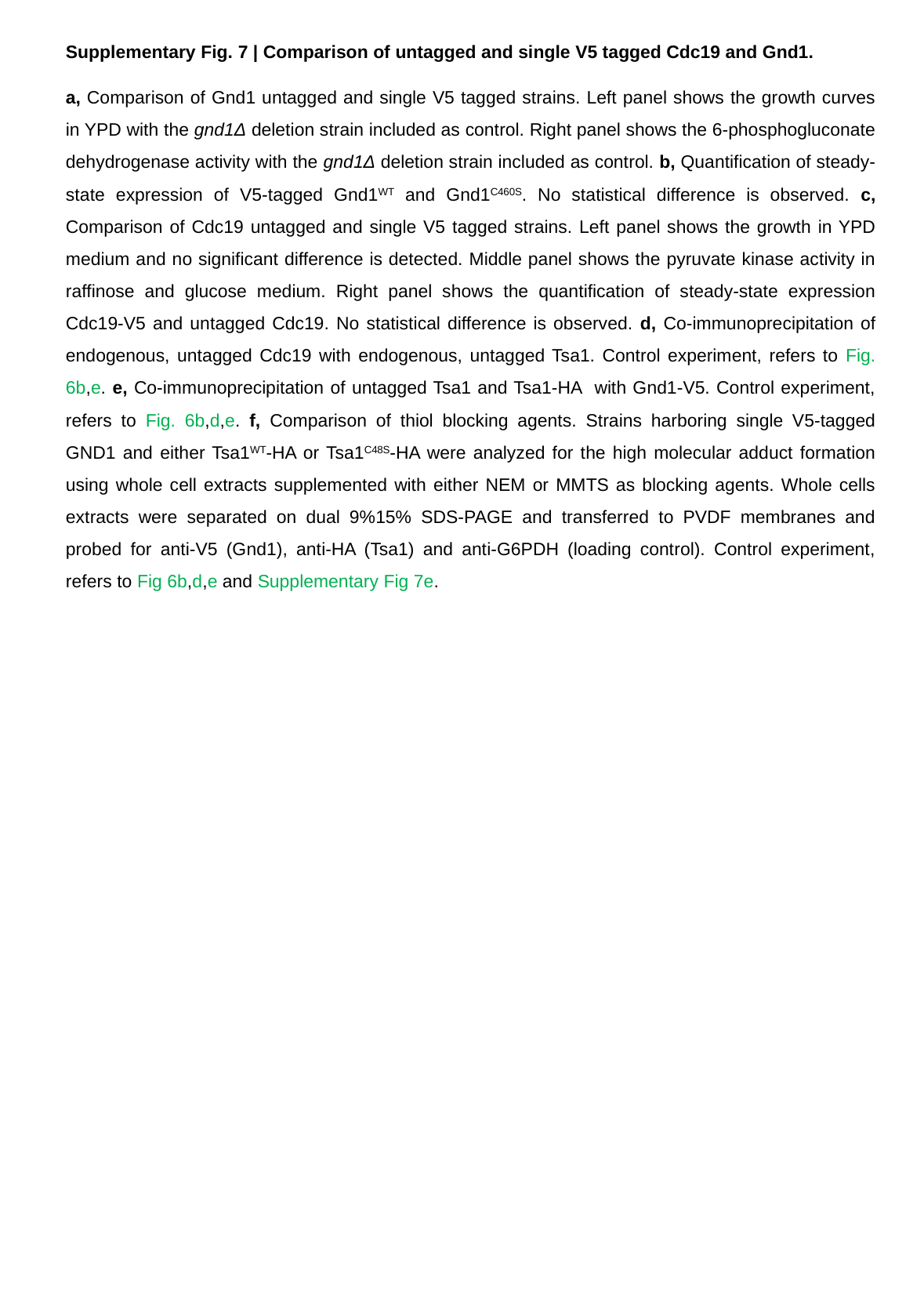

Supplementary Fig. 7 | Comparison of untagged and single V5 tagged Cdc19 and Gnd1.
a, Comparison of Gnd1 untagged and single V5 tagged strains. Left panel shows the growth curves in YPD with the gnd1Δ deletion strain included as control. Right panel shows the 6-phosphogluconate dehydrogenase activity with the gnd1Δ deletion strain included as control. b, Quantification of steady-state expression of V5-tagged Gnd1^WT^ and Gnd1^C460S^. No statistical difference is observed. c, Comparison of Cdc19 untagged and single V5 tagged strains. Left panel shows the growth in YPD medium and no significant difference is detected. Middle panel shows the pyruvate kinase activity in raffinose and glucose medium. Right panel shows the quantification of steady-state expression Cdc19-V5 and untagged Cdc19. No statistical difference is observed. d, Co-immunoprecipitation of endogenous, untagged Cdc19 with endogenous, untagged Tsa1. Control experiment, refers to Fig. 6b,e. e, Co-immunoprecipitation of untagged Tsa1 and Tsa1-HA with Gnd1-V5. Control experiment, refers to Fig. 6b,d,e. f, Comparison of thiol blocking agents. Strains harboring single V5-tagged GND1 and either Tsa1^WT^-HA or Tsa1^C48S^-HA were analyzed for the high molecular adduct formation using whole cell extracts supplemented with either NEM or MMTS as blocking agents. Whole cells extracts were separated on dual 9%15% SDS-PAGE and transferred to PVDF membranes and probed for anti-V5 (Gnd1), anti-HA (Tsa1) and anti-G6PDH (loading control). Control experiment, refers to Fig 6b,d,e and Supplementary Fig 7e.

### Slide 17
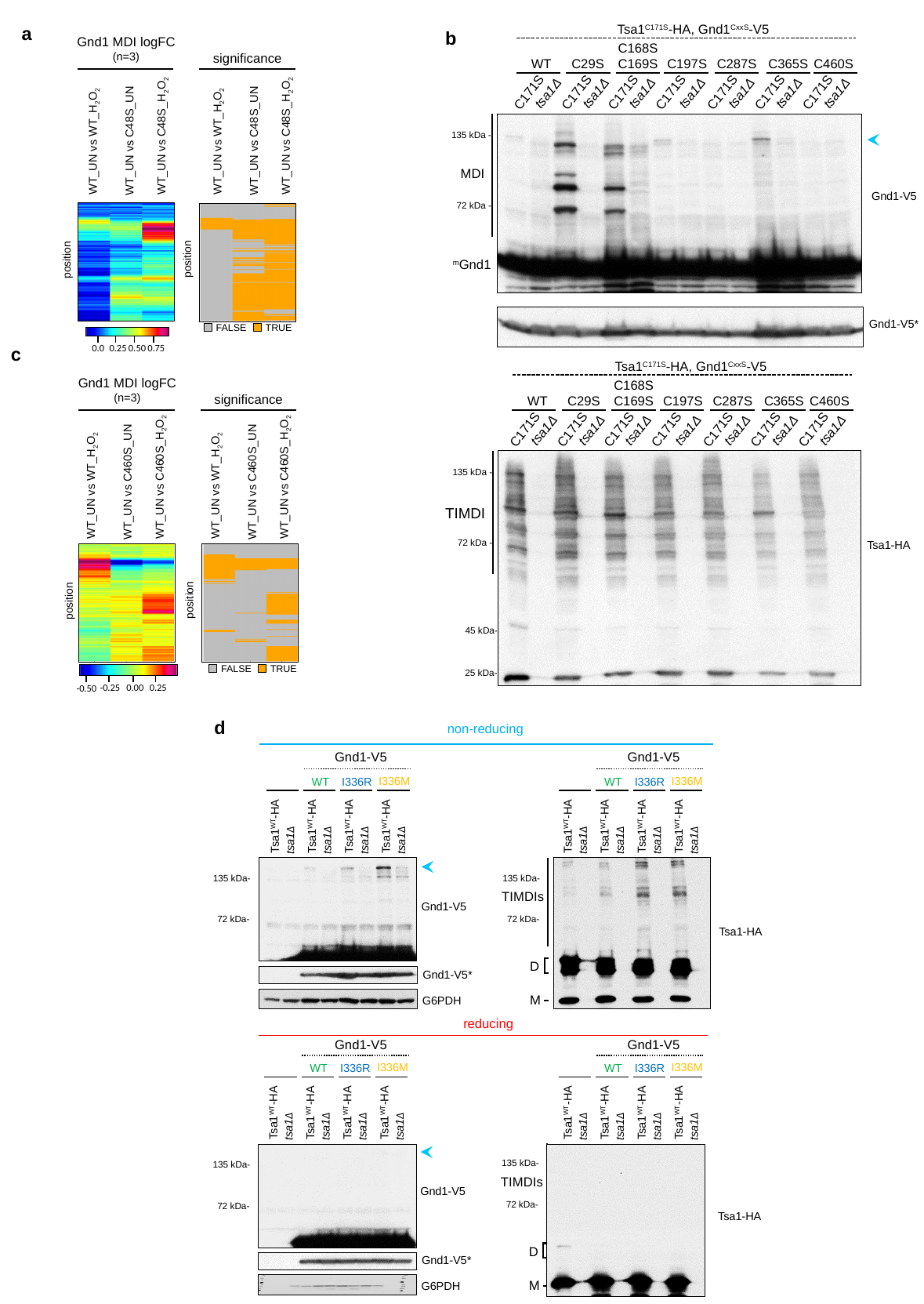

Tsa1^C171S^-HA, Gnd1^CxxS^-V5
C168S
C169S
WT
C29S
C197S
C287S
C365S
C460S
C171S
tsa1Δ
C171S
tsa1Δ
C171S
tsa1Δ
C171S
tsa1Δ
C171S
tsa1Δ
C171S
tsa1Δ
C171S
tsa1Δ
a
b
Gnd1 MDI logFC (n=3)
significance
135 kDa -
72 kDa -
WT_UN vs C48S_H_2_O_2_
WT_UN vs C48S_H_2_O_2_
WT_UN vs C48S_UN
WT_UN vs C48S_UN
WT_UN vs WT_H_2_O_2_
WT_UN vs WT_H_2_O_2_
MDI
Gnd1-V5
position
position
^m^Gnd1
Gnd1-V5*
FALSE
TRUE
0.0
0.25
0.50
0.75
c
Tsa1^C171S^-HA, Gnd1^CxxS^-V5
Gnd1 MDI logFC (n=3)
significance
C168S
C169S
WT
C29S
C197S
C287S
C365S
C460S
C171S
tsa1Δ
C171S
tsa1Δ
C171S
tsa1Δ
C171S
tsa1Δ
C171S
tsa1Δ
C171S
tsa1Δ
C171S
tsa1Δ
WT_UN vs C460S_H_2_O_2_
WT_UN vs C460S_H_2_O_2_
WT_UN vs C460S_UN
WT_UN vs C460S_UN
WT_UN vs WT_H_2_O_2_
WT_UN vs WT_H_2_O_2_
position
position
135 kDa -
72 kDa -
TIMDI
Tsa1-HA
45 kDa-
25 kDa-
FALSE
TRUE
-0.50
-0.25
0.00
0.25
d
non-reducing
Gnd1-V5
Gnd1-V5
I336M
WT
I336R
Tsa1^WT^-HA
tsa1Δ
Tsa1^WT^-HA
tsa1Δ
Tsa1^WT^-HA
tsa1Δ
Tsa1^WT^-HA
tsa1Δ
I336M
WT
I336R
Tsa1^WT^-HA
tsa1Δ
Tsa1^WT^-HA
tsa1Δ
Tsa1^WT^-HA
tsa1Δ
Tsa1^WT^-HA
tsa1Δ
135 kDa-
TIMDIs
72 kDa-
D
M
135 kDa-
Gnd1-V5
72 kDa-
Tsa1-HA
Gnd1-V5*
G6PDH
reducing
Gnd1-V5
Gnd1-V5
I336M
WT
I336R
Tsa1^WT^-HA
tsa1Δ
Tsa1^WT^-HA
tsa1Δ
Tsa1^WT^-HA
tsa1Δ
Tsa1^WT^-HA
tsa1Δ
I336M
WT
I336R
Tsa1^WT^-HA
tsa1Δ
Tsa1^WT^-HA
tsa1Δ
Tsa1^WT^-HA
tsa1Δ
Tsa1^WT^-HA
tsa1Δ
135 kDa-
TIMDIs
72 kDa-
D
M
Tsa1-HA
Gnd1-V5
Gnd1-V5*
G6PDH
135 kDa-
72 kDa-

### Slide 18
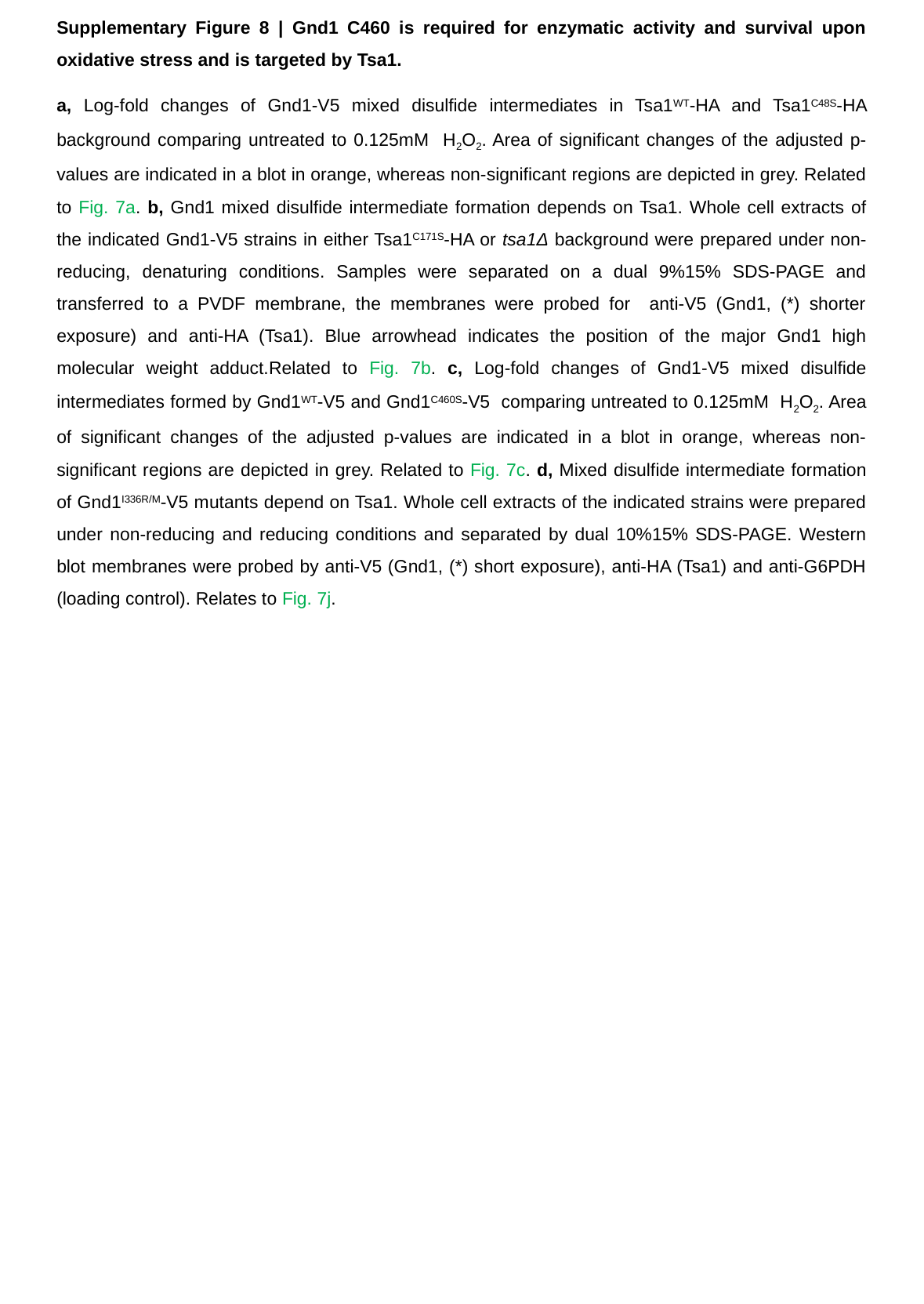

Supplementary Figure 8 | Gnd1 C460 is required for enzymatic activity and survival upon oxidative stress and is targeted by Tsa1.
a, Log-fold changes of Gnd1-V5 mixed disulfide intermediates in Tsa1^WT^-HA and Tsa1^C48S^-HA background comparing untreated to 0.125mM H_2_O_2_. Area of significant changes of the adjusted p-values are indicated in a blot in orange, whereas non-significant regions are depicted in grey. Related to Fig. 7a. b, Gnd1 mixed disulfide intermediate formation depends on Tsa1. Whole cell extracts of the indicated Gnd1-V5 strains in either Tsa1^C171S^-HA or tsa1Δ background were prepared under non-reducing, denaturing conditions. Samples were separated on a dual 9%15% SDS-PAGE and transferred to a PVDF membrane, the membranes were probed for anti-V5 (Gnd1, (*) shorter exposure) and anti-HA (Tsa1). Blue arrowhead indicates the position of the major Gnd1 high molecular weight adduct.Related to Fig. 7b. c, Log-fold changes of Gnd1-V5 mixed disulfide intermediates formed by Gnd1^WT^-V5 and Gnd1^C460S^-V5 comparing untreated to 0.125mM H_2_O_2_. Area of significant changes of the adjusted p-values are indicated in a blot in orange, whereas non-significant regions are depicted in grey. Related to Fig. 7c. d, Mixed disulfide intermediate formation of Gnd1^I336R/M^-V5 mutants depend on Tsa1. Whole cell extracts of the indicated strains were prepared under non-reducing and reducing conditions and separated by dual 10%15% SDS-PAGE. Western blot membranes were probed by anti-V5 (Gnd1, (*) short exposure), anti-HA (Tsa1) and anti-G6PDH (loading control). Relates to Fig. 7j.

### Slide 19
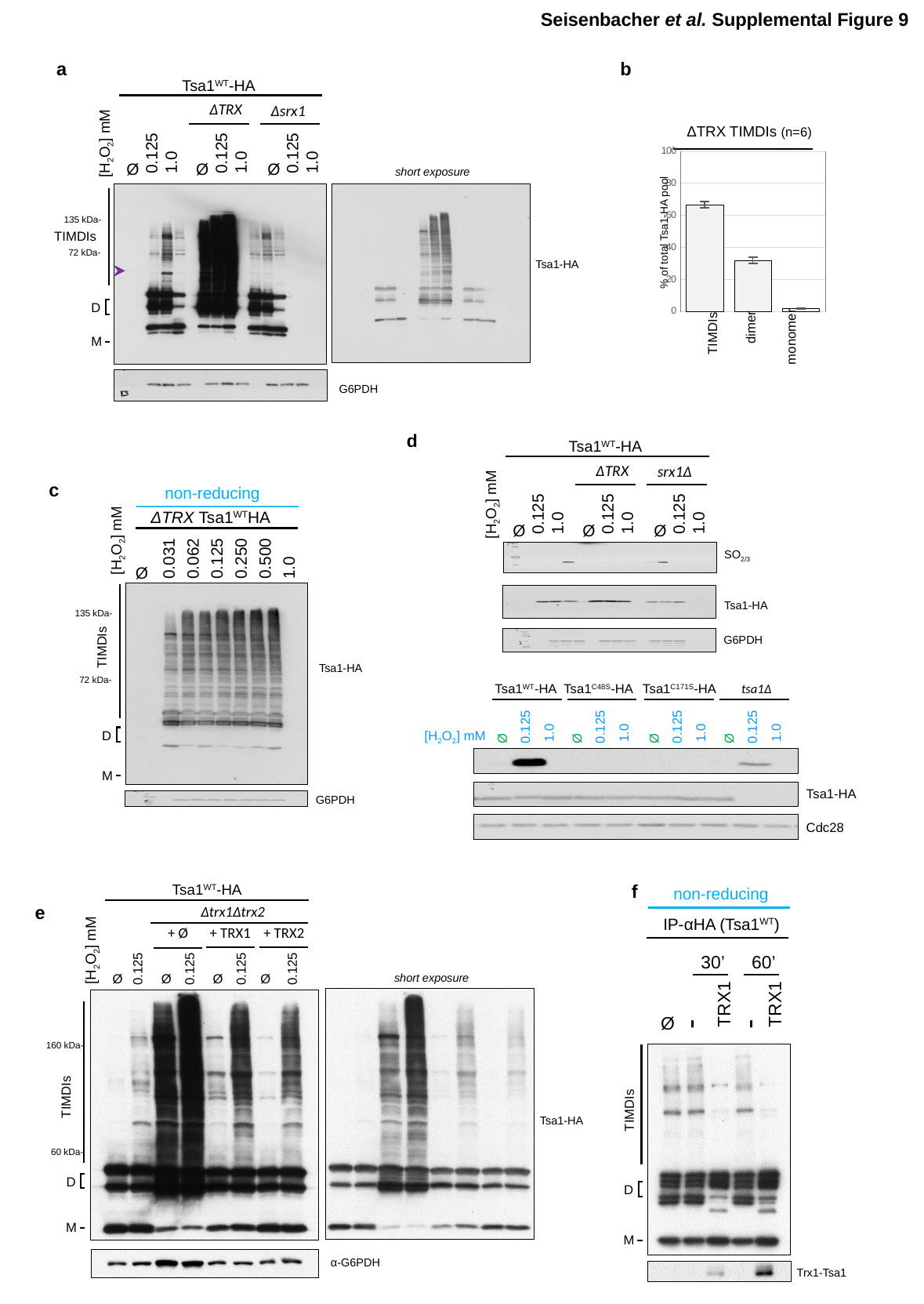

Seisenbacher et al. Supplemental Figure 9
a
b
Tsa1^WT^-HA
ΔTRX
Δsrx1
0.125
1.0
Ø
0.125
1.0
Ø
0.125
1.0
Ø
[H_2_O_2_] mM
Tsa1-HA
G6PDH
ΔTRX TIMDIs (n=6)
#### Chart
| Category | meadian |
|---|---|
| %TIMDI | 66.58242505908336 |
| %DIMER | 31.817743315931317 |
| %MONOMER | 1.675722823257615 |% of total Tsa1-HA pool
dimer
TIMDIs
monomer
short exposure
135 kDa-
TIMDIs
72 kDa-
D
M
d
Tsa1^WT^-HA
ΔTRX
srx1Δ
0.125
1.0
Ø
0.125
1.0
Ø
0.125
1.0
Ø
[H_2_O_2_] mM
c
non-reducing
ΔTRX Tsa1^WT^HA
[H_2_O_2_] mM
0.031
0.062
0.125
0.250
0.500
1.0
Ø
Tsa1-HA
G6PDH
SO_2/3_
Tsa1-HA
135 kDa-
G6PDH
TIMDIs
72 kDa-
Tsa1^WT^-HA
1.0
0.125
Ø
Tsa1^C48S^-HA
1.0
0.125
Ø
Tsa1^C171S^-HA
1.0
0.125
Ø
tsa1Δ
1.0
0.125
Ø
[H_2_O_2_] mM
Tsa1-HA
Cdc28
D
M
f
non-reducing
IP-αHA (Tsa1^WT^)
30’
60’
TRX1
TRX1
-
-
Ø
Trx1-Tsa1
Tsa1^WT^-HA
e
Δtrx1Δtrx2
+ TRX1
+ TRX2
+ Ø
0.125
Ø
0.125
Ø
0.125
Ø
0.125
Ø
[H_2_O_2_] mM
Tsa1-HA
α-G6PDH
short exposure
160 kDa-
TIMDIs
60 kDa-
D
M
TIMDIs
D
M

### Slide 20
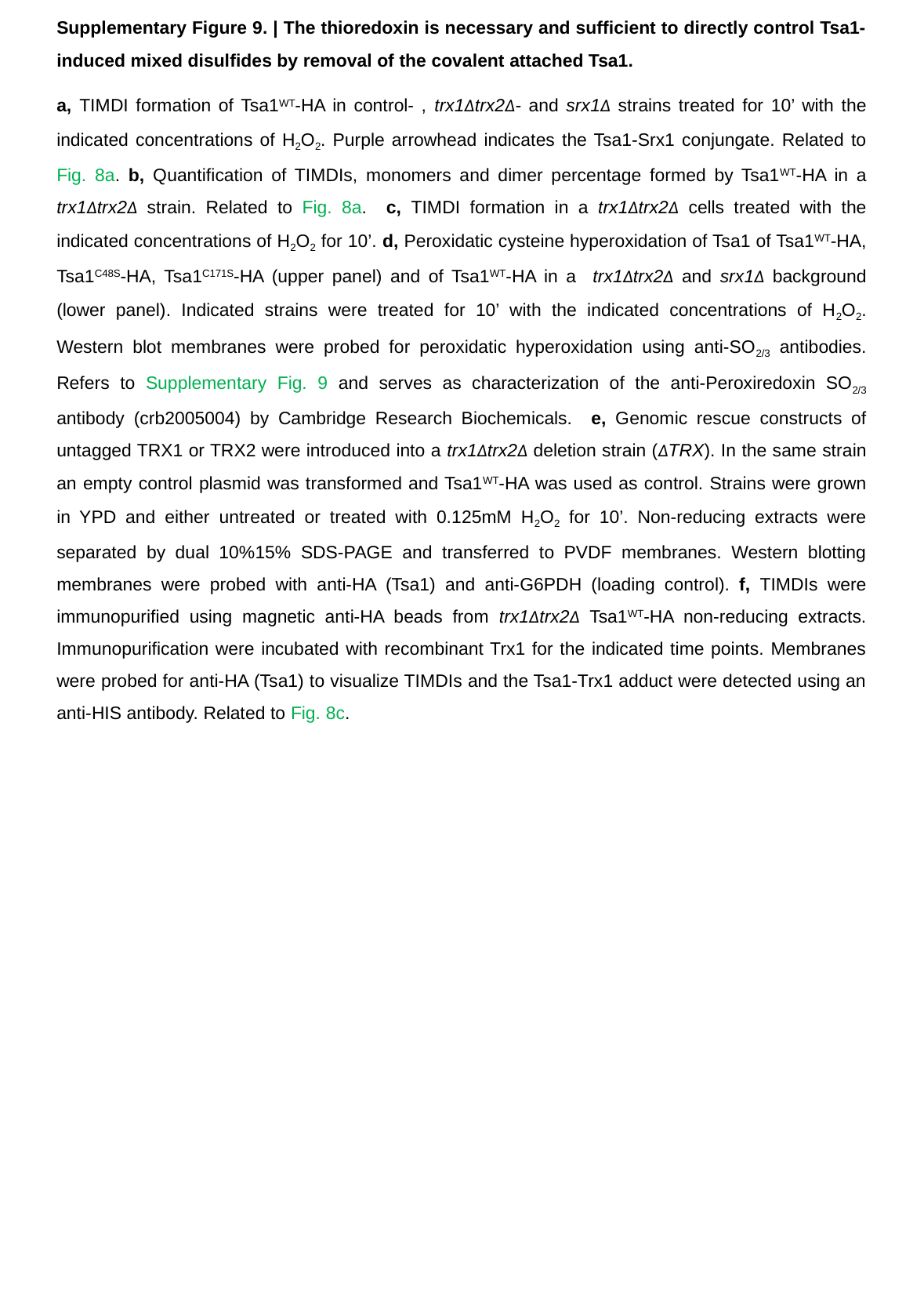

Supplementary Figure 9. | The thioredoxin is necessary and sufficient to directly control Tsa1-induced mixed disulfides by removal of the covalent attached Tsa1.
a, TIMDI formation of Tsa1^WT^-HA in control- , trx1Δtrx2Δ- and srx1Δ strains treated for 10’ with the indicated concentrations of H_2_O_2_. Purple arrowhead indicates the Tsa1-Srx1 conjungate. Related to Fig. 8a. b, Quantification of TIMDIs, monomers and dimer percentage formed by Tsa1^WT^-HA in a trx1Δtrx2Δ strain. Related to Fig. 8a. c, TIMDI formation in a trx1Δtrx2Δ cells treated with the indicated concentrations of H_2_O_2_ for 10’. d, Peroxidatic cysteine hyperoxidation of Tsa1 of Tsa1^WT^-HA, Tsa1^C48S^-HA, Tsa1^C171S^-HA (upper panel) and of Tsa1^WT^-HA in a trx1Δtrx2Δ and srx1Δ background (lower panel). Indicated strains were treated for 10’ with the indicated concentrations of H_2_O_2_. Western blot membranes were probed for peroxidatic hyperoxidation using anti-SO_2/3_ antibodies. Refers to Supplementary Fig. 9 and serves as characterization of the anti-Peroxiredoxin SO_2/3_ antibody (crb2005004) by Cambridge Research Biochemicals. e, Genomic rescue constructs of untagged TRX1 or TRX2 were introduced into a trx1Δtrx2Δ deletion strain (ΔTRX). In the same strain an empty control plasmid was transformed and Tsa1^WT^-HA was used as control. Strains were grown in YPD and either untreated or treated with 0.125mM H_2_O_2_ for 10’. Non-reducing extracts were separated by dual 10%15% SDS-PAGE and transferred to PVDF membranes. Western blotting membranes were probed with anti-HA (Tsa1) and anti-G6PDH (loading control). f, TIMDIs were immunopurified using magnetic anti-HA beads from trx1Δtrx2Δ Tsa1^WT^-HA non-reducing extracts. Immunopurification were incubated with recombinant Trx1 for the indicated time points. Membranes were probed for anti-HA (Tsa1) to visualize TIMDIs and the Tsa1-Trx1 adduct were detected using an anti-HIS antibody. Related to Fig. 8c.
