## Supplementary Data 6 - Sequence of ssDNA oligos used in this study for "Peroxiredoxinylation buffers the redox state of the proteome upon cellular stress"

| Name | Sequence | Purpose |
|---|---|---|
| TSA1aux f01 | CTATCTATATATAAAACCCATAAACTTAATGCTATTCACTGACGAATTCCCGGGGATCCGG | insertion of auxotrophic maker 3' downstream of Tsa1 CDS |
| TSA1aux r01 | CAGACGATGATTGTTATTAGAACTGAGCTAGTGTGAATAGCCTGCAGGTCGACGGATCC | insertion of auxotrophic maker 3' downstream of Tsa1 CDS |
| TSA1 mHA f01 | CGTTGAAGACTCCAAGGAATACTTCGAAGCTGCCAACAAAtatccatacgatgttcctgattatgctggttgaGACGCTTGCAGAGTTGTCTAAATGAC | C-terminal single HA-tagging of TSA1 |
| Ctag TSA1 r03 | AGGGCAGCAACAGAGCTAAAGTGAGTGTGGCAGTG | C-terminal single HA-tagging of TSA1 |
| C48S f02 | TACGTTGTCCTAGCCTTTATTCCATTGGCCTTCACTTTCGTCaGTCCAACCGAAATCATTGC | Knock-IN mutant for Tsa1 peroxidatic C48 cysteine |
| C171S wt f01 | GACGAAGCCTTGAGATTGGTTGAAGCCTTCCAATGGACCGACAAGAACGGTACTGTCTTGCCAaGTAACTGGACTCCAGGTGCTGC | Knock-IN mutant for Tsa1 resolving C171 cysteine |
| Gex Tsa1 BamhI f01 | atgGGATCCGTCGCTCAAGTTCAAAAGCAAGCTCCAAC | For CDS Tsa1 cloning into pGEX |
| Gex Tsa1 HA Noti r01 | atgGCGGCCGCtcaaccagcataatcaggaacatcgtatgg | For CDS Tsa1 cloning into pGEX |
| etm trx f01 | ataCCATGGTTACTCAATTCAAAAC | For CDS Trx1 cloning into pETM series |
| etm trx r01 | ataGCGGCCGCTTAAGCATTAGCAGC | For CDS Trx1 cloning into pETM series |
| CDC19-V5-tbx-f01 | CACTCCAACACTTTGCAAGTCTCTACCGTTggtaagcctatccctaaccctctcctcggtctcgattctacgTAAaagtggaacgatcattcaagagatc | single V5 tagging of CDC19 |
| CDC19 tbx r01 | TTTAAAAAATATAATTCAAAAAAATAATATCTTCATTCAATCATGATTCTTTTTATCGATGAATTCGAGCTCG | single V5 tagging of CDC19 |
| TDH3-V5-tbx-f01 | GTCGACTTGGTTGAACACGTTGCCAAGGCTggtaagcctatccctaaccctctcctcggtctcgattctacgTAAaagtggaacgatcattcaagagatc | single V5 tagging of TDH3 |
| TDH3 tbx r01 | ATAAAGCTATAAAAAGAAAATTTATTTAAATGCAAGATTTAAAGTAAATTCACATCGATGAATTCGAGCTCG | single V5 tagging of TDH3 |
| GND1-V5-tbx-f01 | GGTAATGTTTCTTCCTCTACATACCAAGCTggtaagcctatccctaaccctctcctcggtctcgattctacgTAAaagtggaacgatcattcaagagatc | single V5 tagging of GND1 |
| GND1 tbx ro1 | TACTATAACTAATCTATTTCATTAAATTTTTTTGGTTTATGTCATCGATGAATTCGAGCTCG | single V5 tagging of GND1 |
| GPD1-V5-tbx f01 | ATGATTGAAGAATTAGATCTACATGAAGATggtaagcctatccctaaccctctcctcggtctcgattctacgTAAaagtggaacgatcattcaagagatc | single V5 tagging of GPD1 |
| GPD1-tbx r01 | GAAAAAAGTGGGGGAAAGTATGATATGTTATCTTTCTCCAATAAATATCGATGAATTCGAGCTCG | single V5 tagging of GPD1 |
| UGP1-V5-tbxf01 | GTTACTGGTAATTTGCAAATCTTGGAACATggtaagcctatccctaaccctctcctcggtctcgattctacgTAAaagtggaacgatcattcaagagatc | single V5 tagging of UGP1 |
| UGP1-tbx-r01 | AATAACCTAACGGAGAATTGAAAGTAAAAAAGGGATTGCAAGCCGCACCTATCGATGAATTCGAGCTCG | single V5 tagging of UGP1 |
| eno2V5tbx f01 | [illegible]ACAAGGCTGTCTACGCCGGTGAAAACTTCCACCACGGTGACAAGTTGggtaagcctatccctaaccctctcctcggtctcgattctacgTAACGTACGCTGCAGGTCGAC | single V5 tagging of ENO2 |
| eno2 tbx r01 | AACTATGATGAAAAAATAAGCAGAAAAGACTAATAATTCTTAGTTAAAAGCACTATCGATGAATTCGAGCTCG | single V5 tagging of ENO2 |
| Gnd1 C01 ydp f01 | GCCGTCATGGGTCAAAATTTGATCTTGAACGCTGCTGACCACGGGAATTCCCGGGGATCCGG | deletion of Gnd1 region C29 |
| Gnd1 C01 ydp r01 | TGCCCTTAGCTTCATTGGCCAAGAAATGGTCGACCTTGGATTGAGTTCTGCAGGTCGACGGATCC | deletion of Gnd1 region C29 |
| Gnd1 CS01 f01 | ATGGGTCAAAATTTGATCTTGAACGCTGCTGACCACGGTTTCACTGTTTcTGCTTACAACAGAACTCAATC | Knock-IN mutant for GND1 cysteine C29S |
| Gnd1 C0203 ydp f01 | TGAAGAAGCTTGGCCACATATTAAGAACATCTTCCAATCCATCTCTGCTAAGAATTCCCGGGGATCCGG | deletion of Gnd1 region C168/169 |
| Gnd1 C0203 ydp r01 | ATCACCGTATTCAATACCGTTGTGAACCATCTTGACGTAGTGACCAGCACCCTGCAGGTCGACGGATCC | deletion of Gnd1 region C168/169 |
| Gnd1 CS0203 f01 | AGAAGCTTGGCCACATATTAAGAACATCTTCCAATCCATCTCTGCTAAATCCGACGGTGAACCATcTTcCGAATGGGTTGGCCCAGCC | Knock-IN mutant for GND1 cysteine C168/169S |
| Gnd1 C04 ydp f01 | TGCTGGTCACTACGTCAAGATGGTTCACAACGGTATTGAATACGGTGATGAATTCCCGGGGATCCGG | deletion of Gnd1 region C197 |
| Gnd1 C04 ydp r01 | CAAAAACGTCACTGATTTCCTTATCGGTAAACCCACCCAATCTCTTCATGACTGCAGGTCGACGGATCC | deletion of Gnd1 region C197 |
| Gnd1 CS04 f01 | TCACTACGTCAAGATGGTTCACAACGGTATTGAATACGGTGATATGCAATTGATTTcTGAAGCTTATGACATCATGAAG | Knock-IN mutant for GND1 cysteine C197S |
| Gnd1 C05 ydp f01 | TGGACTGCCATCAACGCCTTGGATTTGGGTATGCCAGTTACTTTGATTGGTGGAATTCCCGGGGATCCGG | deletion of Gnd1 region C287 |
| Gnd1 C05 ydp r01 | CGTCTTTTGGAACTTCTGGGCCTGGTAAGACCTTGGAGGCTCTAATTCTCTCGCTGCAGGTCGACGGATCC | deletion of Gnd1 region C287 |
| Gnd1 CS05 f01 | GTGGACTGCCATCAACGCCTTGGATTTGGGTATGCCAGTTACTTTGATTGGTGAAGCTGTCTTTGCCCGTTcTCTATCTGCTTTGAAGAACG | Knock-IN mutant for GND1 cysteine C287S |
| Gnd1 C06 ydp f01 | TGCTACTTATGGCTGGAAACTAAACAACCCTGCCATCGCTTTGATGTGGGAATTCCCGGGGATCCGG | deletion of Gnd1 region C365 |
| Gnd1 C06 ydp r01 | AAGTTTTCCAAATCTGGTTCTTCTCTGTAGGCCTTTGTGATTTGACCCAAGCTGCAGGTCGACGGATCC | deletion of Gnd1 region C365 |
| Gnd1 CS06 f01 | TGCTGCTACTTATGGCTGGAAACTAAACAACCCTGCCATCGCTTTGATGTGGAGAGGTGGTTcTATCATTAGATCTGTTTTCTTG | Knock-IN mutant for GND1 cysteine C365S |
| Gnd1 C07 ydp f01 | CAGCCAACTTACTACAAGCTCAACGTGACTACTTTGGTGCTCACACTTTCAGAGTGAATTCCCGGGGATCCGG | deletion of Gnd1 region C460 |
| Gnd1 C07 ydp r01 | CTTGCCAGTAGACAAGGATATCCATATCAACTGGACTGGCCACGGTGGTACTGCAGGTCGACGGATCC | deletion of Gnd1 region C460 |
| Gnd1 CS07 f01 | CCAGCCAACTTACTACAAGCTCAACGTGACTACTTTGGTGCTCACACTTTCAGAGTGTTGCCAGAATcTGCTTCTGACAACTTGCCAGTAGAC | Knock-IN mutant for GND1 cysteine C460S |
| Gnd1 check f01 | GATACTGCTGGTCAAAAGGGTACTGGTAAGTGG | sequencing Gnd1 genomic region |
| Gnd1 check r01 | CCACTTACCAGTACCCTTTTGACCAGCAGTATC | sequencing Gnd1 genomic region |
| gnd1 genomic r01 | TTAGAAAACACATAACTTCATATACTATTTATAATTGC | sequencing Gnd1 genomic region |
| gGND1 f01 | ATgcggccgcGTTTATTCCTTCCCATCAACGTCAACGATGTC | sequencing Gnd1 genomic region |
| I366M | TATGGCTGGAAACTAAACAACCCTGCCATCGCTTTGATGTGGAGAGGTGGTTGTATgATTAGATCTGTTTTCTTGGGTC | Knock-In mutant for Gnd1 I266M |
| I366R | TATGGCTGGAAACTAAACAACCCTGCCATCGCTTTGATGTGGAGAGGTGGTTGTcgCATTAGATCTGTTTTCTTGGGTC | Knock-In mutant for Gnd1 I266R |
| gTRX1 f01 | ATActcgagTCGTTGGAAAAGATGTC | cloning of genomic TRX1 |
| gTRX1 r01 | ATAGCGGCCGCACCTCCTTTGTGTGGTGGGC | cloning of genomic TRX1 |
| gTRX2 f01 | ATActcgagTTTCCAGCCAGCCGAAAGAG | cloning of genomic TRX2 |
| gTRX2 r01 | ATAGCGGCCGCACCAAGAGCATGACTGAC | cloning of genomic TRX2 |
| TDH3 Naux f01 | TACACAGAATATATAACATCGTAGGTGTCTGGGTGAACAGTTTATTCGAATTCCCGGGGATCCGG | insertion of auxotrophic maker 5' upstream of Tdh3 CDS |
| TDH3 Naux r01 | GATGCCAGCTTAAAAAGCGGGCTCCATTATATTTAGTGGATGCCTGCAGGTCGACGGATCC | insertion of auxotrophic maker 5' upstream of Tdh3 CDS |
| pTDH3-Tsa1 f01 | ATGCTGTTCTGGCCCGTCGGGTTTTCTGACAAATTGTCCTTTAGGGAGGTGTCTGGGTGAACAGTTTATTC | exchanging pTSA1 for pTDH3 promoter |
| pTDH3-Tsa1 r01 | AGTTTTCTTAAAAGTTGGAGCTTGCTTTTGAACTTGAGCGACCATTTTGTTTGTTTATGTGTGTTTATTCGAAACTAAG | exchanging pTSA1 for pTDH3 promoter |
| Rsp5auxf01 | TTAATGTTTATTTGTTTAGAAAACAGACTGCTGGTTTGTTTACATTTAATTCCCGGTGATCCGGTGATTGATTGAGC | insertion of auxotrophic maker 5' upstream of Rsp5 CDS |
| Rsp5auxr01 | ACGTGACTGTGCGGCTGGTCTGCGCCAGCAATTTTTTTTTTCTGCTCTCTGCAGGTCGACTGATCCGGTGATTGATTG | insertion of auxotrophic maker 5' upstream ofRsp5 CDS |
| pTSA1-rsp5-NAUX f01 | ATGCTGTTCTGGCCCGTCGGGTTTTCTGACAAATTGTCCTTTAGGGACAGACTGCTGGTTTGTTTAC | exchanging pTSA1 for pRsp5 promoter |
| pTSA1-rsp5-NAUX r01 | AGTTTTCTTAAAAGTTGGAGCTTGCTTTTGAACTTGAGCGACCATTGTTACTTTAAATATAATACC | exchanging pTSA1 for pRsp5 promoter |
| TDH3 single HA tbx f01 | GTCGACTTGGTTGAACACGTTGCCAAGGCTtatccatacgatgttcctgattatgctggttgacgggttaattaaggcgc | for single HA tagging of Tdh3 (C-terminus) |
| TDH3 tbx r01 | ATAAAGCTATAAAAAGAAAATTTATTTAAATGCAAGATTTAAAGTAAATTCACATCGATGAATTCGAGCTCG | for single HA tagging of Tdh3 (C-terminus) |
