## Supplementary Data 7 - S cerevisiae strains used in this study for "Peroxiredoxinylation buffers the redox state of the proteome upon cellular stress"

### *S. cerevisiae* strains generated in this study

| Strain Name | Strain genotype |
|---|---|
| BY4741 | *MATa his3Δ1 leu2Δ0 met15Δ0 ura3Δ0* |
| FY4 | *MATa* |
| YGS338 | BY4741 URA::pRSP5 |
| YGS347 | BY4741 Tsa1 3' LEU |
| YGS379 | *TSA1-GFP::HIS3* |
| YGS447 | BY4741 *TSA1-1xHA::LEU* |
| YGS448 | BY4741 *TSA1*$^{C171S}$*-1xHA::LEU* |
| YGS449 | BY4741 *TSA1*$^{C48S}$*-1xHA::LEU* |
| YGS456 | BY4741 *TSA1-TAP::KAN* |
| YGS457 | BY4741 *TSA1*$^{C48S}$*-TAP::KAN* |
| YGS458 | BY4741 *TSA1*$^{C171S}$*-TAP::KAN* |
| YGS471 | FY4 Tsa1-1xHA::KanMX6 |
| YGS472 | FY4 Tsa1$^{C48S}$-1xHA::KanMX6 |
| YGS473 | FY4 Tsa1$^{C171S}$-1xHA::KanMX6 |
| YGS836 | *HSP104-GFP::HIS3;; TSA1-1xHA::LEU* |
| YGS1026 | YGS447 *GND1-1xV5:: HphNT1* |
| YGS1028 | YGS448 *GND1-1xV5:: HphNT1* |
| YGS1030 | YGS449 *GND1-1xV5:: HphNT1* |
| YGS1053 | YGS447 *hsp104::HIS3* |
| YGS1059 | YGS448 *CDC19-1xV5:: HphNT1* |
| YGS1203 | YGS447 *srx1::NatMX4* |
| YGS1257 | YGS448 *UGP2-1xV5:: HphNT1* |
| YGS1267 | YGS448 *ENO2-1xV5:: HphNT1* |
| YSG1271 | YGS448 *GPD1-1xV5:: HphNT1* |
| YGS1275 | YGS447 *ssa1::HphNT1 ssa2::NatMX4* |
| YGS1313 | YGS447 *trx1::KanMX6 trx2::NatMX4* |
| YGS1316 | YGS1026 *trx1::KanMX6 trx2::NatMX4* |
| YGS1379 | YGS447 *GND1*$^{C460S}$*-1xV5::HPH trx1::KanMX6 trx2::NatMX4* |
| YGS1541 | YGS447 *GND1*$^{C460S}$*-1xV5::HphNT1* |
| YGS1558 | YGS448 *TDH3-1xV5:: HphNT1* |
| YGS1551 | YGS448 *GND1*$^{C29S}$*-1xV5:: HphNT1* |
| YGS1552 | YGS448 *GND1*$^{C168/169S}$*-1xV5:: HphNT1* |
| YGS1553 | YGS448 *GND1*$^{C197S}$*-1xV5:: HphNT1* |
| YGS1554 | YGS448 *GND1*$^{C287S}$*-1xV5:: HphNT1* |
| YGS1555 | YGS448 *GND1*$^{C365S}$*-1xV5:: HphNT1* |
| YGS1557 | YGS448 *GND1*$^{C460S}$*-1xV5:: HphNT1* |
| YGS1573 | YGS447 URA::pRsp5>Tsa1 |
| YGS1733 | BY4741 TDH3-1xHA::*HpHNT1* |
| YGS1734 | YGS447 URA::pTDH3>Tsa1 |
| YGS1735 | BY4741 URA::pTDH3 |
