## Supplementary Data 8 - Materials and Reagents used in this study for "Peroxiredoxinylation buffers the redox state of the proteome upon cellular stress"

| Name | Source | Identifier |
|---|---|---|
| **PCR** | | |
| ssDNA oligos | Sigma-Aldrich | Document S7 |
| Expand High Fidelity Polymerase | Roche | 11759078001 |
| dNTPs (set of dATP/dCTP/dTP/dGTp) | Promega | U1420 |
| **Plasmids** | | |
| pGEX-6P-1 | Sigma-Aldrich | GE28-9546-48 |
| pYM16 | EUROSCARF | P30302 |
| pRS413 | Stratagene | |
| pGS558 | pRS413-genomic TRX1 | this study |
| pGS559 | pRS413-genomic TRX2 | this study |
| **Restriction Enzymes** | | |
| BamHI-HF | NEB | R3136S |
| XhoI | NEB | R0146S |
| NotI-HF | NEB | R3189S |
| NcoI-HF | NEB | R3193S |
| **Protein Biochemistry** | | |
| IPTG | Roche | 11411446001 |
| Glutathione Sepharose | Roche | 17-0756-01 |
| L-Glutathione | Sigma | G4251 |
| Centrifugal Filter Columns | Amicon | UFC901008 |
| PreScission Protease | GE Healthcare | 27-0843-01 |
| cOmplete™ His-Tag Purification Resin | Merck | 5893682001 |
| N-ethylmaleimide | Sigma-Aldrich | E3876 |
| S-methyl methanethiosulfonate | Merck | 64306 |
| Iodoacetamide | Sigma | I1149 |
| 1,4-dithiothreitol | Sigma | D0632 |
| rabbit IgG-Agarose | Sigma | A2909 |
| V5-Trap™ Magnetic Agarose | ,Chromotek | v5tma |
| HA-magnetic beads Pierce™ | ThermoFisher | 88836 |
| Cycloheximide solution | Sigma-Aldrich | C4859-1ML |
| L-azetidine-2-carboxylic acid | Sigma-Aldrich | A0760 |
| endoproteinase LysC | Wako | 129-02541 |
| trypsin | Promega | V5113 |
| digested bovine serum albumin | NEB | P8108S |
| **Growth Media** | | |
| YDP broth | Formedium | CCM0210 |
| LB broth base | Invitrogen | 12780052 |
| Yeast Nitrogen Base w/o amino acids | Invitrogen | Q300-09 |
| CSM-URA | MP Biomedicals | 4511222 |
| agar-agar | Merck | 01916-500G |
| **Antibodies** | | |
| α-HA antibody, ( clone 12CA5) | ROCHE | 11583816001 |
| α-V5 antibody | Invitrogen | R960-25 |
| α-Mpk1 (clone E-8) | Santa Cruz Biotechnology | Sc-374440 |
| α-G6PDH | Sigma Aldrich | A-9521 |
| α-Peroxiredoxin-SO3 antibody | Abcam | ab16830 |
| α-peroxiredoxin-SO2/3 antibody | Cambridge Research Chemicals | crb2005004d/T |
| α-mouse IgG HRP-linked whole antibody | GE Healthcare | NA931 |
| α-rabbit IgG whole antibody HRP | GE Healthcare | NA934 |
| IRDye® 680RD donkey α-rabbit | LI-COR | 925-68073 |
| IRDye® 800CW donkey α-mouse | LI-COR | 926-32212 |
| **Western Blot Analysis** | | |
| Clarity Western ECL substrate | BioRad | 1705060 |
| Intercept® blocking buffer | LI-COR | 927-60001 |
| Methanol, pure Ph Eur NF | Scharlab | ME0301005P |
| Tris Base | Sigma | T1378 |
| Glycine | VWR Chemicals | 24403.367 |
| PVDF membrane Immobilon®-P, 0,45 µm | Millipore | IPVH00010 |
| PVDF membrane Immobilon®-FL, 0.45 µm | Millipore | IPFL85R |
| medical X-ray films (18x24 cm) | AGFA | CPBU-NEW |
| pre-stained molecular weight marker | Sigma | SDS7B2-5X1VL |
| Precision Plus Protein Dual Color marker | BioRad | 161-0394SP |
| 30% Acrylamide:Bisacrylamide (37.5:1) | Panreac | A3626.0500 |
| **Microscopy** | | |
| Concanavalin A | Calbiochem | 234567 |

| | | |
|---|---|---|
| µ-Slide 8 well | IBIDI | 80826 |
| Nunc™ MicroWell™ 96-Well Microplates | ThermoFisher | 167008 |
